# Two new *Rhizobiales* species isolated from root nodules of common sainfoin (*Onobrychis viciifolia*) show different plant colonization strategies

**DOI:** 10.1101/2022.03.04.482989

**Authors:** Samad Ashrafi, Nemanja Kuzmanović, Sascha Patz, Ulrike Lohwasser, Boyke Bunk, Cathrin Spröer, Maria Lorenz, Anja Frühling, Meina Neumann-Schaal, Susanne Verbarg, Matthias Becker, Torsten Thünen

## Abstract

Root nodules of legume plants are primarily inhabited by rhizobial nitrogen-fixing bacteria. Here we propose two new *Rhizobiales* species isolated from root nodules of common sainfoin (*Onobrychis viciifolia*), as shown by core-gene phylogeny, overall genome relatedness indices and pan-genome analysis.

*Mesorhizobium onobrychidis* sp. nov., actively induces nodules, and achieves atmospheric nitrogen and carbon dioxide fixation. This species appears to be depleted in motility genes, and is enriched in genes for direct effects on plant growth performance. Its genome reveals functional and plant growth-promoting signatures like a large unique chromosomal genomic island with high density of symbiotic genetic traits. O*nobrychidicola muellerharveyae* gen. nov. sp. nov., is described as type species of the new genus O*nobrychidicola* in *Rhizobiaceae*. This species comprises unique genetic features and plant growth-promoting traits (PGPTs), which strongly indicate its function in biotic stress reduction and motility. We applied a newly developed bioinformatics approach for *in silico* prediction of PGPTs (PGPT-Pred), which supports the different lifestyles of the two new species and the plant growth-promoting performance of *M. onobrychidis* in the greenhouse trial.

## Introduction

Rhizobia is a common term referring to a paraphyletic group of bacteria, which are able to induce nodules on roots of legumes and to fix atmospheric nitrogen (N_2_). They have been investigated since the identification of their roles in nitrogen acquisition for legume plants [1, 2]. Rhizobia show variability in their nodulation strategies. Some of them are host-specific, while others can nodulate various plant species, even members of non-legume plants [3]. Rhizobia comprise genetically diverse group of bacteria. They share a symbiotic nitrogen fixation function that is encoded on symbiotic plasmids or symbiosis islands within the genome [4, 5], jointly termed as symbiotic genome compartments (SGCs) [6]. Legume root nodules are principally inhabited by nitrogen-fixing bacteria. However, this ecological niche contains many other non-rhizobial bacterial species, collectively called nodule-associated bacteria [7, 8]. They are involved in different biological activities *e*.*g*. plant growth-promotion and biocontrol [9]. Nevertheless, our knowledge about the entire biological functions of nodule-associated bacteria is elusive.

Based on the current taxonomical information, rhizobia are classified within a number of families of the alphaproteobacterial order *Rhizobiales*. Non-nitrogen-fixing *Rhizobiaceae* members were also recovered from legume root nodules [10–13]. Members of well-known rhizobial genera *Bradyrhizobium* [14] and *Mesorhizobium* [15] were initially classified into the genus *Rhizobium*, but were later reclassified to separate genera and subsequently placed into new families *Bradyrhizobiaceae* [16] and *Phyllobacteriaceae* [17].

*Onobrychis viciifolia* Scop. (Fabaceae), commonly referred to as common sainfoin, is an autochthonous leguminous plant with the putative origin in Central Asia. It was introduced to Europe in 14^th^ century and was intensively cultivated until the “green revolution”, during which it was replaced by higher-yielding legumes such as alfalfa (*Medicago sativa*). *Onobrychis viciifolia* is known as ‘healthy hay’ (from its old French name “Sain foin”) due to its positive effects on animal health and animal feeding [18–21]. Despite these positive traits, sainfoin is lacking a widespread application in agriculture in northern Europe. One reason may be the reports of inadequate levels of nitrogen fixation, resulting in nitrogen deficiency symptoms, despite the use of bacterial inocula [22–24]. Although sainfoin has been shown to reach similar rates of nitrogen fixation (130-160 kg/ha) as alfalfa (140-160 kg/ha) [25], the rate is highly dependent on the efficiency of the associated rhizobial symbiont [26]. Several rhizobia isolated from other legumes including *Coronilla* spp., *Hedysarum* spp., *Petalostemon* spp., *Oxytropis* spp., and *Astragalus alpinus* can also nodulate *O. viciifolia* [26, 27]. However, not many studies reported rhizobial strains nodulating sainfoin [28].

In rhizobia, nitrogenase genes are part of SGCs. Such large DNA fragments can be shared among bacteria by horizontal gene transfer via plasmids, integrative conjugative elements (ICEs) and/or genomic islands (GEIs) located on chromosomes. Andrews et al. [29] showed that symbiosis genes have been horizontally transferred within and between rhizobial genera. According to their gene content, GEIs and ICEs can be described as pathogenicity, symbiosis, metabolic, fitness or resistance islands [6, 30–32]. Both, pathogenic genome compartments (pathogenicity islands/virulence plasmids) and symbiotic genome compartments (symbiosis islands/symbiotic plasmids), convert environmental strains to strains that are able to form close pathogenic or symbiotic associations with eukaryotic hosts [33]. The community of rhizobial and nodule-associated bacteria is assumed to exchange plant-beneficial traits by transferring SGCs. As an example, Sullivan and co-workers found that the transfer of the symbiosis island of *Mesorhizobium loti* strain ICMP3153 (derivative R7A) converted non-symbiotic *Mesorhizobia* to plant symbionts [34].

In an attempt to identify rhizobial strains associated with sainfoin, different sainfoin varieties planted in an experimental field in the Leibniz Institute of Plant Genetics and Crop Plant Research (IPK), Germany were screened. In this context, two new strains were isolated from root nodules of sainfoin plants. We investigated these strains: *i)* to characterize them using *in silico* and *in vivo* studies, *ii)* to elucidate their taxonomic affiliation, plant growth-promoting traits repertoire, and iii) to evaluated their plant beneficial potential using greenhouse experiments.

## Material and Methods

### Plant material sample collection

One accession (ONO 20) and one cultivar (Taja) were selected for the present study. These plants were selected because of favourable characters like high tannin content and high biomass, respectively. ONO 20 is an old East German cultivar named *Bendelebener D 4 [35]*, which was included into the Genebank in 1958. Taja is a registered cultivar from the Polish breeder Malopolska Hodowla Roslin Spolka z.o.o in Krakow. The plants were cultivated on experimental fields of the Leibniz Institute of Plant Genetics and Crop Plant Research (IPK) in Gatersleben, Germany, during 2017-2019. The fields contain loamy soil, are very fertile and have high ground points (85-95).

### Isolation of bacteria from root nodules

Plant roots were washed to remove soil debris. Nodules were excised, surface sterilised for 1 min in 70 % ethanol, rinsed twice with sterile deionised water (SDW), followed by incubation in 1 % sodium hypochlorite (NaOCl) for 10 min and six rinses with SDW [36]. Surface sterilised nodules from each root sample were separately transferred to 2 ml microtubes and crushed with sterile pestles. Tubes were filled up with 1 ml of SDW or sterile 10 mM MgCl_2_ and vortexed for 1 min. An eight-fold serial dilution was made from 1 ml subsample of the homogenised suspension. A 100 µl subsample of each dilution was plated onto yeast mannitol agar (YMA; Sigma Aldrich, Merck KGaA, Darmstadt, Germany) supplemented with Congo red. Plates were incubated at 28°C and monitored daily for 8 days. The bacterial strains, including isolates studied here (OM4 and TH2) were stored at - 80°C.

### DNA extraction, sequencing and genome assembly

For details regarding DNA extraction, amplification and sequencing of partial 16S rRNA, *atpD* and *recA* genes, as well as complete genome sequencing and assembly, see Text S1.

### Phylogenetic analysis

Phylogenetic analysis was performed based on partial sequences of 16S rRNA gene and housekeeping genes *recA* and *atpD* and also a large number of conserved core genes. For more details, see Text S2.

### Overall genome relatedness indices

For genus and species delimitation, we calculated various overall genome relatedness indices (OGRIs), including whole-proteome average amino acid identity (wpAAI) [37–39], core-proteome average amino-acid identity (cpAAI), average nucleotide identity (ANI) [37, 40] and digital DNA-DNA hybridization (dDDH) [41]. To further determine the taxonomic position of the isolates studied here (TH2 and OM4), their genome sequences were subjected to the Type (Strain) Genome Server (TYGS) pipeline for a whole genome-based taxonomic analysis [42]. For more details, see Text S3.

### Plasmid similarity estimation

Plasmid similarity to known plasmid sequences was calculated via mash v.2.3 [43] in default dist mode. Respective reference plasmid sequences were received from the Plasmid database PLSDB version 2020_06_29 [44] and the Refseq plasmid collection stored on ftp://ftp.ncbi.nlm.nih.gov/refseq/release/plasmid/. For all plasmid reference hit sequences that showed at least one overlapping k-mer hash (Table S1) the pairwise mash distances were recalculated (sketch size 10 000; k-mer size 15) and visualized as neighbor network (NNet2004) by the outline algorithm with Splitstree 5 v.5.2.4 [45, 46].

### Comparative genomics and whole genome alignment

A pan-genome analysis was performed for both isolates, TH2 and OM4, separately, due to the different phylogenetic relationship, which was obtained by core-genome phylogenomic analysis. Computation of a common pan-genome of both isolates failed due to high evolutional distance between both, which led to a significantly decreasing number of core genes. For best comparability during downstream analysis, all genomes were annotated with Prokka v.1.14.6 [47]. Roary [48] v.3.13.0 [48] was applied to the annotated genomes of both isolates, using default parameters. The identity threshold (-i) was set differently according to the respective wpAAI values (Table S2) obtained for isolate TH2 (60 %) and isolate OM4 (80 %), considering the respective group gene/protein similarity. The single nucleotide polymorphism (SNP) tree of core genes was generated with FastTree v.2.1 [49] based on the maximum likelihood method and the generalized time reversible (GTR) model of evolution (parameters: -nt -gtr).

The genomes of isolate OM4 and its close relatives *M. delmotii* STM4623^T^ and *M. temperatum* SDW018^T^ were aligned using MAUVE (snapshot_2015_02_25, default parameters) [50] to find OM4-specific genomic features. Isolate OM4 specific unaligned regions that did not belong to any locally collinear block (LCB weight of 52) were extracted from the alignment file to analyse their functional characteristics and unique gene content.

### Genomic Functional annotation and visualization

Functional KEGG annotations were achieved for all isolates with the KOfamKOALA command line tool (https://www.genome.jp/tools/kofamkoala/) that applies HMM searches. KEGG comparisons between genomes were calculated and visualized with MEGAN6 [51] and custom python scripts. Genomic islands of the isolates TH2 and OM4 were detected online by IslandViewer 4 [52] using default parameters. Genomic prophage and phage-like regions were determined by the webtool PHASTER [53, 54]. AntiSMASH v. 6.0.1 [55] analysis allowed annotation of secondary metabolite biosynthesis gene clusters (BGCs). Selected annotation features were displayed as circular genome plots with BRIG [56]. Unique genes of isolate OM4 were analysed in more detail regarding their functional annotation and genomic position and affiliation to BGCs.

### Genes associated with plant-bacteria symbiosis and plant growth-promotion (PGP)

The KEGG annotations of the proteins of all strains were parsed into an IMG-like KEGG annotation file format via an in-house script and mapped against the plant growth-promotion traits ontology with the PGPT-Pred tool, available on the web platform for plant-associated bacteria PLaBAse (http://plabase.informatik.uni-tuebingen.de/pb/plabase.php [57]). The PGPT annotations of all strains were then merged for comparison. The PGPT density was calculated by division of the PGPT count by the total coding sequence count (CDS) of the respective genomic element (chromosome, plasmid or genomic region). The PGPT count comparison was plotted as z-scaled heatmap with iTol [58].

### Phenotypic characterization and fatty acid analysis

For details regarding phenotypic characterization and fatty acid analysis, see Text S4 and S5.

### Plant-growth promotion assays

Re-inoculation and nodulation tests were conducted as described in detail in Text S6.

## Results

### Phylogenetic inferences

A phylogenetic analysis based on partial sequence of the 16S rRNA gene showed that the isolate OM4 formed a highly supported monophyletic group with strains *Mesorhizobium delmotii* STM4623^T^, *M. prunaredense* STM4891^T^, *M. wenxiniae* WYCCWR 10195^T^, *M. muleiense* CCBAU 83963^T^, *M. robiniae* CCNWYC115^T^, *M. temperatum* SDW018^T^ and *M. mediterraneum* NBRC 102497^T^ (Fig. S1). Aanalyses of the housekeeping genes *recA* and *atpD* revealed a close relationship of OM4 and *M. prunaredense* STM4891^T^ with a high branch support (Figs. S2A and S2B). In addition, whole-genome sequence analysis demonstrated a distant relationship of these strains (see below).

The 16S rRNA gene sequence comparison of isolate TH2 with related *Rhizobiaceae* members suggested a close relationship with *Rhizobium alvei* strain TNR-22^T^ (Acc. No. HE649224.1), sharing 98.08 % nucleotide identity for an alignment length of 1 405 bp. This was below the stringent cutoff of 98.7 % 16S rRNA sequence identity, and proposed to delineate new species [59]. These two strains shared only 86.14 % and 87.65 % nucleotide identity for their partial *atpD* and *recA* gene sequences, respectively, suggesting their distinctiveness. The latter comparison was limited to 496 and 567 bp sequence lengths, because only partial *atpD* (Acc. No. KX938336) and *recA* (Acc. No. KX938338) sequences for *R. alvei* TNR-22^T^ were available. The 16S rRNA and *recA*-based phylogenetic analyses demonstrated that the isolate TH2 and *R. alvei* formed a monophyletic group with high support values (Figs. S3, S4A). The *atpD*-based analysis resulted in a tree with different topology, where TH2 did not cluster with *R. alvei*, but with other representatives of *Rhizobium, Agrobacterium* and *Ciceribacter* (Fig. S4B). Whole-genome analysis however, showed a distant relationship between these strains (see below).

### Core-genome phylogeny, overall genome relatedness indices, and plasmid comparison

Core-genome phylogeny was determined for isolate OM4 and TH2, and 99 additional *Rhizobiales* strains, including representatives of *Rhizobiaceae* and *Phyllobacteriaceae*. The core-genome of strains included in this analysis contained 180 homologous gene clusters. A phylogenomic tree was inferred from 118 top markers that were selected using GET_PHYLOMARKERS software.

The ML core-genome phylogeny indicated that the isolate OM4 grouped within the genus *Mesorhizobium* (Fig. 1). It clustered with strains *M. delmotii* STM4623^T^ and *M. temperatum* SDW018^T^ as its closest relatives. Isolate OM4 exhibited the highest genomic relatedness to these two strains, as they shared ∼94.8 % ANI (ANIb; Table S3). This was below the proposed threshold for species delineation, which ranges between 95-96 % for ANI [40]. To clarify taxonomic assignment of the isolate OM4, we calculated additional OGRIs, in particular orthoANIu and dDDH. Obtained values were also below the thresholds for species definition (Table S3). This suggests that isolate OM4 represents a novel *Mesorhizobium* species, for which we proposed the name *Mesorhizobium onobrychidis* sp. nov. (see Appendix). The novelty of *M. onobrychidis* strain OM4^T^ was also confirmed by TYGS analysis, suggesting that this strain does not belong to any species found on TYGS database (data not shown).

**Fig.1.**
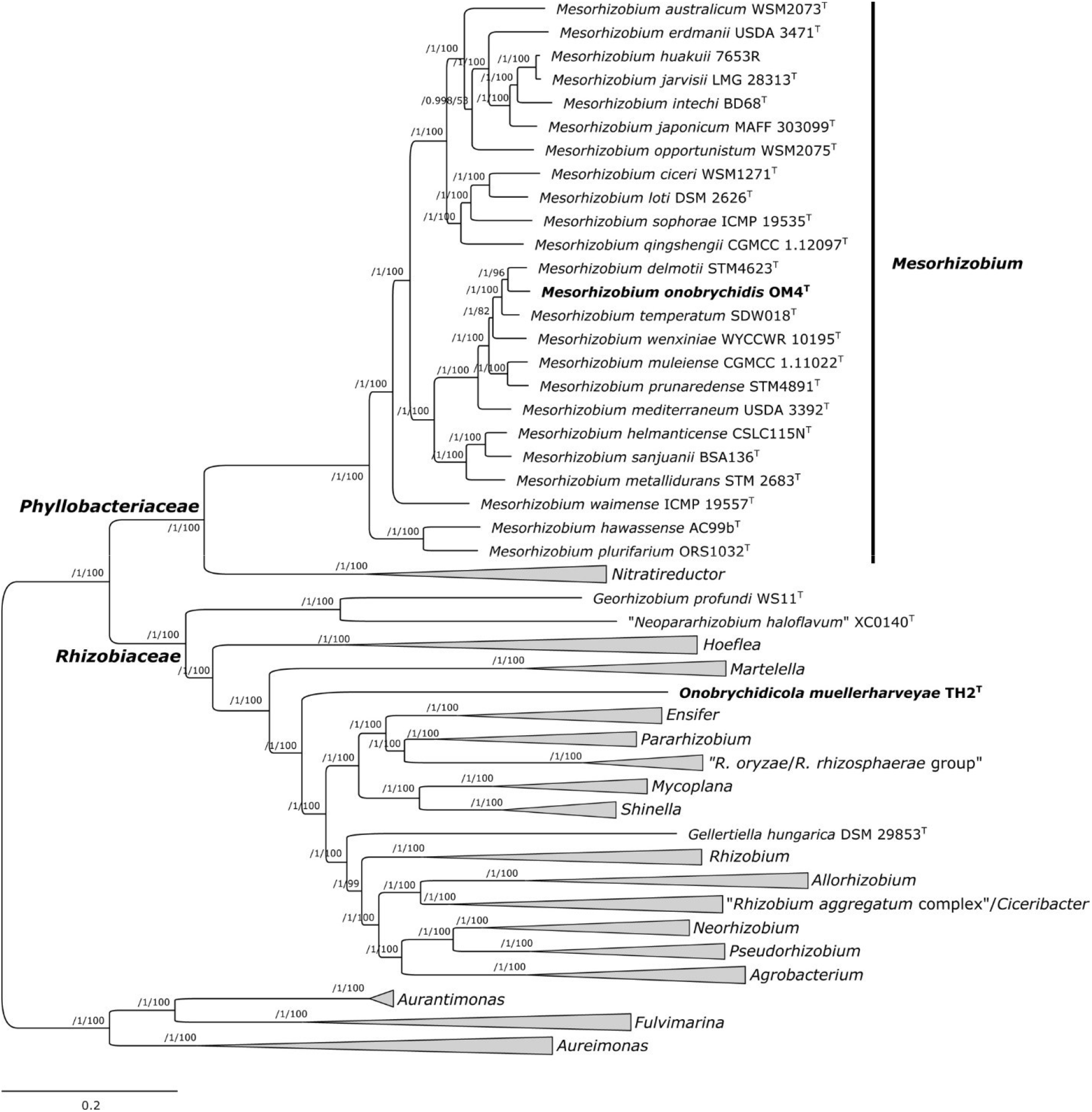
Maximum-likelihood core-genome phylogeny of strains O*nobrychidicola muellerharveyae* TH2^T^ and *Mesorhizobium onobrychidis* OM4^T^, including representatives of *Rhizobiaceae* and *Phyllobacteriaceae* (genera *Mesorhizobium* and *Nitratireductor*). The tree was estimated with IQ-TREE from the concatenated alignment of 118 top-ranked genes selected using GET_PHYLOMARKERS software. The numbers on the nodes indicate the approximate Bayesian posterior probabilities support values (first value) and ultra-fast bootstrap values (second value), as implemented in IQ-TREE. The tree was rooted using the sequences of representatives of genera *Aurantimonas, Aureimonas* and *Fulvimarina* as outgroup. The scale bar represents the number of expected substitutions per site under the best-fitting GTR+F+ASC+R8 model. The same tree, but without collapsing clades (thick gray branches), is presented as Fig. S7.

Phylogenetic analysis assigned isolate TH2 to *Rhizobiaceae* (Fig. 1). It clustered independently and was distantly related to other *Rhizobiaceae* genera described so far. Different OGRIs were computed to further assess the relationship of isolate TH2 to representatives of *Rhizobiaceae*. Because of the distinctiveness of this isolate, the comparisons at the nucleotide level were not satisfactory, and only a limited proportion of the whole-genome DNA sequence could be used for the calculations. For instance, for ANIb, only ∼12- 26 % of the whole-genome sequences were aligned and used for comparisons (data not shown). Therefore, we performed whole-proteome comparisons (wpAAI) that offer a higher resolution. Isolate TH2 exhibited wpAAI values ranging 61.5-67.5 % with the representatives of *Rhizobiaceae* included in the analysis (Table S2). The wpAAI values were notably low and supported the divergence of the isolate TH2, which was evidenced by the separate clustering of the strain on wpAAI dendrogram (Fig. S5). Isolate TH2 exhibited the highest genomic relatedness to strain *Ensifer meliloti* 1021 (67.5 % wpAAI), although they were phylogenetically distantly related (Fig. 1, Table S3). This value was lower than wpAAI values computed between representatives of defined and phylogenetically well-separated genera *Agrobacterium* and *Rhizobium* that ranged 68.12-70.55 % wpAAI. The cpAAI between the isolate TH2 and reference *Rhizobiaceae* spp. were <76 % (Table S4), which was below the threshold of ∼86 % for delimitation of *Rhizobiaceae* genera proposed recently [60]. This suggested that isolate TH2 represents a new genus and species, described here as *Onobrychidicola muellerharveyae* gen. nov sp. nov. Separate taxonomic position of strain *O. muellerharveyae* TH2^T^ was also confirmed by results of TYGS analysis (data not shown).

Plasmids of *M. onobrychidis* OM4^T^ and *O. muellerharveyae* TH2^T^ did not show high similarity to known plasmids based on mash analysis and pan-genome analysis and revealed a high proportion of unique genes (Fig. 2, Figs. S6 and Text S7).

**Figure 2:**
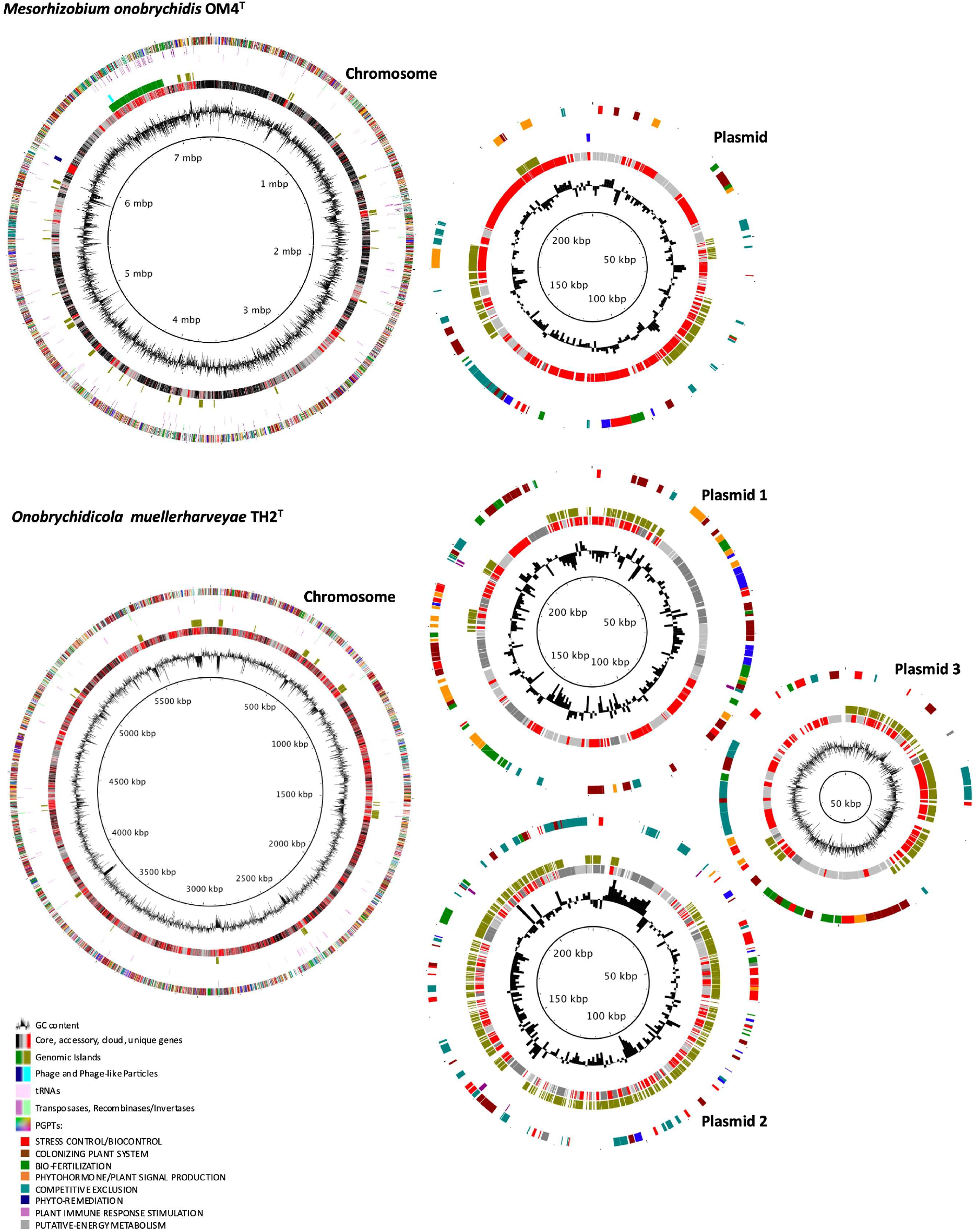
Genome annotation of *Mesorhizobium onobrychidis* OM4^T^ and *Onobrychidicola muellerharveyae* TH2^T^. Each chromosome and plasmid, respectively, is presented by a circular plot containing seven levels, of which the innermost circle 1 displays the G+C content of DNA. The circle 2 summarizes the Roary core-genome results with highlighted core (black), accessory (darkgrey), cloud (lightgrey), and strain-specific (unique) genes (red). Circle 3 presents distribution of genomic island genes predicted by IslandViewer version 4. Among the remaining circles. Circle 4 demonstrates the genes encoding phages or phage tail-like particles, circle 5 tRNAs and circle 6 transposases (violet) or recombinases / invertases (turquoise), i.e. enzymes enabling genome reshuffling. The outermost circle 7 presents genes annotated to plant growth-promoting traits (PGPTs) by PGPT-Pred, here subdivided into eight functional classes on PGPT ontology level 2.

### Genome sequencing and assembly

Genomes of strains *M. onobrychidis* OM4^T^ and *O. muellerharveyae* TH2^T^ were sequenced and circularized employing PacBio and Illumina platforms upon completion. Basic genome assembly statistics of both strains are summarized in Table 1. The complete genome size of strain *M. onobrychidis* OM4^T^ was 7.55 Mb, comprising the circular chromosome of 7.32 Mb and one circular plasmid of 227 kb, stored under the NCBI GenBank accession numbers: CP062229-CP062230 (Fig. 2). The G+C content of the total genome is 61.9 %. Genome size and G+C content of strain *M. onobrychidis* OM4^T^ are similar to other *Mesorhizobium* spp. (Table S5), for instance to the type strain of this genus, strain *M. loti* DSM 2626^T^ (Acc. No. QGGH01000000).

**Table 1:** Genome statistics of *Mesorhizobium onobrychidis* OM4^T^ and *Onobrychidicola muellerharveyae* TH2^T^

|  | *Mesorhizobium onobrychidis* OM4<sup>T</sup> | *Onobrychidicola muellerharveyae* TH2<sup>T</sup> |
| --- | --- | --- |
| Genome content | Chromosome (C) and one Plasmid (P1) | Chromosome (C) and three Plasmids (P1-3) |
| Genome size | Total: 7.55 Mb<br>(C: 7.32 Mb; P1: 227 kb) | Total: 6.44 Mb<br>(C: 5.88 Mb; P1: 238 kb; P2: 223 kb; P3: 98 kb) |
| GC content | C: 61.94 %; P1: 59.8 % | C: 60.64 %; P1: 59.75 %; P2: 57.55 %; P3: 56.31 % |
| Genes | 7,415 (7,347 CDS) | 6,373 (6,312 CDS) |
| Hypothetical genes | 3,779 | 3,157 |
| KEGG annotated genes | 3,676 | 3,365 |
| Unique genes (PG)* | 1,068 | 2,261 |
| Unique hypothetical genes (PG)* | 847 | 1,729 |
| rRNA operons (5S, 23S, 16S) | 2 | 2 |
| tRNAs | 62 | 53 |
| Phage, Phage-like particles (PLPs) and transposases | 2 PLPs; 136 transposases | 1 Phage; 3 PLPs; 18 transposases |
| Recombinases/Invertases (e.g. *xerC*, *xerD*) | 27 | 13 |
| PGPTs (density) | C: 3 973 (0.5545); P: 65 (0.3591) | C: 3 270 (0.57); P1: 110 (0.5263); P2: 227 (0.3910); P3: 50 (0.50) |
| Symbiosis Island (SI) or plasmid (SP) (density) with GC content | SI 1: 421 kb (0.7542); GC: 59.51 % |  |

The genome of strain *O. muellerharveyae* TH2^T^ composed of the circular chromosome (5.88 Mb) and three circular plasmids (98 kb, 223 kb and 238 kb), was deposited under the accession numbers CP062231-CP062234 at NCBI GenBank (Fig. 2). The genome size and G+C content of the total genome were 6.44 Mb and 60.6 %, respectively, which was similar to other representatives of *Rhizobiaceae* (Table S5). Two chromosomal rRNA (5S, 23S, 16S) operons were identified in both strains OM4^T^ and TH2^T^. In *O. muellerharveyae* TH2^T^ they were identical, while rRNA operons of strain *M. onobrychidis* OM4^T^ differed in five SNPs located in the intergenic region. For *M. onobrychidis* OM4^T^, two phage-like particles (PLPs) were identified, while in O. *muellerharveyae* TH2^T^ one phage and three PLPs were found (Fig. 2, circle 4). *Mesorhizobium onobrychidis* OM4^T^ harbored 136 transposases and 33 recombinases / invertases *e*.*g*., *xerC* and *xerD*, whereas *O. muellerharveyae* TH2^T^ revealed respective counts of 28 and 13 only (Fig 1, circle 6). Approximately half of the genes of both genomes were lacking meaningful annotations (hypothetical genes) according to homology-based alignment by PROKKA and KOfamKOALA hmm searches. According to the genomic island prediction tool IslandViewer 4, *M. onobrychidis* OM4^T^ -but not *O. muellerharveyae* TH2^T^, contains of a very large genomic island on its chromosome harbouring 414 genes (Fig. 2, circle 3; Fig. S8A). This is the only larger fragment of the *M. onobrychidis* OM4^T^ chromosome with high density of unique genes (Table S1). The genomic island on the *M. onobrychidis* OM4^T^ chromosome is located next to unique genes that are enriched in particular functions such as catalysing DNA exchange. The plant growth-promoting trait (PGPT) density of the genomic island is with 75 % considerably higher than the average PGPT density of the entire chromosome, which is only 55 % (see also Table 1). Details regarding the differences of PGPTs between *M. onobrychidis* OM4^T^, *O. muellerharveyae* TH2^T^, and other closely related strains are provided below.

### Comparative genomics and functional annotation

Pan-genome analysis of strains *M. onobrychidis* OM4^T^ and *O. muellerharveyae* TH2^T^ revealed a large number of gene clusters, ranging from 36 631 for all *Mesorhizobium* strains to 85 606 for all *Rhizobiaceae* strains here analysed (Fig. 2, circle 2, Fig. S9 A and B). The strain *M. onobrychidis* OM4^T^ revealed 2 683 core, 1 151 accessory, 2 444 cloud and 1 068 unique genes. While 428 unique genes could not be assigned to any KO number (KEGG annotations), 441 KO numbers were detected for *M. onobrychidis* OM4^T^ as unique genes, with various gene copy number. Functions of unique genes were associated with, among others, prokaryotic cellular community, signal transduction, carbohydrate and amino acid metabolism, cofactor and vitamin biosynthesis, energy metabolism, membrane transport and lipid metabolism (Fig. S10). In contrast, the putative novel genus comprising single strain *O. muellerharveyae* TH2^T^, revealed only 1 107 core genes, while counts of 1 839 for accessory, 1 105 for cloud, and 2 261 for unique genes were scored. The results did not allow further pan-genomic analysis for *O. muellerharveyae* TH2^T^ due to a distant phylogenetic relation between *O. muellerharveyae* TH2^T^ and the strains here analysed.

Overall, KEGG functional analysis and respective abundance clustering of all KEGG annotations confirmed that *M. onobrychidis* OM4^T^ contained functional similarities to *M. delmotii* STM4623^T^ and *M. temperatum* SDW018^T^. The analysis also supported the novelty of this species when considering only strain-specific enriched K numbers (Fig. S9 C). Analysing all K numbers for *O. muellerharveyae* TH2^T^ resulted in a distinct clustering, which became more distinct when considering only enriched ones (Fig. S9 D). Both patterns highly supported its status as a new genus.

The KEGG functional annotation for *M. onobrychidis* OM4^T^ and *O. muellerharveyae* TH2^T^ revealed two distinct clusters of level 2 and level 3 KEGG functions (Fig. S11, Text S8). *Onobrychidicola muellerharveyae* TH2^T^ showed higher counts for genes related to membrane transport, cell motility, cell growth and death, antimicrobial drug resistance, signal transduction and replication, repair, transport and catabolism (Fig. S11 A).

The detection of specific secondary metabolite biosynthesis gene clusters (BGCs) further confirmed the different lifestyles of strains OM4^T^ and TH2^T^ (Fig. S12, Text S9). The whole genome alignment of *Mesorhizobium* spp. revealed 85 regions unique to *M. onobrychidis* OM4^T^, harbouring at least five and up to 77 genes as one of its novel species characteristics (Fig. S13, Table S1, Text S10). Twenty-one regions could be assigned to seven of the entire 11 BGCs of OM4^T^. Among them, two BGCs matched with the genomic island, which covers 63 unaligned regions including 364 genes, all assigned as unique genes (Fig. S14, Table S2, Text 10).

### Functional PGPT annotation

The main genetic features and functional PGPT annotations based on KOfam-KEGG to PGPT mapping of all 80 strains, are summarised in a heatmap (Fig. 3). Detailed values are given in Table S1. The pattern of depleted (blue) and enriched (red) traits coincided with the phylogenetic clades – apart from very few exceptions in clade C. Heatmap fractions belonging to the *Mesorhizobium* clade (clade A), the *Ensifer* clade (clade D) and the *Rhizobium* clade (clade F) were dominated by PGPT classes of enriched gene counts.

**Fig. 3:**
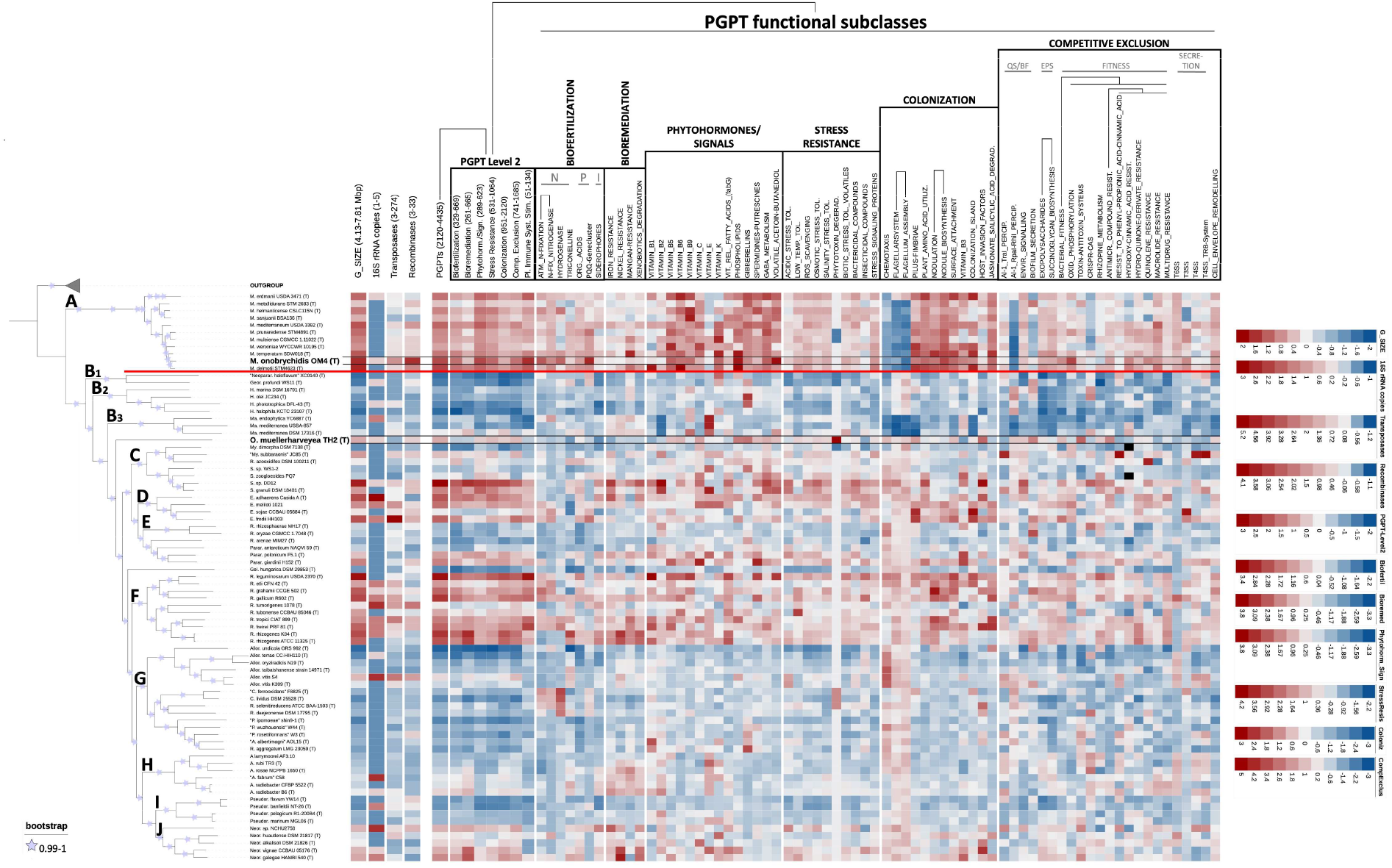
Functional PGPT heatmap based on KEGG annotations highlighting PGPTs abundant differences in functional classes and important genetic characteristics of *Mesorhizobium onobrychidis* OM4^T^, O*nobrychidicola muellerharveyae* TH2^T^, and other *Rhizobiaceae* and *Phyllobacteriaceae*. The reddish colour shows enriched and blueish colour depleted gene numbers based on a trait specific z-scale, which is given on the right. The phylogenetic tree provided on the left hand side allows better understanding of the PGPT distributions. Respective phylogenetic clades are highlighted by capital letters within the tree.

Focusing on the *Mesorhizobium* clade, a fraction of depleted traits refers to three subclasses of the PGPT class “colonization”, namely “chemotaxis” and “flagellar assembly”, important for bacteria to migrate towards chemical stimuli. In total 4 046 genes of *M. onobrychidis* OM4^T^ could be allocated to PGPTs compared to an average of 3 735 genes among other *Mesorhizobium* strains. A similar PGPT count was also found for its closest relative *M. delmotii* STM4623^T^. In general, the newly described species *M. onobrychidis* is very similar to the other species of genus *Mesorhizobium*. Among *Mesorhizobium, M. onobrychidis* OM4^T^ is one of the strains with the highest number of genes in the following PGP level 2 classes: biofertilization, phytohormone, plant signal production, stress resistance, competitive exclusion, and plant immune response stimulation. PGPT counts of *M. onobrychidis* OM4^T^ for the mentioned phytohormone and plant signals and plant immune system stimulation traits were higher compared to its two closest relatives *M. delmotii* STM4623^T^ and/or *M. temperatum* SDW018^T^ (Table S1). In contrast, *M. onobrychidis* strain OM4^T^ showed only an average amount of bioremediation genes, distinguishing it merely from *M. delmotii* STM4623^T^ and *M. temperatum* SDW018^T^. *Mesorhizobium onobrychidis* OM4^T^ harbored genes related to fixing carbon dioxide via RUBISCO as another highly plant beneficial feature (data not shown). It comprised a versatile set of stress resistance and colonization genes, their abundance mostly coincided with its both closest relatives. Furthermore, it contained the genetic ability for nodulation, vitamin B3 and pilus-fimbriae biosynthesis, the use of plant-derived metabolites *e*.*g*. amino acids, and the degradation of jasmonate/salicylic acid. Traits related to competitive exclusion showed a higher PGPT count for bacterial fitness compared to all other *Mesorhizobium* strains, especially for oxidative phosphorylation, resistance against plant antimicrobial compounds hydroxycinnamic acid and quinolene. The most significant differences between *M. onobrychidis* OM4^T^ and other *Mesorhizobium* strains occured in the number of transposases and *xerC*/*xerD* recombinases, which are important PGPTs related to colonization and competitive exclusion. *Mesorhizobium onobrychidis* OM4^T^ has approximately 2.5-times as many genes belonging to these categories as the other *Mesorhizobium* strains on average (transposases 136 compared to 57; recombinases 33 compared to 13). Regarding secretion systems, *M. onobrychidis* OM4^T^ encoded one T6SS, two T3SSs and one T4SS (*trb*) on its chromosome, as well as one copy of the *virB*-specific T4SS on its plasmid. The PGPT distribution alternated in shared pattern or highly varied between *M. onobrychidis* OM4^T^ and its relatives *M. delmotii* STM4623^T^ and/or *M. temperatum* SDW018^T^.

*Onobrychidicola muellerharveyae* TH2^T^ strongly differed in its overall PGPT abundancy profile from any other phylogenomic clades (Fig. 3, B1-3 and C-J). It contained a rather low number of genes for biofertilization and bioremediation, while the classes phytohormone and plant signaling, stress resistance, colonization and competitive exclusion were slightly above average. O*nobrychidicola muellerharveyae* TH2^T^ is one of the *Rhizobiaceae* strains with highest phospholipid and gibberellin encoding PGPT count. In terms of stress resistance, *O. muellerharveyae* TH2^T^ exceeds all other strains in the copy number of the gene for tabtoxin degradation (*ttr*), which is produced by some plant pathogens. While most *Rhizobiaceae* have one tabtoxin degradation gene (Fig. 3), *O. muellerharveyae* TH2^T^ contained four copies of this gene. In terms of competitive exclusion, *O. muellerharveyae* TH2^T^ was superior to all other investigated strains concerning the enrichment of genes for toxin-antitoxin systems (TASs). This is the case also in biofilm secretion and resistance to antimicrobial compounds. In terms of (host) colonization, *O. muellerharveyae* TH2^T^ was remarkable in subclasses “host invasion factors” and subclasses that enable target-oriented movement (“chemotaxis, flagellar system, flagellum assembly”). However, it lacked the nodulation gene cluster, despite possessing single nodulation-associated genes like *nolA*, and *nodD*. It showed an exceptional higher gene count for the plant branching inhibition and embryogenesis compounds spermidine and putrescine that act as plant signals. Regarding secretion systems, *O. muellerharveyae* TH2^T^ only harbored one T4SS (*virB*) on plasmid 2 and one T2SS on plasmid 3.

### Phenotypic characterization and fatty acid analysis

Phenotypic characteristics of strains *O. muellerharveyae* TH2^T^ and *M. onobrychidis* OM4^T^ are summarized in Table S6. Differential characteristics of *O. muellerharveyae* TH2^T^ and the type species from the other genera of family *Rhizobiaceae* are indicated in Table S5. Phenotypic tests performed with API 20NE system and Biolog GEN III microplates were assessed as unreliable, since negative reaction was observed for majority of tests (data not shown). This was likely because of the growth conditions that were inadequate for these strains. Therefore, most of the tests included in API 20NE system were repeated as conventional biochemical assays in test tubes, in order to facilitate the monitoring of bacterial growth and result assessment. Although more satisfactorily results were obtained in this way, no bacterial growth was observed for some tests *i*.*e*. in media containing L-tryptophane as a substrate (indole production test). For strain *M. onobrychidis* OM4^T^, no bacterial growth was observed in media containing aesculine ferric citrate (aesculine activity test) and gelatin (aesculin hydrolysis test) as substrates.

The results of the fatty acid analysis are summarised in the Table S7. The major fatty acids (>5 %) of *O. muellerharveyae* TH2^T^ are C_18:1_ w7c, C_19:0_ CYCLO w7c, C_16:0_ and C_17:0_ CYCLO w7c. Generally, as in other *Rhizobiaceae* members, the dominant fatty acid in *O. muellerharveyae* TH2^T^ was C_18:1_ w7c, which is in some strains comprised in Summed feature 8 (C_18:1_ w7c/C_18:1_ w6c). Unlike other type species from the other genera of *Rhizobiaceae, O. muellerharveyae* TH2^T^ contained relatively high (>5 %) amount of C_17:0_ CYCLO w7c. For *M. onobrychidis* OM4^T^ the major fatty acids (>5 %) were C_18:1_ w7c, C_16:0_, C_19:0_ CYCLO w7c, 11 methyl C_18:1_ w7c and C_18:0_, similarly as in other *Mesorhizobium* spp. [61].

### Plant nodulation and growth experiment

Nodulation and plant growth promotion assays confirmed Koch’s postulates for strains *M. onobrychidis* OM4^T^ and the control *R. leguminosarum* TS1-3-1. Both strains could be re-isolated from surface sterilized nodules. Re-isolation of *O. muellerharveyae* TH2^T^ failed for both single inoculations and co-inoculation with *R. leguminosarum* TS1-3-1. Sainfoin inoculated with *M. onobrychidis* OM4^T^ showed a statistically significant gain in aboveground biomass of all three tested sainfoin varieties (Fig. 4). Plants treated with *O. muellerharveyae* TH2^T^ did not exhibit increased biomass.

**Fig. 4:**
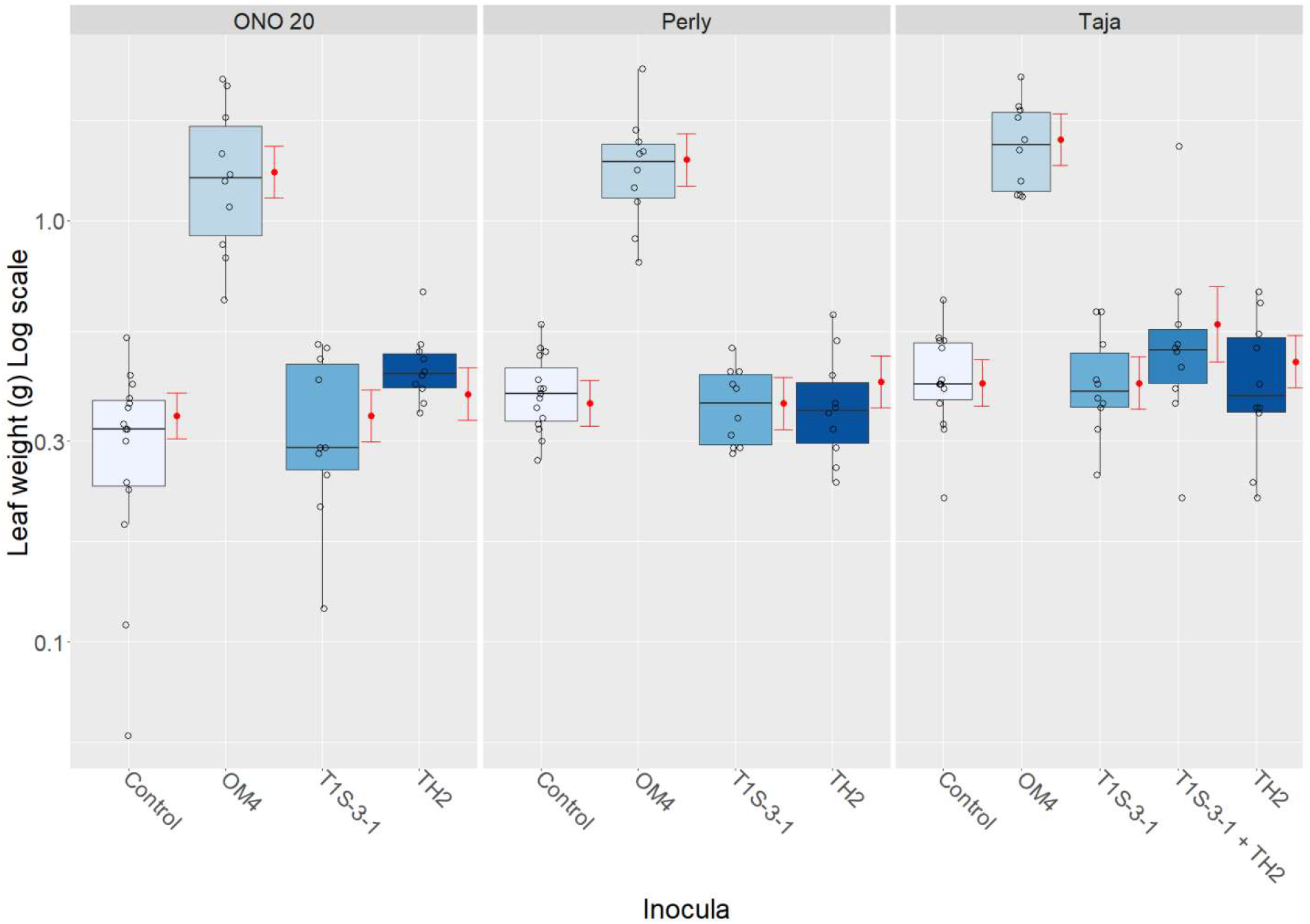
Plant biomass of nodulation assay using sainfoin accession ONO 20 and varieties Perly and Taja inoculated with *Mesorhizobium onobrychidis* OM4^T^ or O*nobrychidicola muellerharveyae* TH2^T^. *Rhizobium leguminosarum* T1S-3-1, known to induce nodulation, was used as a positive control. O*nobrychidicola muellerharveyae* TH2^T^ and *R. leguminosarum* T1S-3-1 were inoculated together into variety Taja as an attempt to piggyback O. *muellerharveyae* TH2^T^ into sainfoin plants. The negative control (“Control”) received no inoculation. Red dots and error-bars showing results of GLM statistical analysis.

## Discussion

### Bioinoculant potential and host interaction

The functional KEGG analyses revealed two contrasting settings for *M. onobrychidis* OM4^T^ and *O. muellerharveyae* TH2^T^, indicating two different lifestyles. Both strains differ in their competitive strategies, especially for colonizing the plant system. *In silico* analysis of PGPTs highlights the potential of *M. onobrychidis* OM4^T^ to improve plant performance via biofertilisation, phytohormone and plant signal production. Genes for root colonisation and adhesion by nodulation (*nod* gene cluster) and biotin metabolism were highly enriched in *M. onobrychidis* OM4^T^, whereas *O. muellerharveyae* TH2^T^ revealed a higher count for genes affiliated to motility, chemotaxis, and host invasion.

*Onobrychidicola muellerharveyae* TH2^T^ possessed all genes to assemble a complete flagellum with three different copies of the flagellin gene (*fliC*) allowing presumably higher diversity of flagellin epitopes acting as microbe-associated molecular patterns (MAMPs). Allelic variation of *fliC* is employed by bacteria to avoid the plant immune response [62]. O*nobrychidicola muellerharveyae* TH2^T^ lacked the nodulation gene cluster but harboured two nodulation-associated genes (*nolA* and *nodD*). Assumed that *O. muellerharveyae* TH2^T^ is not capable of inducing nodulation, this strain can be considered only as a nodule-associated strain. Our greenhouse experiments confirmed the *in silico* analysis.

Although it carries a large amount of PGP genes, *O. muellerharveyae* TH2^T^ showed no effect on sainfoin plants in our inoculation experiment under nitrogen-limited conditions. O*nobrychidicola muellerharveyae* TH2^T^ might achieve better potential under biotic stress condition [9], as its highest number of genes were found in functional classes referring to bacterial fitness/ stress tolerance. In contrast to most other *Rhizobiaceae*, which have only one copy, *O. muellerharveyae* TH2^T^ had four copies of the tabtoxin degradation gene (*ttr*). Plant pathogens such as *Pseudomonas syringae* produce tabtoxin for chlorosis and lesion formation [63] and carry a *ttr* gene for self-protection from tabtoxinine-beta-lactam [64]. It can be assumed that the *ttr* gene products of *O. muellerharveyae* TH2^T^ diminish the deleterious effect of phytotoxin-producing bacteria.

Among all analysed bacteria, *O. muellerharveyae* TH2^T^ contained the highest fraction of genes belonging to toxin-antitoxin systems (TAS). It is uncertain whether TAS provides any advantage to its host plant, since plant pathogenic bacteria such as *Xylella fastidiosa* also employ TASs [65]. It has been argued that TASs do not necessarily provide an advantage to the producing bacterial strain. For example, chromosomal TASs of *Pseudomonas putida* were reported to be rather selfish than beneficial, and an indirect positive effect for plants cannot be ruled out [66]. This example on TAS illustrates the need for further functional studies of particular genes, and shows the difficulty to assign them to a unique purpose. Further effort is needed to identify plant beneficial traits for robust and reliable prediction of PGPTs. However, *M. onobrychidis* OM4^T^ provides a convincing example that *in silico* analyses can already be used for identification of bacterial strains exhibiting a beneficial impact on plants. Here we showed that comparative genomics of PGPTs, based on the novel ontology, is a solid tool that considers widely acknowledged PGP pathways such as nitrogen and carbon dioxide fixation together as one entity (bio-fertilization). The use of bioinformatics for determining genomic islands *e*.*g*. via IslandViewer combined with PGPT enrichment analyses demands a reclassification of symbiotic island and symbiotic plasmids, as not all criteria defined by Ling et al. [67] match. The lack of these genes in the genus *Mesorhizobium* (exception for strain USDA 3471) and in *M. onobrychidis* OM4^T^ suggests that most of the *Mesorhizobium* strains included in our *in silico* analyses are immotile or at least do not move by means of flagella.

### Genetic features of OM4

*Mesorhizobium onobrychidis* OM4^T^ possessed 136 transposases (and 33 *xerC*/*xerD* recombinases), which is extraordinarily high among the representatives of *Phyllobacteriaceae* and *Rhizobiaceae* used in our analyses (Tab. 1, Fig. 3). A higher rate of transposable elements can be related to sessile endosymbiotic bacteria [68]. However, this pattern was associated with reductive genome evolution of such sessile strains, which is not given for the strain described here. It has been discussed that the development of the nitrogen-fixing symbioses in legume nodules required co-evolution of legumes and rhizobia [69]. Zhao et al. [70] however showed that adaptive evolution of symbiotic compatibility could be achieved by spontaneous transposition of inserted sequences (ISs). This was demonstrated by the observation that different *Sinorhizobium* strains do form either nitrogen-fixing nodules or uninfected pseudonodules [70]. Next to ISs, site-specific recombinases *xerC* and *xerD* contribute to genome plasticity and mediate *e*.*g*. formation and resolution of plasmid co-integrates [71]. It was shown that *xerC* is crucial for competitive root colonization [72], [73]. Accordingly, the high number of *xer*C / *xer*D genes in *M. onobrychidis* OM4^T^ suggests its competitive root colonization ability [72, 74].

Particular secretion systems are known to be crucial for bacteria/plant interaction (T1SS, T3SS, T4SS) and competitive plant colonization (T6SS) [75], and plasmid transfer across the rhizobial community (T4SS) [76]. Type III secretion systems (T3SSs) are well known for effector translocation into eukaryotic host cells and thus a major mediator for pathogenicity [77, 78]. Such systems are however found to be present in symbiotic bacteria, where they contribute to a stable host-microbe interaction [79, 80]. *Mesorhizobium onobrychidis* OM4^T^ encodes two T3SSs and two T4SSs suggesting an effective interaction with its host.

One genomic region of *M. onobrychidis* OM4^T^ fulfilled some but not all of the criteria of a symbiosis island defined by Ling and co-workers [67]. The tool IslandViewer however supported its nature as a genomic island. The fact of a presence of higher density of PGPTs on this island compared to the density of the total genome raised the question if the criteria of a symbiosis island have to be extended by the PGPT density. PGPT annotation is challenging and not standardized. The use of the novel PLaBAse database and supporting online tools close this gap [57].

Based on our functional analysis, the following functional characteristics for *M. onobrychidis* OM4^T^ can be proposed. It is a rather sessile strain, as it lacks the genes for “chemotaxis” and “flagellar assembly”. The strain is adapted to the plant metabolism, as it does not harbour an enriched set of carbohydrate, amino acid and nucleotide metabolic genes compared to other plant-associated bacteria here analysed. It carries a remarkable set of direct plant-growth promotion traits and might achieve its colonization towards or inside the plant via biofilms and/or seed transmission [81].

The observed growth promotion during the greenhouse experiments suggests the bacterium as an ‘efficient’ rhizobial species for sainfoin (*O. viciifolia*) under nitrogen-limited plant growth conditions.

## Conclusion

The economic benefit of these newly discovered species still needs to be determined, but their phylogenetically distant position suggests them as interesting research subjects. O*nobrychidicola muellerharveyae* TH2^T^ is the type strain of the monotypic genus *Onobrychidicola*. Since closely related strains have not been described, a large fraction of its genes is unique due to the overall low homology of genes. *Onobrychidicola muellerharveyae* TH2^T^ carries a high proportion of PGPTs that contributes to colonisation, stress resistance and competitiveness, rather than to direct plant beneficial effects. Sufficient PGP potential for commercial application needs to be determined further in *in planta* experiments. Due to its likely potential to antagonize phytopathogens, the strain still could be considered for biocontrol purposes while developing alternatives to chemical pesticides.

A number of recent studies suggested sainfoin to be integrated in modern and sustainable agriculture due to its beneficial properties on animal nutrition, and animal and soil health [18]. Overall performance of sainfoin highly depends on an effective symbiosis with rhizobial strains, many of which do not meet the plants nitrogen-requirements [26]. The study presented here, describes the well-performing novel plant growth-promoting bacterial species *M. onobrychidis* OM4^T^, which is supported by its PGPTs. Our greenhouse experiments showed that this bacterium can be inoculated into a variety of sainfoin cultivars to improve their biomass production, and might be a promising candidate for application in a sustainable agricultural system.

## Supporting information

Table S1 Genome annotation

Table S2 AAI matrix

## Appendix

### Formal descriptions of two new *Rhizobiales* species, including new *Rhizobiaceae* genus

The following paragraphs provide formal descriptions (protologues) of the new *Rhizobiaceae* genus and the two new *Rhizobiales* species (Fig. 5).

**Fig. 5.**
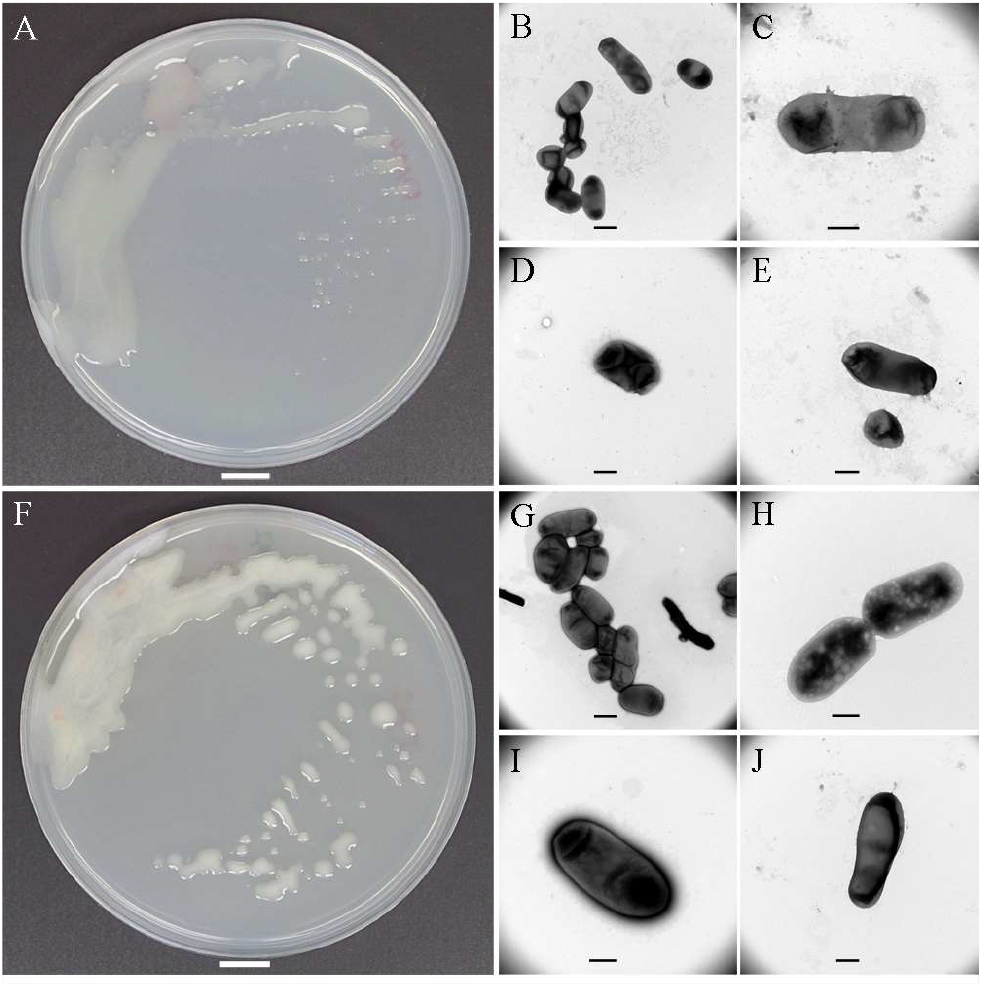
Colonies and cells of *Mesorhizobium onobrychidis* strain OM4^T^ and O*nobrychidicola muellerharveyae* strain TH2^T^. A) colonies of OM4^T^ on YMA after 10 days incubation at 28 °C, B-E) TEM micrographs of cells of OM4^T^ grown in YMA. F) colonies of *O. muellerharveyae* strain TH2^T^ on YMA after 10 days incubation G-J) TEM micrographs of cells of *O. muellerharveyae* strain TH2^T^ grown in YMA liquid. Scale: A, F: 100 mm, B, G: 1 µm, C, D, E, H, I 500 nm.

### Description of *Mesorhizobium onobrychidis* sp. nov

*Mesorhizobium onobrychidis* (o.no.bry’chi.dis. N.L. gen. n. *onobrychidis* of the plant genus *Onobrychis*). Cells are Gram-negative, non-spore-forming and rod-shaped, 1.1 – 2.3 µm (1.7 ± 0.3 SD) in length, 0.7 – 1.15 µm (0.95 ± 0.1 SD) wide, (n=15). They are non-motile and non-flagellated. Aerobic, oxidase and catalase positive. Bacteria grow on YMA, TY and R2A medium. Colonies very slow growing, on YMA medium appearing within 8-10 days, white, glistening, circular, and convex, 1 mm diameter after 10 days incubation at 28°C. Growth is observed at temperatures between 10 and 25°C. Nitrate reduction is negative. Glucose fermentation is negative. Arginine dihydrolase and β-galactosidase tests are negative. Production of urease is positive. Glucose, L-arabinose, D-mannose, D-mannitol, D-maltose, gluconate and malate are assimilated. A weak assimilation was observed for caprate. Adipate, citrate and phenylacetate are not assimilated. The major fatty acids (>5 %) are C_18:1_ w7c, C_16:0_, C_19:0_ CYCLO w7c, 11-methyl C_18:1_ w7c and C_18:0_. OM4^T^ induces effective nodules on its original host plant (cvs. Taja, Perly, ONO 20). Additionally, genes involved in legume nodulation and nitrogen fixation were identified in the genome of the strain OM4^T^.

The genome size of the type strain (OM4^T^) is 7.55 Mb. The genome is composed of the circular chromosome (7.32 Mb) and circular plasmid (227 kb). The G+C content of total genomic and/or chromosomal DNA is 61.9 %.

The type strain OM4^T^ (=DSM 109849 =NCCB 100791) was isolated from a root nodule of *Onobrychis viciifolia*, Germany, in 2019. The DDBJ/EMBL/GenBank accession numbers for the genome sequence are CP062229 (chromosome) and CP062230 (plasmid).

### Description of *Onobrychidicola* gen. nov

*Onobrychidicola* (O.no.bry.chi.di’co.la. N.L. fem. n. *Onobrychis*, a plant genus; L. suff. –cola [from L. masc. or fem. n. *incola*], inhabitant, dweller; N.L. masc. n. *Onobrychidicola*, a dweller of *Onobrychis*). Cells are aerobic, Gram-negative, non-spore-forming, rod-shaped, non-motile and non-flagellated. Oxidase and catalase positive. The major fatty acids (>5 %) are C_18:1_ w7c, C_19:0_ CYCLO w7c, C_16:0_ and C_17:0_ CYCLO w7c. The G + C content of total genomic DNA of the type strain of the type species is 60.4 and 60.6 %, respectively. The genus *Onobrychidicola* has been separated from other *Rhizobiaceae* genera based on core-genome phylogeny, whole-and core-proteome comparisons (wpAAI and cpAAI, respectively), as well as pan-genome and functional analyses.

The type species is *Onobrychidicola muellerharveyae*.

### Description of *Onobrychidicola muellerharveyae* sp. nov

*Onobrychidicola muellerharveyae* (muel.ler.har’vey.ae. N.L. gen. n. muellerharveyae named in honour of Dr. Irene Mueller-Harvey, for her outstanding work on sainfoin).

Cells are aerobic, Gram-negative, non-spore-forming, rod-shaped 1.44–2.63 µm (2.0 ± 0.3) in length, 0.8– 1.4 µm (1.1 ± 0.1) wide (n= 20), non-motile and non-flagellated, oxidase and catalase positive bacteria that shows relatively good growth on YMA, TY and R2A medium. Colonies slow growing on YMA medium whitish to pale creamy, variable in size and shape, 1—4 mm diameter after 10 days growth at 28 °C. Growth was observed at temperature range between 10 and 30 °C. Nitrate reduction and glucose fermentation are negative. Arginine dihydrolase and gelatin hydrolysis tests are negative. Production of urease and esculin hydrolysis are positive. Production of β-galactosidase is weakly positive. D-mannose and D-glucose are assimilated. A weak assimilation was observed for L-arabinose, D-mannitol, D-maltose, gluconate and caprate. Adipate, malate, citrate and phenylacetate are not assimilated. The major fatty acids (>5 %) are C_18:1_ w7c, C_19:0_ CYCLO w7c, C_16:0_ and C_17:0_ CYCLO w7c. The strain TH2^T^ could not induce nodules on its original host plant. Accordingly, genes involved in legume nodulation and nitrogen fixation were absent in the genome of the strain TH2^T^.

The genome size of the type strain (TH2^T^) is 6.44 Mb. The genome is composed of the circular chromosome (5.88 Mb) and three circular plasmids (98-238 kb). The G+C content of total genomic and chromosomal DNA is 60.4 and 60.6 %, respectively.

The type strain TH2^T^ (=DSM 109848^T^ =NCCB 100790^T^) was isolated from a root nodule of *Onobrychis viciifolia* in Germany, in 2019. The DDBJ/EMBL/GenBank accession numbers for the genome sequence are CP062231-CP062234.

## Acknowledgements

This work was partially funded by the Federal Ministry of Food and Agriculture (BMEL) based on a decision of the Parliament of the Federal Republic of Germany via the Agency of Renewable Resources (FNR). The work of NK was funded by the Deutsche Forschungsgemeinschaft (DFG, German Research Foundation) – Projektnummer 429677233.

Special thanks to Dr. Roland Kölliker for providing sainfoin cultivar Taja from the Polish breeder Malopolska Hodowla Roslin Spolka z.o.o in Krakow. The authors gratefully acknowledge Prof. Aharon Oren (The Hebrew University of Jerusalem, Jerusalem, Israel), Prof. Bernhard Schink (University of Konstanz, Konstanz, Germany) and Prof. George M. Garrity (Michigan State University, East Lansing, MI, USA) for their valuable help on nomenclature aspects. We thank Kristin Müller, Heike Bosse, Gesa Martens for excellent technical support for molecular studies, culture maintenance, physiological analyses, and plant maintains. We thank Jessica Ponath for valuable contribution to TEM studies. We thank Simone Severitt and Nicole Heyer for excellent technical assistance regarding complete genome sequencing. Special thank goes to Prof. Daniel H. Huson from the University of Tübingen for providing his facilities for batch genome annotation and comparison. We would like to thank Drs Yvonne Becker and Wolfgang Maier for facilitating laboratory and greenhouse experiments and Dr. Ahmed Elhady for scientific discussions during the laboratory works. This research was partially enabled through computational resources provided by BMBF-funded de.NBI Cloud within the German Network for Bioinformatics Infrastructure (de.NBI) (031A537B, 031A533A, 031A538A, 031A533B, 031A535A, 031A537C, 031A534A, 031A532B).

## Authors contribution

SA and TT conceived and designed the study. SA, UL, ML and TT carried out the plant growth experiments. SA, NK, AF, SV and MN performed phenotypic and physiological tests. SA, NK, SP, MB and TT performed the data analysis and figure drawing. BB and CS performed whole genome sequencing. All authors contributed to drafting and revising the manuscript.

## Data availability

Genome Sequences are available at NCBI GenBank under the accession numbers CP062229-CP062230 and CP062231-CP062234, respectively. Sequences of single genes are available at NCBI GenBank under the accession numbers MW915806 - MW915808, and MW917139 - MW917144.

## Compliance with ethical standards

Not applicable

## References

[1] Hellriegel H, Wilfarth H. Untersuchungen über die Stickstoffnahrung der Gramineen und Leguminosen, von H. Hellriegel und H. Wilfarth unter mitwirkung von H. Roemer, R. Günther, H. Moeller und G. Wimmer. (Referent: H. Hellriegel.). Berlin: Buchdruckerei der “Post” Kayssler 1888.

[2] Lajudie PM de, Young JPW. International committee on systematics of Prokaryotes subcommittee for the taxonomy of Rhizobium and Agrobacterium minutes of the meeting, Budapest, 25 August 2016. Int J Syst Evol Microbiol 2017; 67(7): 2485–94 [https://doi.org/10.1099/ijsem.0.002144][PMID: 28771120]

[3] Hirsch AM, Lum MR, Downie JA. What makes the rhizobia-legume symbiosis so special? Plant Physiol. 2001; 127(4): 1484–92 [https://doi.org/10.1104/pp.010866]

[4] Andrews M, Meyer S de, James EK, et al. Horizontal Transfer of Symbiosis Genes within and Between Rhizobial Genera: Occurrence and Importance. Genes (Basel) 2018; 9(7) [https://doi.org/10.3390/genes9070321][PMID: 29954096]

[5] Remigi P, Zhu J, Young JPW, Masson-Boivin C. Symbiosis within Symbiosis: Evolving Nitrogen-Fixing Legume Symbionts. Trends Microbiol 2016; 24(1): 63–75 [https://doi.org/10.1016/j.tim.2015.10.007][PMID: 26612499]

[6] González V, Bustos P, Ramírez-Romero MA, et al. The mosaic structure of the symbiotic plasmid of Rhizobium etli CFN42 and its relation to other symbiotic genome compartments. Genome Biol 2003; 4(6): R36 [https://doi.org/10.1186/gb-2003-4-6-r36][PMID: 12801410]

[7] Martínez-Hidalgo P, Hirsch AM. The nodule microbiome: N 2-fixing rhizobia do not live alone. Phytobiomes Journal 2017; 1(2): 70–82 [https://doi.org/10.1094/PBIOMES-12-16-0019-RVW]

[8] Peix A, Ramírez-Bahena MH, Velázquez E, Bedmar EJ. Bacterial associations with legumes. Critical Reviews in Plant Sciences 2015; 34(1-3): 17–42 [https://doi.org/10.1080/07352689.2014.897899]

[9] Tokgöz S, Lakshman DK, Ghozlan MH, Pinar H, Roberts DP, Mitra A. Soybean Nodule-Associated Non-Rhizobial Bacteria Inhibit Plant Pathogens and Induce Growth Promotion in Tomato. Plants (Basel) 2020; 9(11): 1494 [https://doi.org/10.3390/plants9111494][PMID: 33167465]

[10] Yao LJ, Shen YY, Zhan JP, Xu W, Cui GL, Wei GH. Rhizobium taibaishanense sp. nov., isolated from a root nodule of Kummerowia striata. Int J Syst Evol Microbiol 2012; 62(Pt 2): 335–41 [https://doi.org/10.1099/ijs.0.029108-0][PMID: 21421926]

[11] Yan J, Li Y, Han XZ, et al. Agrobacterium deltaense sp. nov., an endophytic bacteria isolated from nodule of Sesbania cannabina. Arch Microbiol 2017; 199(7): 1003–9 [https://doi.org/10.1007/s00203-017-1367-0][PMID: 28386665]

[12] Delamuta JRM, Scherer AJ, Ribeiro RA, Hungria M. Genetic diversity of Agrobacterium species isolated from nodules of common bean and soybean in Brazil, Mexico, Ecuador and Mozambique, and description of the new species Agrobacterium fabacearum sp. nov. Int J Syst Evol Microbiol 2020; 70(7): 4233–44 [https://doi.org/10.1099/ijsem.0.004278][PMID: 32568030]

[13] Wang ET, Tan ZY, Willems A, Fernández-López M, Reinhold-Hurek B, Martínez-Romero E. Sinorhizobium morelense sp. nov., a Leucaena leucocephala-associated bacterium that is highly resistant to multiple antibiotics. Int J Syst Evol Microbiol 2002; 52(Pt 5): 1687–93 [https://doi.org/10.1099/00207713-52-5-1687][PMID: 12361275]

[14] Jordan DC. NOTES: Transfer of Rhizobium japonicum Buchanan 1980 to Bradyrhizobium gen. nov., a genus of slow-growing, root nodule bacteria from leguminous plants. International Journal of Systematic Bacteriology 1982; 32(1): 136–9 [https://doi.org/10.1099/00207713-32-1-136]

[15] Jarvis BDW, van Berkum P, Chen WX, et al. Transfer of Rhizobium loti, Rhizobium huakuii, Rhizobium ciceri, Rhizobium mediterraneum, and Rhizobium tianshanense to Mesorhizobium gen. nov. Int J Syst Evol Microbiol 1997; 47(3): 895–8 [https://doi.org/10.1099/00207713-47-3-895]

[16] Garrity GM, Bell JA, and Lilburn T. Family VII. Bradyrhizobiaceae fam. nov. In: Brenner DJ, Krieg NR, Staley JT, Garrity GM, editors. Bergey’s Manual® of systematic bacteriology: Volume Two The Proteobacteria Part C The Alpha-, Beta-, Delta-, and Epsilonproteobacteria. 2nd ed. 2005. New York, NY: Springer US 2005; 438.

[17] Mergaert J, Swings J. Family IV. Phyllobacteriaceae fam. nov. In: Brenner DJ, Krieg NR, Staley JT, Garrity GM, editors. Bergey’s Manual® of systematic bacteriology: Volume Two The Proteobacteria Part C The Alpha-, Beta-, Delta-, and Epsilonproteobacteria. 2nd ed. 2005. New York, NY: Springer US 2005; 393.

[18] Mora-Ortiz M, Smith LMJ. Onobrychis viciifolia ; a comprehensive literature review of its history, etymology, taxonomy, genetics, agronomy and botany. Plant Genet. Resour. 2018; 16(5): 403–18 [https://doi.org/10.1017/S1479262118000230]

[19] Hayot Carbonero C, Mueller-Harvey I, Brown TA, Smith L. Sainfoin (Onobrychis viciifolia): a beneficial forage legume. Plant Genet. Resour. 2011; 9(01): 70–85 [https://doi.org/10.1017/S1479262110000328]

[20] McMahon LR, McAllister TA, Berg BP, et al. A review of the effects of forage condensed tannins on ruminal fermentation and bloat in grazing cattle. Can. J. Plant Sci. 2000; 80(3): 469–85 [https://doi.org/10.4141/P99-050]

[21] Sheppard SC, Cattani DJ, Ominski KH, Biligetu B, Bittman S, McGeough EJ. Sainfoin production in western Canada: A review of agronomic potential and environmental benefits. Grass Forage Sci 2019; 74(1): 6–18 [https://doi.org/10.1111/gfs.12403]

[22] Burton JC, Curley RL. Nodulation and nitrogen fixation in sainfoin (Onobrychis sativa, Lam.) as influenced by strains of rhizobia. Bull. Mont. agric. Exp. Stn. 1970; 627: 3–5.

[23] Sims JR, Muir MK, Carleton AE. Evidence of ineffective rhizobia and its relation to the nitrogen nutrition of sainfoin. Bull. Montana. Agricultural Experimental Station 1970; 627: 8–12.

[24] Sheehy JE, Popple SC. Photosynthesis, water relations, temperature and canopy structure as factors influencing the growth of sainfoin (Onobrychis viciifolia Scop.) and Lucerne (Medicago sativa L.). Annals of Botany 1981; 48(2): 113–28.

[25] Provorov NA, Tikhonovich IA. Genetic resources for improving nitrogen fixation in legume-rhizobia symbiosis. Genetic Resources and Crop Evolution 2003; 50(1): 89–99 [https://doi.org/10.1023/A:1022957429160]

[26] Prévost D, Bordeleau LM, Antoun H. Symbiotic effectiveness of indigenous arctic rhizobia on a temperate forage legume: Sainfoin (Onobrychis viciifolia). Plant Soil 1987; 104(1): 63–9 [https://doi.org/10.1007/BF02370626]

[27] Prévost D, Bordeleau LM, Caudry-Reznick S, Schulman HM, Antoun H. Characteristics of rhizobia isolated from three legumes indigenous to the Canadian high arctic: Astragalus alpinus, Oxytropis maydelliana, and Oxytropis arctobia. Plant Soil 1987; 98(3): 313–24 [https://doi.org/10.1007/BF02378352]

[28] Laguerre G, van Berkum P, Amarger N, Prévost D. Genetic diversity of rhizobial symbionts isolated from legume species within the genera Astragalus, Oxytropis, and Onobrychis. Appl Environ Microbiol 1997; 63(12): 4748–58 [https://doi.org/10.1128/AEM.63.12.4748-4758.1997][PMID: 9406393]

[29] Andrews M, Andrews ME. Specificity in legume-rhizobia symbioses. Int J Mol Sci 2017; 18(4) [https://doi.org/10.3390/ijms18040705][PMID: 28346361]

[30] Sengupta M, Austin S. Prevalence and significance of plasmid maintenance functions in the virulence plasmids of pathogenic bacteria. Infect. Immun. 2011; 79(7): 2502–9 [https://doi.org/10.1128/IAI.00127-11][PMID: 21555398]

[31] Juhas M, van der Meer JR, Gaillard M, Harding RM, Hood DW, Crook DW. Genomic islands: tools of bacterial horizontal gene transfer and evolution. FEMS Microbiol Rev 2009; 33(2): 376–93 [https://doi.org/10.1111/j.1574-6976.2008.00136.x][PMID: 19178566]

[32] Bañuelos-Vazquez LA, Torres Tejerizo G, Brom S. Regulation of conjugative transfer of plasmids and integrative conjugative elements. Plasmid 2017; 91: 82–9 [https://doi.org/10.1016/j.plasmid.2017.04.002][PMID: 28438469]

[33] Sullivan JT, Ronson CW. Evolution of rhizobia by acquisition of a 500-kb symbiosis island that integrates into a phe-tRNA gene. Proceedings of the National Academy of Sciences 1998; 95(9): 5145–9 [https://doi.org/10.1073/pnas.95.9.5145][PMID: 9560243]

[34] Sullivan JT, Patrick HN, Lowther WL, Scott DB, Ronson CW. Nodulating strains of Rhizobium loti arise through chromosomal symbiotic gene transfer in the environment. Proceedings of the National Academy of Sciences 1995; 92(19): 8985–9 [https://doi.org/10.1073/pnas.92.19.8985][PMID: 7568057]

[35] Schieblich J. Beitrag zur Züchtung von Esparsette (Onobrychis viciaefolia Scop.). Der Züchter 1951; 21: 132–6.

[36] Elhady A, Heuer H, Hallmann J. Plant parasitic nematodes on soybean in expanding production areas of temperate regions. J Plant Dis Prot 2018; 125(6): 567–76 [https://doi.org/10.1007/s41348-018-0188-y]

[37] Goris J, Konstantinidis KT, Klappenbach JA, Coenye T, Vandamme P, Tiedje JM. DNA-DNA hybridization values and their relationship to whole-genome sequence similarities. Int J Syst Evol Microbiol 2007; 57(Pt 1): 81–91 [https://doi.org/10.1099/ijs.0.64483-0][PMID: 17220447]

[38] Konstantinidis KT, Tiedje JM. Towards a genome-based taxonomy for prokaryotes. J Bacteriol 2005; 187(18): 6258–64 [https://doi.org/10.1128/JB.187.18.6258-6264.2005][PMID: 16159757]

[39] Konstantinidis KT, Rosselló-Móra R, Amann R. Uncultivated microbes in need of their own taxonomy. ISME J 2017; 11(11): 2399–406 [https://doi.org/10.1038/ismej.2017.113][PMID: 28731467]

[40] Richter M, Rosselló-Móra R. Shifting the genomic gold standard for the prokaryotic species definition. Proc Natl Acad Sci U S A 2009; 106(45): 19126–31 [https://doi.org/10.1073/pnas.0906412106][PMID: 19855009]

[41] Meier-Kolthoff JP, Auch AF, Klenk H-P, Göker M. Genome sequence-based species delimitation with confidence intervals and improved distance functions. BMC Bioinformatics 2013; 14: 60 [https://doi.org/10.1186/1471-2105-14-60][PMID: 23432962]

[42] (YYYYYYY) Meier-Kolthoff JP, Göker M. TYGS is an automated high-throughput platform for state-of-the-art genome-based taxonomy. Nat Commun 2019; 10(1): 2182 [https://doi.org/10.1038/s41467-019-10210-3][PMID: 31097708]

[43] Ondov BD, Treangen TJ, Melsted P, et al. Mash: fast genome and metagenome distance estimation using MinHash. Genome Biol 2016; 17(1): 132 [https://doi.org/10.1186/s13059-016-0997-x][PMID: 27323842]

[44] Galata V, Fehlmann T, Backes C, Keller A. PLSDB: a resource of complete bacterial plasmids. Nucleic Acids Res 2019; 47(D1): D195–D202 [https://doi.org/10.1093/nar/gky1050][PMID: 30380090]

[45] Huson DH, Bryant D. Application of phylogenetic networks in evolutionary studies. Mol Biol Evol 2006; 23(2): 254–67 [https://doi.org/10.1093/molbev/msj030][PMID: 16221896]

[46] Bagci C, Bryant D, Cetinkaya B, Huson DH. Microbial Phylogenetic Context Using Phylogenetic Outlines. Genome Biol Evol 2021; 13(9) [https://doi.org/10.1093/gbe/evab213][PMID: 34519776]

[47] Seemann T. Prokka: rapid prokaryotic genome annotation. Bioinformatics 2014; 30(14): 2068–9 [https://doi.org/10.1093/bioinformatics/btu153][PMID: 24642063]

[48] Page AJ, Cummins CA, Hunt M, et al. Roary: rapid large-scale prokaryote pan genome analysis. Bioinformatics 2015; 31(22): 3691–3 [https://doi.org/10.1093/bioinformatics/btv421][PMID: 26198102]

[49] Price MN, Dehal PS, Arkin AP. FastTree 2--approximately maximum-likelihood trees for large alignments. PLoS One 2010; 5(3): e9490.[https://doi.org/10.1371/journal.pone.0009490][PMID: 20224823]

[50] Darling ACE, Mau B, Blattner FR, Perna NT. Mauve: multiple alignment of conserved genomic sequence with rearrangements. Genome Res 2004; 14(7): 1394–403 [https://doi.org/10.1101/gr.2289704][PMID: 15231754]

[51] Huson DH, Auch AF, Qi J, Schuster SC. MEGAN analysis of metagenomic data. Genome Res 2007; 17(3): 377–86 [https://doi.org/10.1101/gr.5969107][PMID: 17255551]

[52] Bertelli C, Laird MR, Williams KP, et al. IslandViewer 4: expanded prediction of genomic islands for larger-scale datasets. Nucleic Acids Res 2017; 45(W1): W30–W35 [https://doi.org/10.1093/nar/gkx343]

[53] Arndt D, Grant JR, Marcu A, et al. PHASTER: a better, faster version of the PHAST phage search tool. Nucleic Acids Res 2016; 44(W1): W16–21 [https://doi.org/10.1093/nar/gkw387][PMID: 27141966]

[54] Zhou Y, Liang Y, Lynch KH, Dennis JJ, Wishart DS. PHAST: a fast phage search tool. Nucleic Acids Res 2011; 39(Web Server issue): W347–52 [https://doi.org/10.1093/nar/gkr485][PMID: 21672955]

[55] Blin K, Shaw S, Kloosterman AM, et al. antiSMASH 6.0: improving cluster detection and comparison capabilities. Nucleic Acids Res 2021; 49(W1): W29–W35 [https://doi.org/10.1093/nar/gkab335][PMID: 33978755]

[56] Alikhan N-F, Petty NK, Ben Zakour NL, Beatson SA. BLAST Ring Image Generator (BRIG): simple prokaryote genome comparisons. BMC Genomics 2011; 12(1): 402 [https://doi.org/10.1186/1471-2164-12-402][PMID: 21824423]

[57] Patz S, Gautam A, Becker M, Ruppel S, Rodríguez-Palenzuela P, Huson D. PLaBAse: A comprehensive web resource for analyzing the plant growth-promoting potential of plant-associated bacteria 2021.

[58] Letunic I, Bork P. Interactive Tree Of Life (iTOL) v5: an online tool for phylogenetic tree display and annotation. Nucleic Acids Res 2021; 49(W1): W293–W296 [https://doi.org/10.1093/nar/gkab301][PMID: 33885785]

[59] Chun J, Oren A, Ventosa A, et al. Proposed minimal standards for the use of genome data for the taxonomy of prokaryotes. Int J Syst Evol Microbiol 2018; 68(1): 461–6 [https://doi.org/10.1099/ijsem.0.002516][PMID: 29292687]

[60] Kuzmanović N, Fagorzi C, Mengoni A, Lassalle F, diCenzo GC. Taxonomy of Rhizobiaceae revisited: proposal of a new framework for genus delimitation 2021.

[61] Nguyen TM, van Pham HT, Kim J. Mesorhizobium soli sp. nov., a novel species isolated from the rhizosphere of Robinia pseudoacacia L. in South Korea by using a modified culture method. Antonie Van Leeuwenhoek 2015; 108(2): 301–10 [https://doi.org/10.1007/s10482-015-0481-8][PMID: 25980835]

[62] Clarke CR, Chinchilla D, Hind SR, et al. Allelic variation in two distinct Pseudomonas syringae flagellin epitopes modulates the strength of plant immune responses but not bacterial motility. New Phytol 2013; 200(3): 847–60 [https://doi.org/10.1111/nph.12408][PMID: 23865782]

[63] Bender CL, Alarcón-Chaidez F, Gross DC. Pseudomonas syringae phytotoxins: mode of action, regulation, and biosynthesis by peptide and polyketide synthetases. Microbiol Mol Biol Rev 1999; 63(2): 266–92 [https://doi.org/10.1128/MMBR.63.2.266-292.1999][PMID: 10357851]

[64] Wencewicz TA, Walsh CT. Pseudomonas syringae self-protection from tabtoxinine-β-lactam by ligase TblF and acetylase Ttr. Biochemistry 2012; 51(39): 7712–25 [https://doi.org/10.1021/bi3011384][PMID: 22994681]

[65] Merfa MV, Niza B, Takita MA, Souza AA de. The MqsRA Toxin-Antitoxin System from Xylella fastidiosa Plays a Key Role in Bacterial Fitness, Pathogenicity, and Persister Cell Formation. Front Microbiol 2016; 7: 904 [https://doi.org/10.3389/fmicb.2016.00904][PMID: 27375608]

[66] Rosendahl S, Tamman H, Brauer A, Remm M, Hõrak R. Chromosomal toxin-antitoxin systems in Pseudomonas putida are rather selfish than beneficial. Sci Rep 2020; 10(1): 9230 [https://doi.org/10.1038/s41598-020-65504-0][PMID: 32513960]

[67] Ling J, Wang H, Wu P, et al. Plant nodulation inducers enhance horizontal gene transfer of Azorhizobium caulinodans symbiosis island. Proc Natl Acad Sci U S A 2016; 113(48): 13875–80 [https://doi.org/10.1073/pnas.1615121113][PMID: 27849579]

[68] Ran L, Larsson J, Vigil-Stenman T, et al. Correction: Genome Erosion in a Nitrogen-Fixing Vertically Transmitted Endosymbiotic Multicellular Cyanobacterium. PLoS One 2010; 5(9) [https://doi.org/10.1371/annotation/835c5766-5128-41c4-b636-adfe0c503103]

[69] La Coba de Peña T, Fedorova E, Pueyo JJ, Lucas MM. The Symbiosome: Legume and Rhizobia Co-evolution toward a Nitrogen-Fixing Organelle? Front Plant Sci 2017; 8: 2229 [https://doi.org/10.3389/fpls.2017.02229][PMID: 29403508]

[70] Zhao R, Liu LX, Zhang YZ, et al. Adaptive evolution of rhizobial symbiotic compatibility mediated by co-evolved insertion sequences. ISME J 2018; 12(1): 101–11 [https://doi.org/10.1038/ismej.2017.136][PMID: 28800133]

[71] Cameranesi MM, Morán-Barrio J, Limansky AS, Repizo GD, Viale AM. Site-Specific Recombination at XerC/D Sites Mediates the Formation and Resolution of Plasmid Co-integrates Carrying a blaOXA-58-and TnaphA6-Resistance Module in Acinetobacter baumannii. Front Microbiol 2018; 9: 66 [https://doi.org/10.3389/fmicb.2018.00066][PMID: 29434581]

[72] Dekkers LC, Phoelich CC, van der Fits L, Lugtenberg BJ. A site-specific recombinase is required for competitive root colonization by Pseudomonas fluorescens WCS365. Proceedings of the National Academy of Sciences 1998; 95(12): 7051–6 [https://doi.org/10.1073/pnas.95.12.7051][PMID: 9618537]

[73] Dekkers LC, Mulders IH, Phoelich CC, Chin-A-Woeng TF, Wijfjes AH, Lugtenberg BJ. The sss colonization gene of the tomato-Fusarium oxysporum f. sp. radicis-lycopersici biocontrol strain Pseudomonas fluorescens WCS365 can improve root colonization of other wild-type pseudomonas spp.bacteria. Molecular Plant-Microbe Interactions 2000; 13(11): 1177–83 [https://doi.org/10.1094/MPMI.2000.13.11.1177][PMID: 11059484]

[74] Cornet F, Hallet B, Sherratt DJ. Xer recombination in Escherichia coli. Site-specific DNA topoisomerase activity of the XerC and XerD recombinases. Journal of Biological Chemistry 1997; 272(35): 21927–31 [https://doi.org/10.1074/jbc.272.35.21927][PMID: 9268326]

[75] Lucke M, Correa MG, Levy A. The Role of Secretion Systems, Effectors, and Secondary Metabolites of Beneficial Rhizobacteria in Interactions With Plants and Microbes. Front Plant Sci 2020; 11: 589416 [https://doi.org/10.3389/fpls.2020.589416][PMID: 33240304]

[76] Trokter M, Waksman G. Translocation through the Conjugative Type IV Secretion System Requires Unfolding of Its Protein Substrate. J Bacteriol 2018; 200(6) [https://doi.org/10.1128/JB.00615-17][PMID: 29311273]

[77] Coburn B, Sekirov I, Finlay BB. Type III secretion systems and disease. Clin Microbiol Rev 2007; 20(4): 535–49 [https://doi.org/10.1128/CMR.00013-07][PMID: 17934073]

[78] Tang X, Xiao Y, Zhou J-M. Regulation of the type III secretion system in phytopathogenic bacteria. Molecular Plant-Microbe Interactions 2006; 19(11): 1159–66 [https://doi.org/10.1094/MPMI-19-1159][PMID: 17073299]

[79] Dale C, Plague GR, Wang B, Ochman H, Moran NA. Type III secretion systems and the evolution of mutualistic endosymbiosis. Proceedings of the National Academy of Sciences 2002; 99(19): 12397–402 [https://doi.org/10.1073/pnas.182213299][PMID: 12213957]

[80] Songwattana P, Noisangiam R, Teamtisong K, et al. Type 3 Secretion System (T3SS) of Bradyrhizobium sp. DOA9 and Its Roles in Legume Symbiosis and Rice Endophytic Association. Front Microbiol 2017; 8: 1810 [https://doi.org/10.3389/fmicb.2017.01810][PMID: 28979252]

[81] Mora Y, Díaz R, Vargas-Lagunas C, et al. Nitrogen-fixing rhizobial strains isolated from common bean seeds: phylogeny, physiology, and genome analysis. Appl Environ Microbiol 2014; 80(18): 5644–54 [https://doi.org/10.1128/AEM.01491-14][PMID: 25002426]

