## Supplementary Information for "Two new *Rhizobiales* species isolated from root nodules of common sainfoin (*Onobrychis viciifolia*) show different plant colonization strategies"

**List of supplementary data** **Page**


**Text**

| | | |
|---|---|---|
| Text S1 | DNA extraction, sequencing and genome assembly | 3 |
| Text S2 | Phylogenetic analysis | 4 |
| Text S3 | Overall genome relatedness indices | 5 |
| Text S4 | Phenotypic characterization | 5 |
| Text S5 | Fatty acid analysis | 6 |
| Text S6 | Plant-growth promotion assays | 6 |
| Text S7 | Plasmid similarity | 7 |
| Text S8 | Functional comparison | 7 |
| Text S9 | Secondary metabolite biosynthesis gene clusters | 7 |
| Text S10 | Whole genome alignment | 7 |

**Tables**

| | | |
|---|---|---|
| Table S1 | Genome annotation | 9 |
| Table S2 | Whole-proteome average amino acid identity (wpAAI) comparisons | 9 |
| Table S3 | Pairwise OGRI comparisons | 9 |
| Table S4 | Core-proteome average amino acid identity (cpAAI) comparisons | 10 |
| Table S5 | Differential characteristics of *Rhizobiaceae* | 11 |
| Table S6 | Phenotypic characteristics of the new strains | 12 |
| Table S7 | Cellular fatty acid composition of the new strains | 12 |
| Table S8 | List of strains and GenBank/EMBL/DDBJ accession numbers | 13 |

**Figures**

| | | |
|---|---|---|
| Figure S1 | 16S RNA based phylogenetic analysis of *Mesorhizobium onobrychidis* | 14 |
| Figure S2 | *recA* and *atpD* based phylogenetic analysis of *Mesorhizobium onobrychidis* | 15 |
| Figure S3 | 16S RNA based phylogenetic analysis of *Onobrychidicola muellerharveyae* | 16 |
| Figure S4 | *recA* and *atpD* based phylogenetic analysis of *Onobrychidicola muellerharveyae* | 17 |
| Figure S5 | Dendrogram and heatmap based on average amino acid identity (AAI) matrics | 18 |
| Figure S6 | Outline Neighbor Network for plasmids based on mash distances | 19 |
| Figure S7 | Core-genome phylogenetic tree (uncollapsed) | 20 |
| Figure S8 | Genomic islands and phage annotations | 21 |
| Figure S9 | Pan-genome analysis and KEGG abundance clustering | 22 |
| Figure S10 | Functional annotation of unique genes | 23 |
| Figure S11 | KEGG annotations of the new strains | 24 |
| Figure S12 | AntiSMASH analyses of the new strains | 25 |
| Figure S13 | Whole-genome MAUVE alignment | 26 |
| Figure S14 | AntiSMASH analysis of secondary metabolite biosynthesis gene cluster | 27 |

**Literature** **28**

**Text S1 -** DNA extraction, sequencing and genome assembly

Genomic DNA was extracted from bacterial cultures grown on yeast mannitol agar (YMA, Sigma Aldrich, Merck KGaA, Darmstadt, Germany) at 28°C for 3 days, using a silica-based kit (Silica bead DNA Gel Extraction Kit; Thermo Scientific, St. Leon-Rot, Germany) according to the manufacturer's instructions. The extracted genomic DNA was stored at -20°C until further uses.

Three genes were amplified and partially sequenced: 16S rRNA gene, and the housekeeping genes *atpD* and *recA*. The primer pairs U8-27 (AGAGTTTGATCMTGGCTCAG) [1] and R1494-1514 (CTACGGYTACCTTGTTACGAC) [2], 273F (SCTGGGSCGYATCMTGAACGT) and 771R (GCCGACACTTCCGAACCNGCCTG), and *rec*A-6F (CGKCTSGTAGAGGAYAAATCGGTGGA) and *rec*A-555R (CGRATCTGGTTGATGAAGATCACCAT) [3] were used for amplification and sequencing of 16S rRNA, *atpD* and *recA* genes, respectively.

The 16S rRNA gene fragment was amplified in a 25 µl PCR reaction containing TrueStart Buffer, 3.75 mM MgCl2, 0.2 mM dNTP, 5 % w/v DMSO, 0.1 mg/ml BSA, 0.1 mM forward and reverse primers, 1 U TrueStart-Taq DNA Polymerase (Thermo Scientific, Germany), and 1 µl of DNA template. The PCR conditions consisted of the initial denaturation step at 94°C for 5 min, 30 cycles of 94°C for 1 min, 56°C for 1 min, 72°C for 1 min, and a final elongation step at 72°C for 10 min. The partial housekeeping genes *atpD* and *recA* were amplified in a 25µl volume with master mix containing 1 × Colourless GoTaq Flexi buffer (Promega Corp., Madison, WI, USA), 1.5 mmol $l^{-1}$ $MgCl_2$, 0.2 mmol $l^{-1}$ of each dNTP, 0.2 µmol $l^{-1}$ of each primer, 0.5 U of GoTaq Flexi DNA polymerase (Promega Corp., Madison, WI, USA), and 1 µl of DNA template. The thermal programs were performed as described by Kuzmanovic et al. [4]. All amplifications were carried out on a T-GRADIENT thermocycler (Biometra, Goettingen, Germany). The PCR products were purified using the DNA Clean & ConcentratorTM-5 kit (Zymo Research Corp., Irvine, CA, USA) according to the manufacturer's instructions, and sequence on an ABI 3730XL sequencing machine (Eurofins Genomics GmbH, Ebersberg, Germany). Obtained sequences were assembled, edited, and trimmed with Sequencher 5.4.1 (Gene Codes Corporation, Ann Arbor, MI, USA), and deposited in GenBank under accession numbers MW915806 - MW915808, and MW917139 - MW917144. Using BLASTn searches [5], sequence similarity of all strains were compared with the publically available defined species.

High molecular weight DNA was prepared using Qiagen Genomic Tip/100 G (Qiagen, Hilden, Germany) according to the manufacturer´s instructions. SMRTbell™ template library was prepared according to the instructions from PacificBiosciences, Menlo Park, CA, USA, following the Procedure & Checklist – Greater Than 10 kb Template Preparation. Briefly, for preparation of 15 kb libraries 8 µg genomic DNA was sheared using g-tubes™ (Covaris, Woburn, MA, USA) according to the manufacturer´s instructions. DNA was end-repaired and ligated overnight to hairpin adapters applying components using the DNA/Polymerase Binding Kit P6 (Pacific Biosciences, Menlo Park, CA, USA). Reactions were carried out according to the instructions of the manufacturer. BluePippin™ Size-Selection to greater than 4 kb was performed according to the manufacturer´s instructions (Sage Science, Beverly, MA, USA). Conditions for annealing of sequencing primers and binding of polymerase to purified SMRTbell™ template were assessed with the Calculator in RS Remote (PacificBiosciences, Menlo Park, CA, USA). SMRT sequencing was carried out on the PacBio *RSII* (PacificBiosciences, Menlo Park, CA, USA) taking one 240-minutes movie on one SMRT cell per sample using the P6 Chemistry.

In addition, short-read Illumina sequencing was also performed. DNA libraries (paired-end) with insert length of approx. 300 bp were obtained with Nextera XT DNA Library Prep Kit (Illumina, San Diego, CA,

USA). Paired-end sequencing (2 × 151 bp) was performed on an Illumina NextSeq 500 platform. Adapter clipping was done using bcl2fastq2 conversion software.

The genome sequences reported in this study have been deposited at DDBJ/EMBL/GenBank under the accession numbers CP062231-CP062234 (*O. muellerharveyae* strain TH2$^T$) and CP062229-CP062230 (*M. onobrychidis* strain OM4$^T$).

SMRT Cell data was assembled using the "RS_HGAP_Assembly.3" protocol included in SMRT Portal version 2.3.0 using default parameters. Replicons revealed were circularized and adjusted to *dnaA* (chromosomes) or *repA* (chromids and plasmids) as the first gene. Error-correction was performed by a mapping of paired-end reads of 2x150 bp generated on an Illumina NextSeq 500 onto finished genomes using BWA [6] and/or Bowtie2 [7] with subsequent variant and consensus calling using VarScan [8]. The obtained replicons with mapped reads were also visually inspected to correct remaining errors. A consensus concordance of QV60 could be confirmed for the genome. The genome sequences were annotated using Prokka (Galaxy Version 1.13) [9] and NCBI Prokaryotic Genomes Annotation Pipeline (PGAP) [10].

**Text S2 -** Phylogenetic analyses

A dataset of 16S rRNA gene sequences including the newly generated sequences as well as of the representatives of *Rhizobiaceae* and *Phyllobacteriaceae* described so far was prepared. The latter were extracted from LPSN - List of Prokaryotic names with Standing in Nomenclature (https://lpsn.dsmz.de/). Two other datasets of the housekeeping genes *recA* and *atpD* were obtained from already published sequences available in GenBank. DNA sequences were aligned automatically using the online version of MAFFT v. 7 (https://mafft.cbrc.jp/alignment/server) [11]. Alignments were visually examined and manually trimmed using AliView v. 1.26 [12].

Phylogenetic analyses were applied using Bayesian Inference (BI), and maximum-likelihood (ML). Maximum-likelihood analyses were performed using the web server of IQ-Tree2 (http://www.iqtree.org/) [13] following the default parameters, the best-fitting substitution model and Ultrafast Bootstrap (1000) [14]. Bayesian inference analyses was performed through Markov Chain Monte Carlo (MCMC) sampling based on best fitting models settings in MrBayes v3.2 [15]. A 50 % majority rule consensus tree was computed only from the trees of the plateau, and if, additionally, the split frequencies were below 0.01. The trees representing the “burn-in phase” were discarded and the remaining trees were used to infer posterior probabilities (PP). The best-fit model of DNA substitution was estimated using MrModeltest v2.2 [16].

Core-genome-based phylogenetic analysis was performed on a dataset containing two strains studied here (*O. muellerharveyae* TH2$^T$ and *M. onobrychidis* OM4$^T$) and 99 reference strains belonging to *Phyllobacteriaceae* and *Rhizobiaceae* (Table S8). As outgroups, representatives of genera *Aurantimonas*, *Aureimonas* and *Fulvimarina* were included. GenBank files generated by Prokka were used as an input for the script get_homologues.pl implemented into GET_HOMOLOGUES software package Version 11042019 [17]. Computations were performed by bidirectional best-hit (BDBH), Clusters of Orthologous Groups-triangles (COGtriangles), and OrthoMCL (Markov Clustering of orthologs, OMCL) algorithms using a stringent 90 % coverage cut-off for BLASTP alignments (-C 90). A consensus core-genome was computed as the intersection of the clusters computed by the BDBH, COG-triangles and OMCL algorithms by employing script compare_clusters.pl (-t 101, number of genomes). Furthermore, the resulting core-genome clusters were processed and used for phylogenomic analysis by the pipeline for DNA-based phylogenies (-R 1 -t DNA) of GET_PHYLOMARKERS software package Version 2.2.8_18Nov2018 [18]. In particular, ML phylogeny was estimated under the best-fitting substitution model using IQ-TREE v1.6.10 [19].

**Text S3 -** Overall genome relatedness indices

The software CompareM Version 0.0.23 (https://github.com/dparks1134/CompareM) was used for computation of whole-proteome average amino acid identity (wpAAI) [20–22] values using default options. To visualize wpAAI results Morpheus software (https://software.broadinstitute.org/morpheus) was used to generate heatmaps and for hierarchical clustering (metric: matrix values [for a precomputed similarity values]; linkage method: single; cluster: rows and columns), using pairwise identity matrix as an input. Moreover, core-proteome average amino-acid identity (cpAAI) between the strain TH2 and reference *Rhizobiaceae* spp. was computed using cpAAI_*Rhizobiaceae* pipeline (github.com/flass/cpAAI_Rhizobiaceae). For this purpose, a pre-aligned reference set of 170 non-recombining core protein markers conserved among a set of 97 *Rhizobiaceae* genomes was used [23].

The average nucleotide identity (ANI) [20, 24] between the strains was calculated using PyANI program Version 0.2.9, with scripts employing BLAST+ (ANIb) algoritm to align the input sequences (https://github.com/widdowquinn/pyani). Additionally, ANI values were calculated by OrthoANIu Version 1.2 (calculates orthologous ANI using USEARCH algorithm) [25]. The digital DNA-DNA hybridization (dDDH) values by the Genome-to-Genome Distance Calculator (GGDC 2.1; http://ggdc.dsmz.de/distcalc2.php) using the recommended BLAST+ alignment and formula 2 (identities/HSP length) [26].

To further determine the taxonomic position of strains TH2 and OM4, their genome sequences were subjected to the Type (Strain) Genome Server (TYGS) pipeline for a whole genome-based taxonomic analysis [27], involving thorough comparison of query genomes with a comprehensive collection of microbial type-strain genomes. The results were provided by the TYGS on 2021-11-23.

**Text S4 -** Phenotypic characterization

Colony morphology was observed on YMA medium. The presence of flagella and motility was tested by soft agar (0.1 % yeast extract, 0.01 % $K_2HPO_4$, 0.2 % agar) and monitored microscopically after growth in liquid medium [28, 29]. The motility of bacteria was also assessed on LB, TY (DSMZ medium 1143) and Motility medium reported by Bouzar et al. [30]. Cell size, shape, arrangement and flagellation were examined with transmission electron microscopy (TEM). For TEM, cells from an exponentially grown culture on TY and YMA were negatively stained with 1% (w/v) uranyl acetate on a carbon-coated Formvar grid. After air drying, the grid was examined using a FEI Tecnai G2 Spirit Twin transmission electron microscope (ThermoFisher Scientific, Germany).

The gram reaction was determined by KOH [31] and aminopeptidase [32]; Bactident Aminopeptidase, (Merck, Cat. No.113301, Germany) tests and confirmed by gram-staining. Oxidase activity was tested by the method of Kovacs et al. [33]. Catalase test was performed by mixing freshly grown bacterial cells with 10% $H_2O_2$, followed by examination of gas bubble formation. Growth at 10, 15, 20, 25, 30, 37 and 40°C was determined in YMA modified medium (DSMZ medium 1704) for up to 14 days. Additionally, the strains TH2 and OM4 were phenotypically characterized by using API 20NE system (bioMérieux, Marcv L'Etoile, France) following the instructions provided by the manufacturer. All tests performed in the API

20NE strip, except for β-galactosidase and N-acetyl-glucosamine assimilation tests, which were repeated as conventional biochemical assays in test tubes. Nitrate reduction, indole production, glucose fermentation, arginine dihydrolase production, urease production, esculin hydrolysis and gelatin hydrolysis were tested in YMA-based medium. Substrate utilization (glucose to phenylacetic acid) was tested in mineral salt liquid medium (0.8 g/L $K_2HPO_4$, 0.2 g/L $KH_2PO_4$, 0.2 g/L $MgSO_4$ x $7H_2O$, 0.005 g/L $NH_4$-FeIII-Citrate, 1 g/L $(NH_4)_2SO_4$, 0.05 g/l $CaCl_2$ x $2H_2O$ and 3.3 g/L biomaris sea salt) supplemented with the corresponding substrates (2 g/L) at 28°C for up to 14 days. Furthermore, utilization of sole carbon sources and chemical sensitivity assays were performed with Biolog GEN III microplates by using protocol C2 according to the instructions of the manufacturer (Biolog, Inc., Hayward, CA, USA).

**Text S5 -** Fatty acid analysis

The strains TH2 and OM4 were cultured on solid R2A medium (DSMZ medium 830) at 28 °C for three days. The cellular fatty acids were analyzed using the Microbial Identification System (MIDI; Sherlock version 6.1, TSBA40 method), according to instructions provided by the manufacturer [34]. A combined analysis by gas chromatography coupled to a mass spectrometer was used to confirm the identity of the fatty acids based on retention time and mass spectral data [35].

**Text S6 -** Plant-growth promotion assays

Re-inoculation tests were conducted to evaluate the ability of nodule formation of the bacterial isolates. Seed of the Sainfoin cultivars Taja and Perly, and sainfoin accession ONO20 were surface sterilized in 1 % ethanol for 1 min, followed by thrice rinses with sterilised distilled water (SDW), then transferred to 3% NaOCl for 5 min, and washed six times with SDW. Seeds were air-dried under laminar flow, and sown in 1 L pots containing a nitrogen free substrate (sterilised siliceous sand). Plants were grown in a greenhouse with the growth conditions of 16 h photoperiod, and 20 – 16°C day – night temperature. Each pot received four seeds. After germination, plants were reduced to one per pot. Pots were supplied twice a week with a modified nitrogen-free nutrient solution [36] consisted of $K_2HPO_4$ 0.5 mM, $CaCl_2$ 0.5 mM, $MgSO_4$ 0.25 mM, $K_2SO_4$ 0.5 mM, $Ca(H_2PO_4)_2$ 0.5mM, Fe-EDTA 0.06 mM, $MnSO_4$ 1µM, $H3BO_3$ 2 µM, $ZnSO_4$ 0.5 µM, $CuSO_4$ 0.2 µM, $CoSO_4$ 0.1 µM, $Na_2MoO_4$ 0.1 µM.

Bacterial inocula were prepared by growing bacterial strains of interest on YMA for up to 1 week. *Rhizobium leguminosarum* TS1-3-1, a strain frequently isolated from surface sterilized nodules of sainfoin during our broad scale screening (unpublished data) was used as a positive control for nodulation. The growing bacterial cultures were collected and suspended in 10 mM $MgCl_2$. The bacterial suspension was adjusted to $OD_{600}$ = 0.2. Each pot was inoculated with 10 ml of the corresponding bacterial strain upon sowing, one and two weeks after sowing. Controls received only 10 ml of 10 mM $MgCl_2$. Because strain TH2 did not have a nitrogenase cluster, a co-inoculation of *this* strain with *R. leguminosarum* TS1-3-1 was added for sainfoin cultivar Taja as the initial host of strain TH2. A total of 15 replicates were prepared for each bacterial strain of interest. Twelve weeks after inoculation, plants were harvested to determine the efficacy of the bacterial strains on nodulation of Sainfoin. Plant roots were washed with running tap water and the nodules formed were excised from the roots. Nodules were processed for bacterial isolation as described earlier. Bacterial colonies grown were subcultured on new plates. The growing cultures were processed for DNA-based identification. Genomic DNA of the cultures obtained from each treatment was extracted and processed to sequence the 16S rRNA as mentioned above. The obtained sequences from each bacterial treatment from the nodules of the given hosts (Taja, Perly, ONO20) were compared with the reference bacterial strains (the newly found strains) to determine their identity. In order to evaluate the effect of the different inocula on plant performance, above and below ground biomass was recorded. Statistical analyses were performed using R v. 3.6.1 (R Core Team 2018). Plant biomass of the nodulation assay was analysed using generalized linear models (GLM) with gamma family and log link for strictly positive continuous data.

**Text S7 -** Plasmid similarity

Plasmids of *M. onobrychidis* OM4$^T$ and *O. muellerharveyae* TH2$^T$ do not show high similarity to yet known mobile elements based on mash analysis. The outline neighbor network was computed based on mash distances for all plasmid sequences stored in the Refseq or PLS databases that shared at least 1 k-mer hash with the plasmids of *M. onobrychidis* OM4$^T$ or *O. muellerharveyae* TH2$^T$ (Fig. S6). Among the remaining 204 reference plasmid sequences, a maximum of 9 hashes could be identified as overlapping k-mers, when applying the mash tool, indicating a low similarity of all the novel plasmids (Table S1). The outline graph revealed the current sole branch of the *M. onobrychidis* OM4$^T$ plasmid related to *Mesorhizobium* spp. and the three *O. muellerharveyae* TH2$^T$ plasmids branching together between various genera. The very best hit to any of the three plasmids of *O. muellerharveyae* TH2$^T$ were in each case the other plasmids of the same strain. That might point to the origin of all three plasmids in one common ancestor of *O. muellerharveyae* TH2$^T$ and its precursor lineage, respectively. Thus, for both strains the plasmid neighbor network seems to foster the view of a species or genus of its own.

**Text S8 -** Functional comparison

On KEGG level 3, *M. onobrychidis* OM4$^T$ reveals several functions, *e. g.* enrichments in the glycolysis, pentosephosphate, fructose/mannose, vitamins, cell adhesion / invasion, carbon fixation in photosynthetic organism, bacterial secretion and *Escherichia coli*-like biofilm formation pathways (Fig. S11B). In contrast, *O. muellerharveyae* TH2$^T$ is more specialized in starch and sucrose metabolism, carotenoid biosynthesis, bacterial chemotaxis, xylene and tolulene degradation, carbon fixation in prokaryotic systems, biofilm formation of other bacteria, glucosinolate biosynthesis pathways.

**Text S9 -** Secondary metabolite biosynthesis gene clusters

*Mesorhizobium onobrychidis* OM4$^T$ encodes for 11 BGCs on its chromosome (Fig. S12 A), comprising types of indole, redox cofactor, T3PKS (surfactin), terpene, RiPP-like, siderophore, RRE-containing/lasspeptide, hserlactone (homoserine lactone) (2), proteusin, and phosphonate (9-methylstreptimidone). O*nobrychidicola muellerharveyae* TH2$^T$ only harbors 6 chromosomal and 1 plasmid-encoded BGCs, comprising the types thioamidites (2 on chromosome and 1 on plasmid 1), hserlactone, terpene, RRE-containing (galactoglucan), NRPS/T1PKS, and ectoine (Fig. S12 B).

**Text S10 -** Whole genome alignment

A whole genome alignment was performed for *M. onobrychidis* OM4$^T$, *M. delmotii* STM4623$^T$ and *M. temperatum* SDW018$^T$ to determine *M. onobrychidis* OM4$^T$-specific unique regions, which were not part of conserved collinear blocks (Fig. S13, Table S1). Here in total 135 of 486 genomic regions with minimal length of 2,000 bp and a maximum of 91,884 bp were detected. Among them, 85 regions harbored 5 to

77 genes. 21 regions showed affiliation to 7 of the 11 BGCs. Here, two BGCs were located in the genomic island (Fig. S14, Table S1). That island covered 63 unaligned regions including 364 genes, all assigned as unique genes. In total 628 of 1068 unique genes fall into these unaligned regions.

**Table S1 -** Genome Annotation of *Mesorhizobium onobrychidis* OM4$^T$ and *Onobrychidicola muellerharveyae* TH2$^T$ including pan genome, gene cluster and island information, KEGG functional protein annotations and their PGPT class associations, as well as their enrichments compared to other related strains. Beside the annotation files, the genomic circular visualisation files for BRIG, PGPT comparison file showing raw gene counts on selected classes, and the overlapping plasmid hits by mash analysis against plasmid databases are given.
(see Excel table: Table_S1 - SUPPL_GENOME_ANNOTATION.xlsx)

**Table S2 -** Whole-proteome average amino acid identity (wpAAI) comparisons between *Mesorhizobium onobrychidis* OM4$^T$, *Onobrychidicola muellerharveyae* TH2$^T$, and other members of the families *Rhizobiaceae* (various genera) and *Phyllobacteriaceae* (genera *Mesorhizobium* and *Nitratireductor*). The representatives of more distantly related genera *Aurantimonas*, *Aureimonas* and *Fulvimarina* were also included.
(see Excel table: Table_S2_AAI_matrix.xlsx)

**Table S3 –** Pairwise OGRI comparisons between *Mesorhizobium onobrychidis* OM4$^T$ and related *Mesorhizobium* spp.

| **Strain** | **ANIb** | **orthoANIu** | **dDDH** |
|---|---|---|---|
| *Mesorhizobium delmotii* STM4623$^T$ | 94,79 | 94,88 | 60,8 |
| *Mesorhizobium temperatum* SDW018$^T$ | 94,78 | 94,69 | 59,8 |
| *Mesorhizobium wenxiniae* WYCCWR 10195$^T$ | 93,02 | 92,98 | 51,4 |
| *Mesorhizobium prunaredense* STM4891$^T$ | 92,95 | 93,03 | 51,3 |
| *Mesorhizobium muleiense* CGMCC 1.11022$^T$ | 92,33 | 92,29 | 48,9 |
| *Mesorhizobium mediterraneum* USDA 3392$^T$ | 91,92 | 91,86 | 47,1 |

**Table S4** - Core-proteome average amino acid identity (cpAAI) comparisons between *Mesorhizobium onobrychidis* OM4$^T$, *Onobrychidicola muellerharveyae* TH2$^T$, and other members of the families *Rhizobiaceae* (various genera) and *Phyllobacteriaceae* (genera *Mesorhizobium* and *Nitratireductor*). The representatives of more distantly related genera *Aurantimonas*, *Aureimonas* and *Fulvimarina* were also included.

| | ***Onobrychidicola muellerharveyae* TH2$^T$** | | ***Onobrychidicola muellerharveyae* TH2$^T$** |
|---|---|---|---|
| ***Onobrychidicola muellerharveyae* TH2$^T$** | 100 | "*Rhizobium glycinendophyticum*" CL12$^T$ | 74,20135773 |
| *Shinella granuli* DSM 18401$^T$ | 75,37456195 | *Gellertiella hungarica* DSM 29853$^T$ | 74,1996648 |
| *Pararhizobium polonicum* F5.1$^T$ | 75,31530921 | *Pseudorhizobium marinum* R1-200B4$^T$ | 74,19627893 |
| *Shinella zoogloeoides* PQ7$^T$ | 75,31530921 | "*Peteryoungia rhizophila*" 7209-2$^T$ | 74,18104251 |
| *Rhizobium leguminosarum* USDA 2370$^T$ | 75,31192334 | *Rhizobium populi* CCTCC AB 2013068$^T$ | 74,12348271 |
| *Ensifer sesbaniae* CCBAU 65729$^T$ | 75,27975757 | *Rhizobium tarimense* CCTCC AB 2011011$^T$ | 74,12178977 |
| *Rhizobium grahamii* CCGE 502$^T$ | 75,25097767 | *Pseudorhizobium banfieldii* NT-26$^T$ | 74,12009684 |
| *Pararhizobium giardinii* H152$^T$ | 75,24589886 | "*Peteryoungia wuzhouensis*" W44$^T$ | 74,10147455 |
| *Ensifer adhaerens* Casida A$^T$ | 75,20188254 | *Pseudorhizobium flavum* YW14$^T$ | 74,04560768 |
| *Ensifer morelensis* Lc04$^T$ | 75,18156732 | *Allorhizobium pseudoryzae* DSM 19479$^T$ | 74,04222181 |
| *Sinorhizobium terangae* SEMIA 6460$^T$ | 75,15109448 | *Rhizobium tumorigenes* 1078$^T$ | 73,99651255 |
| *Ensifer mexicanus* ITTG R7$^T$ | 75,14940155 | "*Allorhizobium paknamense*" DSM 100301$^T$ | 73,98804788 |
| *Rhizobium rhizogenes* ATCC 11325$^T$ | 75,11554284 | "*Neorhizobium lilium*" 24NR$^T$ | 73,90340111 |
| *Sinorhizobium saheli* LMG 7837$^T$ | 75,09014881 | *Agrobacterium radiobacter* CFBP 5522$^T$ | 73,89493643 |
| *Ensifer glycinis* CCBAU 23380$^T$ | 75,08845587 | "*Peteryoungia ipomoeae*" shin9-1$^T$ | 73,86446359 |
| *Ensifer psoraleae* CCBAU 65732$^T$ | 75,06475478 | *Rhizobium smilacinae* CCTCC AB 2013016$^T$ | 73,84414837 |
| *Sinorhizobium americanum* CFNEI 156$^T$ | 75,05967597 | *Neorhizobium* sp. NCHU2750 | 73,7882815 |
| *Rhizobium tropici* CIAT 899$^T$ | 75,05459717 | *Allorhizobium undicola* ORS 992$^T$ | 73,76627334 |
| "*Rhizobium album*" NS-104$^T$ | 75,03428194 | *Agrobacterium larrymoorei* AF3.10$^T$ | 73,76119454 |
| *Pararhizobium arenae* MIM27$^T$ | 75,02920314 | *Rhizobium* sp. NFR03 | 73,74934399 |
| *Ensifer shofinae* CCBAU 251167$^T$ | 74,92762701 | *Rhizobium cellulosilyticum* DSM 18291$^T$ | 73,73918638 |
| *Pararhizobium antarcticum* NAQVI 59$^T$ | 74,90900472 | *Rhizobium* sp. 9140 | 73,73241463 |
| *Neorhizobium alkalisoli* DSM 21826$^T$ | 74,86668134 | *Agrobacterium rubi* TR3$^T$ | 73,66131135 |
| *Rhizobium selenitireducens* ATCC BAA-1503$^T$ | 74,84975198 | *Rhizobium* sp. Leaf371 | 73,61221622 |
| *Sinorhizobium fredii* USDA 205$^T$ | 74,84298024 | *Endobacterium cereale* RZME27$^T$ | 73,57497164 |
| *Mycoplana dimorpha* DSM 7138 | 74,84128731 | *Rhizobium yantingense* CCTCC AB 2014007$^T$ | 73,57158577 |
| *Neorhizobium vignae* CCBAU 05176$^T$ | 74,8362085 | *Rhizobium borbori* LMG 23925$^T$ | 73,55634935 |
| *Neorhizobium huautlense* DSM 21817$^T$ | 74,81927915 | *Rhizobium helianthi* CGMCC 1.12192$^T$ | 73,486939 |
| *Ensifer sojae* CCBAU 05684$^T$ | 74,76171935 | *Rhizobium rhizoryzae* DSM 29514$^T$ | 73,46323791 |
| *Sinorhizobium meliloti* USDA1002$^T$ | 74,75664054 | *Rhizobium* sp. Leaf383 | 73,43107214 |
| *Neorhizobium galegae* HAMBI 540$^T$ | 74,73463238 | *Allorhizobium vitis* K309$^T$ | 73,41244985 |
| *Rhizobium petrolearium* DSM 26482$^T$ | 74,71939596 | *Allorhizobium taibaishanense* 14971$^T$ | 73,39382756 |
| *Neorhizobium tomejilense* T17$^T$ | 74,68723019 | *Rhizobium halophytocola* DSM 21600$^T$ | 73,37012646 |
| "*Mycoplana subbaraonis*" JC85$^T$ | 74,68045845 | *Allorhizobium terrae* CC-HIH110$^T$ | 73,26008566 |
| *Rhizobium azooxidifex* DSM 100211$^T$ | 74,6686079 | *Allorhizobium oryziradicis* N19$^T$ | 73,18898238 |
| *Sinorhizobium arboris* LMG 14919$^T$ | 74,66352909 | *Rhizobium rhizosphaerae* MH17$^T$ | 73,03153939 |
| *Sinorhizobium kostiense* DSM 13372$^T$ | 74,60258342 | *Rhizobium oryzae* CGMCC 1.7048$^T$ | 72,89441162 |
| *Rhizobium pakistanense* CCBAU 101086$^T$ | 74,58226819 | *Hoeflea olei* JC234$^T$ | 70,13661989 |
| *Ensifer alkalisoli* YIC4027$^T$ | 74,56533884 | *Hoeflea marina* DSM 16791$^T$ | 69,83866326 |
| *Rhizobium daejeonense* DSM 17795$^T$ | 74,55518123 | *Hoeflea phototrophica* DFL-43$^T$ | 69,77941052 |
| *Ciceribacter lividus* DSM 25528$^T$ | 74,51285784 | *Martelella endophytica* YC6887$^T$ | 69,66936972 |
| *Sinorhizobium medicae* A321$^T$ | 74,50777904 | *Martelella mediterranea* DSM 17316$^T$ | 69,51192673 |
| "*Ciceribacter ferrooxidans*" F8825$^T$ | 74,5060861 | *Hoeflea halophila* KCTC 23107$^T$ | 69,4628316 |
| *Ciceribacter thiooxidans* F43B$^T$ | 74,47053446 | *Georhizobium profundi* WS11$^T$ | 69,37987777 |
| "*Peteryoungia rosettiformans*" W3$^T$ | 74,36557247 | "*Neopararhizobium haloflavum*" XC0140$^T$ | 69,37479896 |
| "*Agrobacterium albertimagni*" AOL15$^T$ | 74,30124092 | *Martelella mediterranea* USBA-857 | 69,21227717 |
| *Rhizobium naphthalenivorans* TSY03b$^T$ | 74,26061047 | *Mesorhizobium huakuii* 7653R | 67,38390696 |
| *Rhizobium aggregatum* DSM 1111$^T$ | 74,25383873 | *Mesorhizobium ciceri* WSM1271$^T$ | 67,251858 |
| *Pseudorhizobium marinum* MGL06$^T$ | 74,2521458 | *Mesorhizobium loti* DSM 2626$^T$ | 67,10626555 |

**Table S5 -** Differential characteristics of *Onobrychidicola muellerharveyae* TH2$^{T}$ and the type species from the other genera of family *Rhizobiaceae.*

| | ***Onobrychidicola muellerharveyae* TH2$^{T}$** | ***Agrobacterium radiobacter* CFBP 5522$^{T}$** | ***Allorhizobium undicola* ORS 992$^{T}$** | ***Ciceribacter lividus* DSM 25528$^{T}$** | ***Ensifer adhaerens* Casida A$^{T}$** | ***Gellertiella hungarica* DSM 29853$^{T}$** | ***Mycoplana dimorpha* DSM 7138$^{T}$** | ***Neorhizobium galegae* HAMBI 540$^{T}$** | ***Pseudorhizobium marinum* MGL06$^{T}$** | ***Pararhizobium giardinii* H152$^{T}$** | ***Rhizobium leguminosarum* USDA 2370$^{T}$** | ***Shinella granuli* DSM 18401$^{T}$** |
|---|---|---|---|---|---|---|---|---|---|---|---|---|
| **Isolation source** | *Onobrychis viciifolia* nodules | ND | *Neptunia natans* nodules | Rhizosphere soil of *Cicer arietinum* | Soil | Pool water of a thermal bath | Soil | *Galega orientalis* nodules | Surface seawater | *Phaseolus vulgaris* nodules | *Pisum sativum* nodules | Upflow anaerobic sludge blanket reactor |
| **Colony color (on YMA)** | White | White-cream[1] | White-cream[1] | Bluish black[1] | White[1] | ND | ND | White-pink[1] | ND[20] | White-cream[1] | White-cream[1] | Ivory[1] |
| **Flagellation** | - | 1-4 peritrichous[2] | ND[7] | ND[10] | 3-5 subpolar flagella[12] | Polar, monotrichous[15] | Peritrichous[17] | 1-2 polar or subpolar flagella[18] | ≥3 flagella[21] | ND | 1 or 2 polar flagella or 2–6 peritrichous flagella[24] | Multiple polar flagella[27] |
| **Growth temperature range (°C)** | 10-30 | 15-37[3] | ND | 25-37[11] | 20-37[1] | 15-45[15] | 4-45[15] | ND | 20-35[21] | 4-37[3] | 10-35[25] | 4-40[27] |
| **Growth at 37°C** | - | +[1] | +[1] | +[1,11] | +[1,13] | +[15] | +[15] | +[1] | -[21] | w[1] | -[1,25] | +[1,27] |
| **Reduction of nitrate** | - | +[1] | -[1] | -[1] | +[1,13,14] | -[15] | v[17] | -[1,18,19] | +[19,21] | -[1] | -[1,25] | +[1,27] |
| **Enzyme activity:** | | | | | | | | | | | | |
| **Urease** | + | +[4] | ND | +[11] | +[14] | -[15] | +[17] | +[18,19] | +[19] or -[21] | -[3] or +[23] | +[25] | +[27] |
| **β-galactosidase** | w | +[4] | ND | ND | +[14] | -[15] | ND | +[19] | +[19,21] | -[23] | +[25] | +[27] |
| **Hydrolysis of:** | | | | | | | | | | | | |
| **Aesculin** | + | +[4] | -[8] | ND | +[14] | -[15] | -[15] | +[19] | +[19,21] | ND | +[15] | +[28] |
| **Gelatin** | - | -[4] | ND | -[11] | -[14] | +[15] | -[15,17] | -[19] | -[19,21] | ND | -[15,25] | -[27] |
| **Assimilation of:** | | | | | | | | | | | | |
| **Gluconate** | w | +[1,4] | -[1] | -[1] | -[1,14] | -[16] | ND | +[1,19] | +[19,21] | -[1] | +[1] | +[1,27] |
| **Malate** | - | +[1,4] | +[1] | +[1] | +[1,14] | ND | ND | +[1,19] | - [19,21] | +[1] | w[1] | +[1,27] |
| **Major fatty acids (>5 %)** | $C_{18:1}$ w7c, $C_{19:0}$ CYCLO w7c, $C_{16:0}$ and $C_{17:0}$ CYCLO w7c | $C_{18:1}$ w7c, $C_{14:0}$ 3OH/$C_{16:1}$ iso I/$C_{12:0}$ aldehyde, $C_{16:0}$[5] | $C_{18:1}$ w7c/ $C_{18:1}$ w6c, $C_{14:0}$ 3OH/ $C_{16:1}$ iso I/ $C_{12:0}$ aldehyde[1] | $C_{18:1}$ w7c, $C_{19:0}$ CYCLO w8c, $C_{18:0}$[11] | $C_{18:1}$ w7c/ $C_{18:1}$ w6c, $C_{14:0}$ 3OH/ $C_{16:1}$ iso I/ $C_{12:0}$ aldehyde[1] | $C_{18:1}$ w7c, 11-Me $C_{18:1}$ w7c[15] | $C_{18:1}$ w7c, $C_{19:0}$ CYCLO w8c, $C_{17:0}$[15] | $C_{18:1}$ w7c/ $C_{18:1}$ w6c, $C_{16:0}$, $C_{19:0}$ cyclo w8c[1] | $C_{18:1}$ w7c/ $C_{18:1}$ w6c, $C_{19:0}$ cyclo w8c, $C_{16:0}$, $C_{14:0}$ 3OH/ $C_{16:1}$ iso I/ $C_{12:0}$ aldehyde[21] | $C_{18:1}$ w7c/ $C_{18:1}$ w6c, $C_{14:0}$ 3OH/ $C_{16:1}$ iso I/ $C_{12:0}$ aldehyde, $C_{16:1}$ w7c/ $C_{16:1}$ w6c, $C_{16:0}$[1] | $C_{18:1}$ w7c/ $C_{18:1}$ w6c, $C_{18:0}$, $C_{16:0}$, $C_{18:1}$ w7c 11-methyl[1] | $C_{18:1}$ w7c/ $C_{18:1}$ w6c, $C_{19:0}$ cyclo w8c, $C_{16:0}$, $C_{18:1}$ w7c 11-methyl[1] |
| **Reference genome sequence** | CP062231-CP062234 | LMVJ01 | JHXQ01 | QPIX01 | CP015880.1-CP015882.1 | JACIEZ01 | PZZZ010 | HG938353.1, HG938354.1 | JMQK01 | KB902578-KB902767 | MRDL01 | SLVX01 |
| **Genome size (Mbp)** | 6.44 | 5.50 | 4.13 | 4.52 | 7.27 | 4.77 | 4.59 | 6.46 | 5.06 | 6.81 | 7.81 | 6.59 |
| **Genome architecture** | 1 circular chromosome + plasmids | ND[6] | ND[9] | ND | Circular chromosome + two additional large replicons (chromids and/or megaplasmids) | ND | ND | Circular chromosome + circular chromid | ND[22] | ND | ND[26] | ND |
| **Genome G+C Content (%)** | 60.4 | 59.4 | 59.3 | 63.2 | 62.3 | 63.5 | 64.5 | 61.1 | 63.0 | 60.7 | 60.6 | 65.0 |

+, positive; -, negative; w, weak reaction; v, variable; ND, not determined or not available.

[1] Data from Kimes et al. (2014), http://dx.doi.org/10.1016/j.syapm.2015.05.003; [2] Data from Allen and Holding (1974) - Allen ON, Holding AJ. Genus II. Agrobacterium Conn 1942, 359; Nom. gen. cons. Opin. 33, Jud. Comm. 1970, 10. In: Buchanan RE, Gibbons NE (eds), Bergey's Manual of Determinative Bacteriology, Eighth Edition, The Williams and Wilkins Co., Baltimore, 1974, p. 264-267.; [3] Data from Bibi et al. (2012), 10.1099/ijs.0.029488-0; [4] Data from Leibniz Institut DSMZ-Deutsche Sammlung von Mikroorganismen und Zellkulturen GmbH ; Curators of the DSMZ; DSM 30147; [5] Data from Puławska et al. (2021), 10.1099/ijs.0.032532-0; [6] It was reported that the genome organization of the genus *Agrobacterium* is characterized by the presence of a circular chromosome and a secondary linear chromid (Ramírez-Bahena et al., 2014, 10.1016/j.ympev.2014.01.005; Harrison et al., 2010, 10.1016/j.tim.2009.12.010; Slater et al., 2009, https://doi.org/10.1128/JB.01779-08); [7] Lin et al. (2020), DOI 10.1099/ijsem.0.003770, reported polar flagellation for a species *Allorhizobium terrae* belonging to the same genus; [8] Data from de Lajudie et al. (1998), https://doi.org/10.1099/00207713-48-4-1277; [9] Genome comprising circular chromosome, circular chromid and multiple plasmids was reported for species *Allorhizobium vitis* (strain S4) belonging to the same genus (Harrison et al., 2010, 10.1016/j.tim.2009.12.010; Slater et al., 2009, https://doi.org/10.1128/JB.01779-08); [10] Deng et al. (2017, 2020), https://doi.org/10.1007/s12275-020-9471-2, 10.1099/ijsem.0.002367, reported a single flagellum for two *Ciceribacter* species. [11] Data from Kathiravan et al. (2013), 10.1099/ijs.0.049726-0; [12] Data from Balkwill (2005), "Balkwill DL (2005). "Ensifer Casida 1982, 343VP". In Brenner DJ, Krieg NR, Garrity GM, Staley JT, Boone DR, De Vos P, Goodfellow M, Rainey FA, Schleifer KH (eds.). Bergey's Manual of Systematic Bacteriology, Volume Two: The Proteobacteria, Part C: The Alpha-, Beta-, Delta-, and Epsilonproteobacteria. New York, New York: Springer. pp. 354–361"; [13] Data from Casida (1982), https://doi.org/10.1099/00207713-32-3-339; [14] Data from Willems et al. (2003), https://doi.org/10.1099/ijs.0.02264-0; [15] Data from Tóth et al. (2017), 10.1099/ijsem.0.002332; [16] Data from Leibniz Institut DSMZ-Deutsche Sammlung von Mikroorganismen und Zellkulturen GmbH ; Curators of the DSMZ; DSM 29853; [17] Data from Urakami et al. (1990), https://doi.org/10.1099/00207713-40-4-434; [18] Data from Lindström (1989), https://doi.org/10.1099/00207713-39-3-365; [19] Lassalle et al. (2020), https://doi.org/10.1016/j.syapm.2020.126165; [20] *Pseudorhizobium* strain R1-200B4, which belongs to the same species as *P. marinum* (Lasalle et al., 2020, https://doi.org/10.1016/j.syapm.2020.126165), forms white colonies on YMA medium (Kimes et al., 2014, http://dx.doi.org/10.1016/j.syapm.2015.05.003); [21] Data from Liu et al. (2015), 10.1099/ijsem.0.000593; [22] Other Pseudorhizobium species carries circular chromosome and variable number of plasmids (Lasalle et al., 2020, https://doi.org/10.1016/j.syapm.2020.126165); [23] Naqvi et al. (2017), https://doi.org/10.1099/ijsem.0.001828; [24] Data from Kuykendall et al. (2005), "Kuykendall LD (2005) Family I. *Rhizobiaceae* Conn 1938, 321[AL]. In: Brenner DJ, Krieg NR, Stanley JT (eds) Bergey's manual of systematic bacteriology, vol 2. Springer, New York, pp 324–361"; [25] Data from Ramírez-Bahena et al. (2008), 10.1099/ijs.0.65621-0; [26] Rhizobium leguminosarum complex genomes typically have circular chromosome, chromids and plasmids (Young et al., 2021, https://doi.org/10.3390/genes12010111); [27] Data from An et al. (2006), 10.1099/ijs.0.63942-0; [28] Data from Subhash and Lee (2016), https://doi.org/10.1099/ijsem.0.001290.

**Table S6 -** Phenotypic characteristics of *Mesorhizobium onobrychidis* OM4$^T$ and *Onobrychidicola muellerharveyae* TH2$^T$.

| **Test** | ***Onobrychidicola muellerharveyae* sp. nov.** | ***Mesorhizobium onobrychidis* sp. nov.** |
|---|---|---|
| | **TH2$^T$** | **OM4$^T$** |
| Flagellation | - | - |
| Motility | - | - |
| KOH test | + | + |
| Aminopeptidase test | + | + |
| Catalase test | + | + |
| Oxidase test | + | + |
| Growth temperature range (°C) | 10-30 | 10-25 |
| Growth at 37°C | - | - |
| Nitrate reduction[1] | - | - |
| Indole production[1] | n.g. | n.g. |
| Glucose fermentation[1] | - | - |
| Arginine dihydrolase production[1] | - | - |
| Urease production[1] | + | + |
| Aesculin hydrolysis[1] | + | n.g. |
| Gelatin hydrolysis[1] | - | n.g. |
| β-galactosidase test[2] | w | - |
| D-glucose assimilation[1] | + | + |
| L-arabinose assimilation[1] | w | + |
| D-mannose assimilation[1] | + | + |
| D-mannitol assimilation[1] | w | + |
| D-maltose assimilation[1] | w | + |
| Potassium gluconate assimilation[1] | w | + |
| Capric acid assimilation[1] | w | w |
| Adipic acid assimilation[1] | - | - |
| Malic acid assimilation[1] | - | + |
| Trisodium citrate assimilation[1] | - | - |
| Phenylacetic acid assimilation[1] | - | - |

+, positive; -, negative; w, weak reaction; n.g., no bacterial growth observed
[1] Results obtained with conventional biochemical tests.
[2] Results obtained with API 20NE system.

**Table S7 -** Cellular fatty acid composition of *Mesorhizobium onobrychidis* OM4$^T$ and *Onobrychidicola muellerharveyae* TH2$^T$.

| Fatty Acid[1] | ***Onobrychidicola muellerharveyae* sp. nov.** | ***Mesorhizobium onobrychidis* sp. nov.** |
|---|---|---|
| | **TH2$^T$** | **OM4$^T$** |
| $C_{18:1}$ w7c | 48.64 | 60.21 |
| $C_{19:0}$ CYCLO w7c | 23.05 | 7.65 |
| $C_{16:0}$ | 12.96 | 12.8 |
| $C_{17:0}$ CYCLO w7c | 5.36 | - |
| $C_{14:0}$ 3OH | 2.26 | - |
| $C_{18:0}$ | 1.57 | 5.23 |
| $C_{16:1}$ w7c | 1.35 | 1.01 |
| 11 methyl $C_{18:1}$ w7c | 1.34 | 6.42 |
| $C_{17:0}$ ISO | - | 2.87 |
| $C_{18:1}$ w9c | - | 1.04 |

[1] Fatty acids present in amounts lower than 1% are not shown.

**Table S8 -** List of strains and GenBank/EMBL/DDBJ accession numbers for their nucleotide sequences used in this study

| No. | Strain | Accession Number | No. | Strain | Accession Number |
|---|---|---|---|---|---|
| 1 | "*Agrobacterium fabrum*" C58 | AE007869.2-AE007872.2 | 51 | *Mesorhizobium qingshengii* CGMCC 1.12097$^T$ | FMXM00000000.1 |
| 2 | *Agrobacterium larrymoorei* AF3.10$^T$ | JADW00000000.1 | 52 | *Mesorhizobium sanjuanii* BSA136$^T$ | NWQG00000000.1 |
| 3 | *Agrobacterium radiobacter* B6$^T$ | FCNL00000000.1 | 53 | *Mesorhizobium sophorae* ICMP 19535$^T$ | NNRI00000000.1 |
| 4 | *Agrobacterium radiobacter* CFBP 5522$^T$ | LMVJ00000000.1 | 54 | *Mesorhizobium temperatum* SDW018$^T$ | NPKJ00000000.1 |
| 5 | *Agrobacterium rosae* NCPPB 1650$^T$ | NXEJ00000000.1 | 55 | *Mesorhizobium waimense* ICMP19557$^T$ | QZWZ00000000.1 |
| 6 | *Agrobacterium rubi* TR3$^T$ | BBJU00000000.1 | 56 | *Mesorhizobium wenxiniae* WYCCWR 10195$^T$ | NPKH00000000.1 |
| 7 | *Allorhizobium oryziradicis* N19$^T$ | MKIM00000000.1 | 57 | *Mycoplana dimorpha* DSM 7138$^T$ | PZZZ00000000.1 |
| 8 | *Allorhizobium taibaishanense* 14971$^T$ | MKIN00000000.1 | 58 | "*Mycoplana subbaraonis*" JC85$^T$ | OBQD00000000.1 |
| 9 | *Allorhizobium terrae* CC-HIH110$^T$ | SSOA00000000.1 | 59 | "*Neopararhizobium haloflavum*" XC0140$^T$ | NWUC00000000.1 |
| 10 | *Allorhizobium undicola* ORS 992$^T$ | JHXQ00000000.1 | 60 | *Neorhizobium alkalisoli* DSM 21826$^T$ | PVBN00000000.1 |
| 11 | *Allorhizobium vitis* K309$^T$ | LMVL00000000.2 | 61 | *Neorhizobium galegae* HAMBI 540$^T$ | HG938353.1; HG938354.1 |
| 12 | *Allorhizobium vitis* S4 | CP000633.1-CP000639.1 | 62 | *Neorhizobium huautlense* DSM 21817$^T$ | PVBM00000000.1 |
| 13 | *Aurantimonas coralicida* DSM 14790$^T$ | ATXK00000000.1 | 63 | *Neorhizobium* sp. NCHU2750 | CP030827.1-CP030833.1 |
| 14 | *Aurantimonas manganoxydans* SI85-9A1$^T$ | AAPJ00000000.1 | 64 | *Neorhizobium vignae* CCBAU 05176$^T$ | JNNU00000000.1 |
| 15 | *Aureimonas altamirensis* DSM 21988$^T$ | BBWQ00000000.1 | 65 | *Nitratireductor aquibiodomus* JCM 21793$^T$ | BAMP00000000.1 |
| 16 | *Aureimonas ureilytica* DSM 18598$^T$ | ARQE00000000.1 | 66 | *Nitratireductor indicus* C115$^T$ | AMSI00000000.1 |
| 17 | *Ciceribacter lividus* DSM 25528$^T$ | QPIX00000000.1 | 67 | *Nitratireductor pacificus* pht-3B$^T$ | AMRM00000000.1 |
| 18 | "*Ciceribacter ferrooxidans*" F8825$^T$ | SDVB00000000.1 | 68 | *Pararhizobium antarcticum* NAQVI 59$^T$ | LSRP00000000.1 |
| 19 | *Ensifer adhaerens* Casida A$^T$ | CP015880.1; CP015881.1; CP015882.1 | 69 | *Pararhizobium giardinii* H152$^T$ | ARBG00000000.1 |
| 20 | *Ensifer fredii* HH103 | HE616890.1-HE616893.1; HE616899.1 | 70 | *Pararhizobium polonicum* F5.1$^T$ | LGLV00000000.1 |
| 21 | *Ensifer meliloti* 1021 | AL591688.1; AE006469.1; AL591985.1 | 71 | *Pseudorhizobium banfieldii* NT-26$^T$ | FO082820.1-FO082822.1 |
| 22 | *Ensifer sojae* CCBAU 05684$^T$ | AJQT00000000.1 | 72 | *Pseudorhizobium flavum* YW14$^T$ | MUXO00000000.1 |
| 23 | *Fulvimarina manganoxydans* CGMCC 1.10972$^T$ | FWXR00000000.1 | 73 | *Pseudorhizobium marinum* MGL06$^T$ | JMQK00000000.1 |
| 24 | *Fulvimarina pelagi* HTCC2506$^T$ | AATP00000000.1 | 74 | *Pseudorhizobium pelagicum* R1-200B4$^T$ | JOKI00000000.1 |
| 25 | *Gellertiella hungarica* DSM 29853$^T$ | Ga0373349 (IMG) | 75 | *Rhizobium aggregatum* LMG 23059$^T$ | NR |
| 26 | *Georhizobium profundi* WS11$^T$ | CP032509.1 | 76 | "*Agrobacterium albertimagni*" AOL15$^T$ | ALJF00000000.1 |
| 27 | *Hoeflea halophila* KCTC 23107$^T$ | OCPC00000000.1 | 77 | *Rhizobium arenae* MIM27$^T$ | MOOY00000000.1 |
| 28 | *Hoeflea marina* DSM 16791$^T$ | QGTR00000000.1 | 78 | *Rhizobium azooxidifex* DSM 100211$^T$ | JACIEE000000000.1 |
| 29 | *Hoeflea olei* JC234$^T$ | LQZT00000000.1 | 79 | *Rhizobium daejeonense* DSM 17795$^T$ | Ga0196644 (IMG) |
| 30 | *Hoeflea phototrophica* DFL-43$^T$ | ABIA03000001.1 | 80 | *Rhizobium etli* CFN 42$^T$ | CP000133.1-CP000138.1; U80928.5 |
| 31 | *Martelella endophytica* YC6887$^T$ | CP010803.1 | 81 | *Rhizobium freirei* PRF 81$^T$ | AQHN00000000.1 |
| 32 | *Martelella mediterranea* DSM 17316$^T$ | AQWH00000000.1 | 82 | *Rhizobium gallicum* R602$^T$ | CP006877.1-CP006880.1 |
| 33 | *Martelella mediterranea* USBA-857 | PVUG00000000.1 | 83 | *Rhizobium grahamii* CCGE 502$^T$ | AEYE00000000.2 |
| 34 | *Mesorhizobium australicum* WSM2073$^T$ | CP003358.1 | 84 | "*Peteryoungia ipomoeae*" shin9-1$^T$ | STGV00000000.1 |
| 35 | *Mesorhizobium ciceri* WSM1271$^T$ | CP002447.1; CP002448.1 | 85 | *Rhizobium leguminosarum* USDA 2370$^T$ | MRDL00000000.1 |
| 36 | *Mesorhizobium delmotii* STM4623$^T$ | FUIG00000000.1 | 86 | *Rhizobium oryzae* CGMCC 1.7048$^T$ | FOAM00000000.1 |
| 37 | *Mesorhizobium erdmanii* USDA 3471$^T$ | AXAE00000000.1 | 87 | *Rhizobium rhizogenes* ATCC 11325$^T$ | BAYX00000000.1 |
| 38 | *Mesorhizobium hawassense* AC99b$^T$ | QMBP00000000.1 | 88 | *Rhizobium rhizogenes* K84 | CP000628.1-CP000632.1 |
| 39 | *Mesorhizobium helmanticense* CSLC115N$^T$ | PZJX00000000.1 | 89 | *Rhizobium rhizosphaerae* MH17$^T$ | MKIO00000000.1 |
| 40 | *Mesorhizobium huakuii* 7653R | CP006581.1-CP006583.1 | 90 | "*Peteryoungia rosettiformans*" W3$^T$ | STGU00000000.1 |
| 41 | *Mesorhizobium intechi* BD68$^T$ | PNOT00000000.2 | 91 | *Rhizobium selenitireducens* ATCC BAA-1503$^T$ | JAEG00000000.1 |
| 42 | *Mesorhizobium japonicum* MAFF 303099$^T$ | BA000012.4, BA000013.4, AP003017.1 | 92 | *Rhizobium tropici* CIAT 899$^T$ | CP004015.1-CP004018.1 |
| 43 | *Mesorhizobium jarvisii* LMG 28313$^T$ | QZXA00000000.1 | 93 | *Rhizobium tubonense* CCBAU 85046$^T$ | PCDP00000000.1 |
| 44 | *Mesorhizobium loti* DSM 2626$^T$ | QGGH00000000.1 | 94 | *Rhizobium tumorigenes* 1078$^T$ | PCDQ00000000.1 |
| 45 | *Mesorhizobium mediterraneum* USDA 3392$^T$ | NPKI00000000.1 | 95 | "*Peteryoungia wuzhouensis*" W44$^T$ | QJRY00000000.1 |
| 46 | *Mesorhizobium metallidurans* STM 2683$^T$ | CAUM00000000.1 | 96 | *Shinella granuli* DSM 18401$^T$ | SLVX00000000.1 |
| 47 | *Mesorhizobium muleiense* CGMCC 1.11022$^T$ | FNEE00000000.1 | 97 | *Shinella* sp. DD12 | AYLZ00000000.2 |
| 48 | *Mesorhizobium opportunistum* WSM2075$^T$ | CP002279.1 | 98 | *Shinella* sp. WS1-2 | QWBU00000000.1 |
| 49 | *Mesorhizobium plurifarium* ORS1032$^T$ | CCND00000000.1 | 99 | *Shinella zoogloeoides* PQ7 | QNUJ00000000.1 |
| 50 | *Mesorhizobium prunaredense* STM4891$^T$ | FTPD00000000.1 | | | |

NR, not released

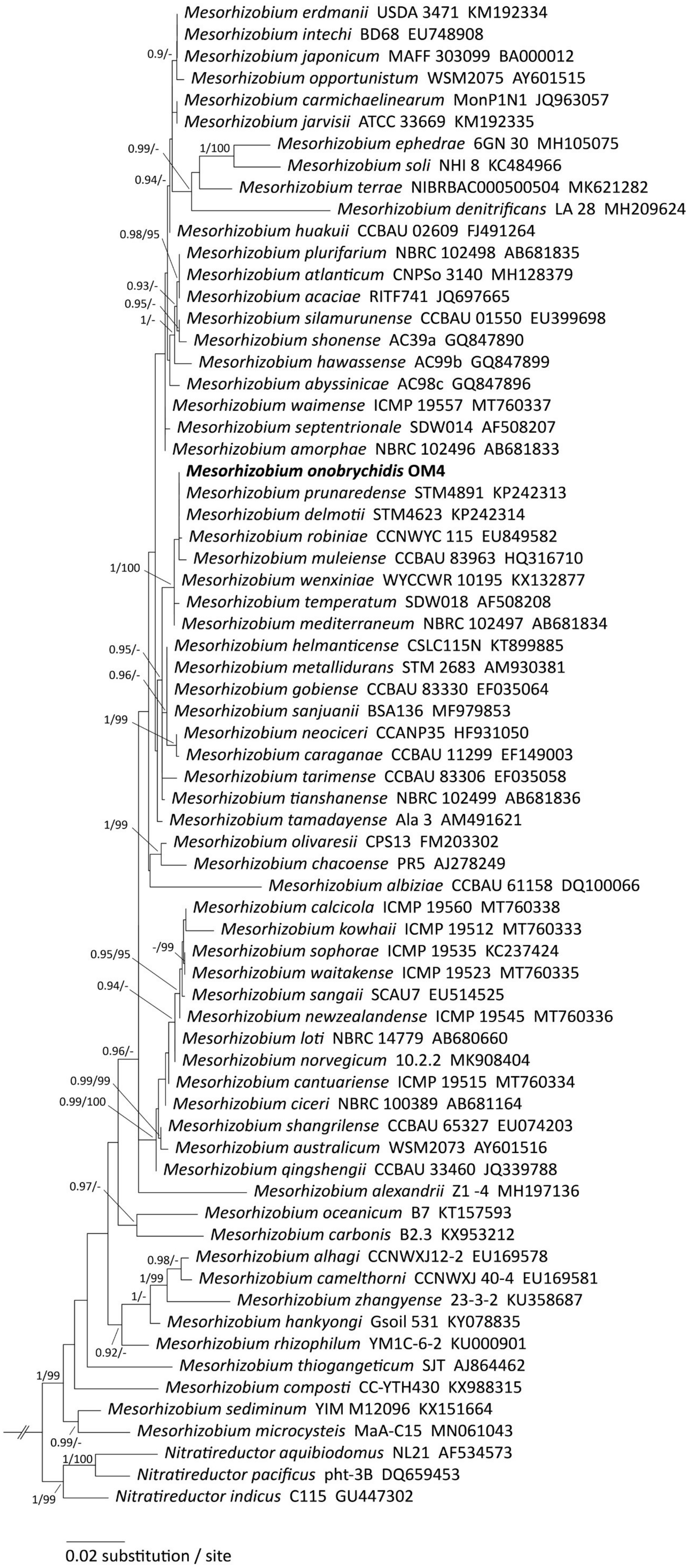


**Figures S1 –** A maximum likelihood topology estimated by the IQ-TREE analysis of 16S rRNA gene partial sequences of taxa of *Phyllobacteriaceae* using TIM2+F+I+G4 as best-fit substitution model. Alignment contains 69 sequences with 1422 nucleotide sites, 281 distinct patterns, 131 parsimony-informative, 109 singleton sites, and 1182 constant sites. Support values of Bayesian posterior probabilities ≥90 % and ultrafast bootstrap ≥95 % are given above each node, respectively. The new species is highlighted in bold. The tree is rooted by representatives of *Nitratireductor*.

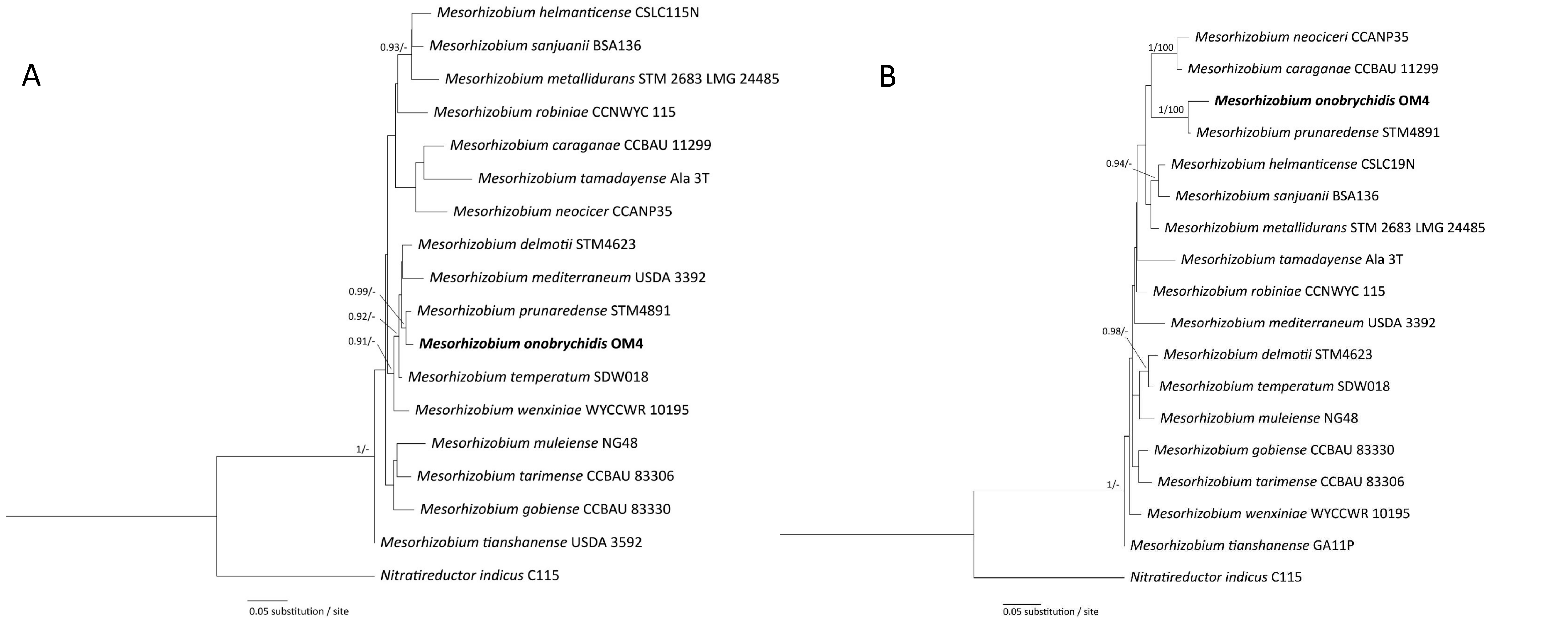


**Figures S2 –** *recA* and *atpD* tree *Mesorhizobium onobrychidis* OM4$^T$. **A.** A maximum likelihood topology estimated by the IQ-TREE analysis of *recA* gene partial sequences of taxa of *Rhizobiaceae* using TIM2+F+G4 as best-fit substitution model. Alignment contains 18 sequences with 566 nucleotide sites, 183 distinct patterns, 61 parsimony-informative, 78 singleton sites, and 427 constant sites. **B.** A maximum likelihood topology estimated by the IQ-TREE analysis of *atpD* gene partial sequences of taxa of *Rhizobiaceae* using TN+F+I+G4 as best-fit substitution model. Alignment contains 18 sequences with 513 nucleotide sites, 157 distinct patterns, 61 parsimony-informative, 71 singleton sites, and 381 constant sites. Support values of Bayesian posterior probabilities ≥90 % and ultrafast bootstrap ≥95 % are given above each node, respectively. The new species is highlighted in bold. Both trees are rooted by representatives of *Nitratireductor indicus*.

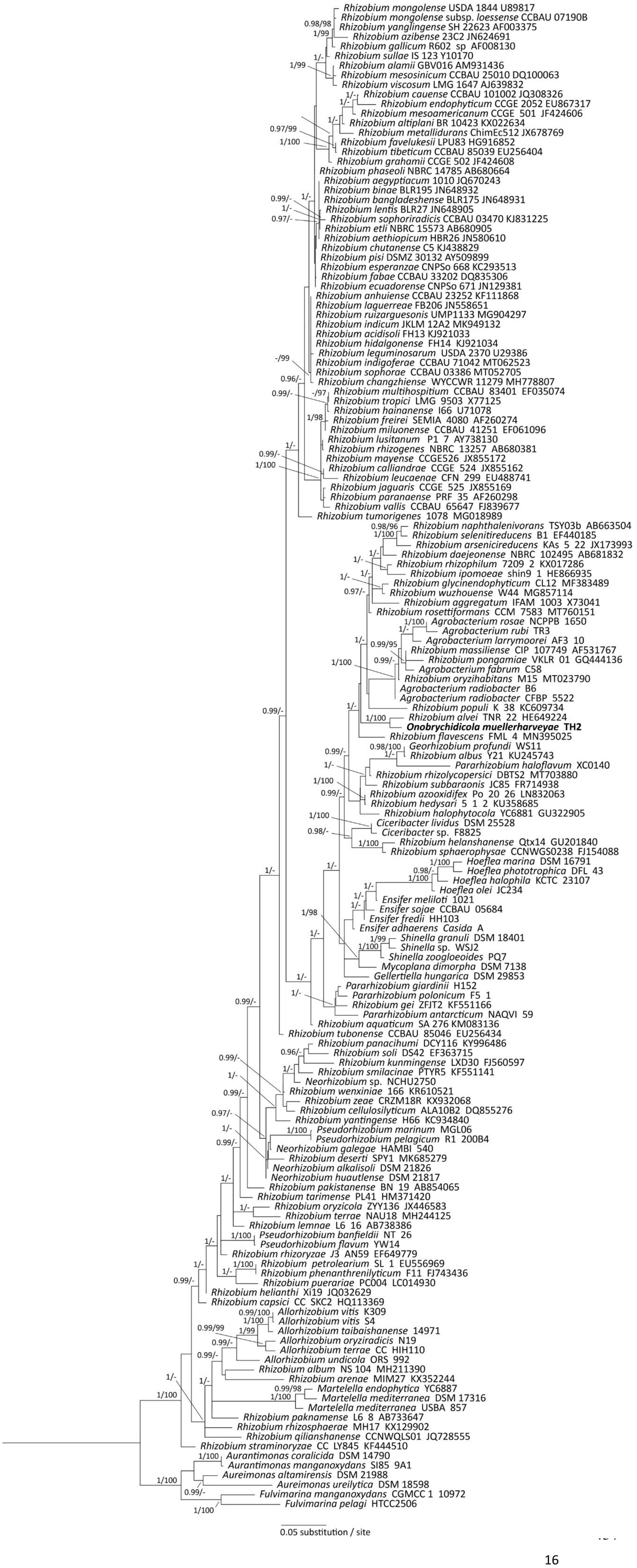


**Figures S3 –** A maximum likelihood topology estimated by the IQ-TREE analysis of 16S rRNA gene partial sequences of taxa of *Rhizobiaceae* using TIM3+F+I+G4 as best-fit substitution model. Alignment contains 157 sequences with 1405 nucleotide sites, 436 distinct patterns, 254 parsimony-informative, 105 singleton sites, and 1050 constant sites. Support values of approximate Bayesian posterior probabilities and ultrafast bootstrap ≥95 % are given above each node, respectively. The new species is highlighted in bold. The tree is rooted by representatives of A*urantimonas*, *Aureimonas* and *Fulvimarina*.

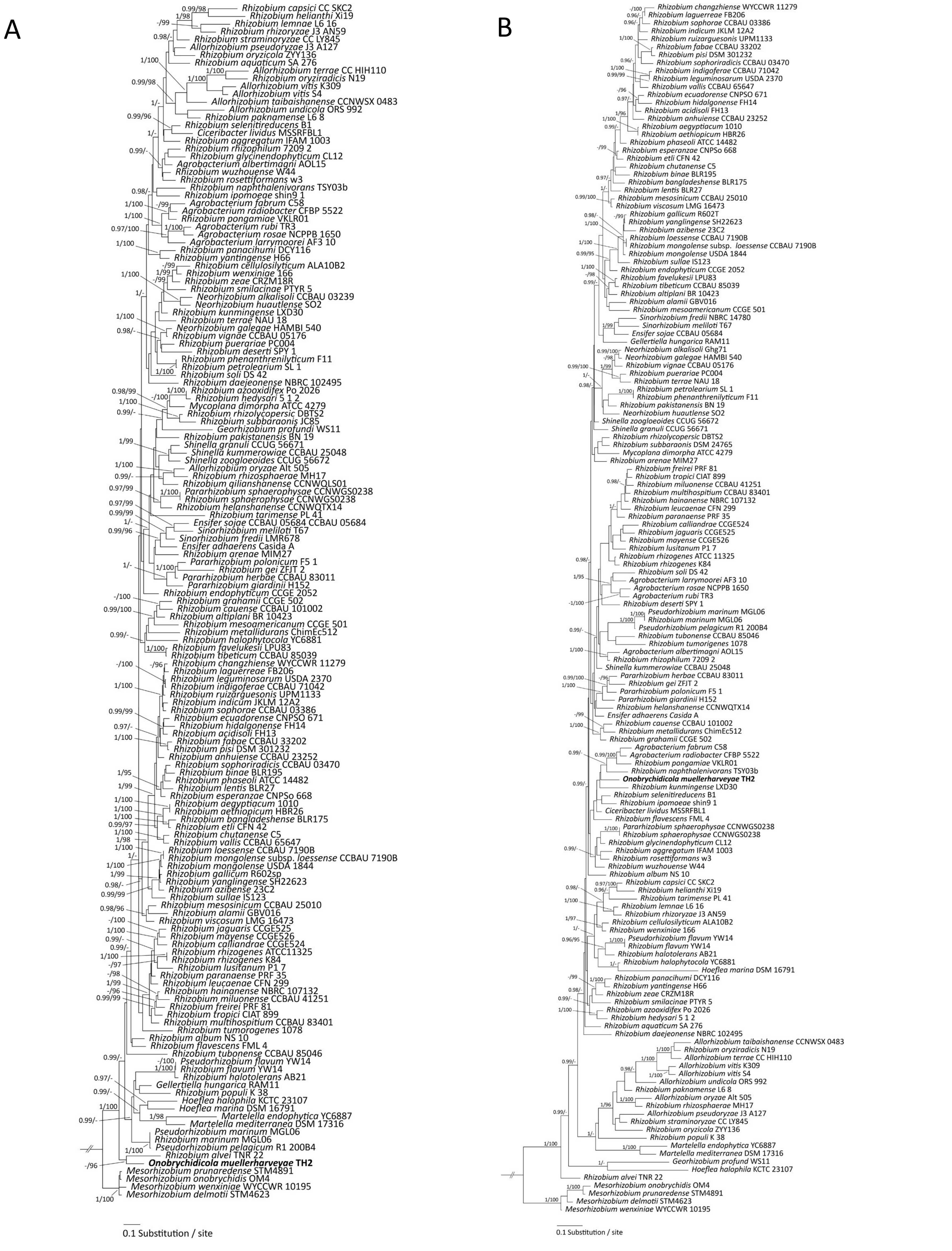


**Figures S4 –** *recA* and *atpD Onobrychidicola muellerharveyae* TH2$^{T}$. **A.** A maximum likelihood topology estimated by the IQ-TREE analysis of *recA* gene partial sequences of taxa of *Rhizobiaceae* using TIM2+F+I+G4 as best-fit substitution model. Alignment contains 153 sequences with 569 nucleotide sites, 393 distinct patterns, 261 parsimony-informative, 39 singleton sites, and 269 constant sites. **B.** A maximum likelihood topology estimated by the IQ-TREE analysis of *atpD* gene partial sequences of taxa of *Rhizobiaceae* using GTR+F+I+G4 as best-fit substitution model. Alignment contains 152 sequences with 512 nucleotide sites, 310 distinct patterns, 227 parsimony-informative, 26 singleton sites, and 259 constant sites. Support values of approximate Bayesian posterior probabilities and ultrafast bootstrap ≥95 % are given above each node, respectively. The new species is highlighted in bold. The tree is rooted by representatives of *Mesorhizobium*

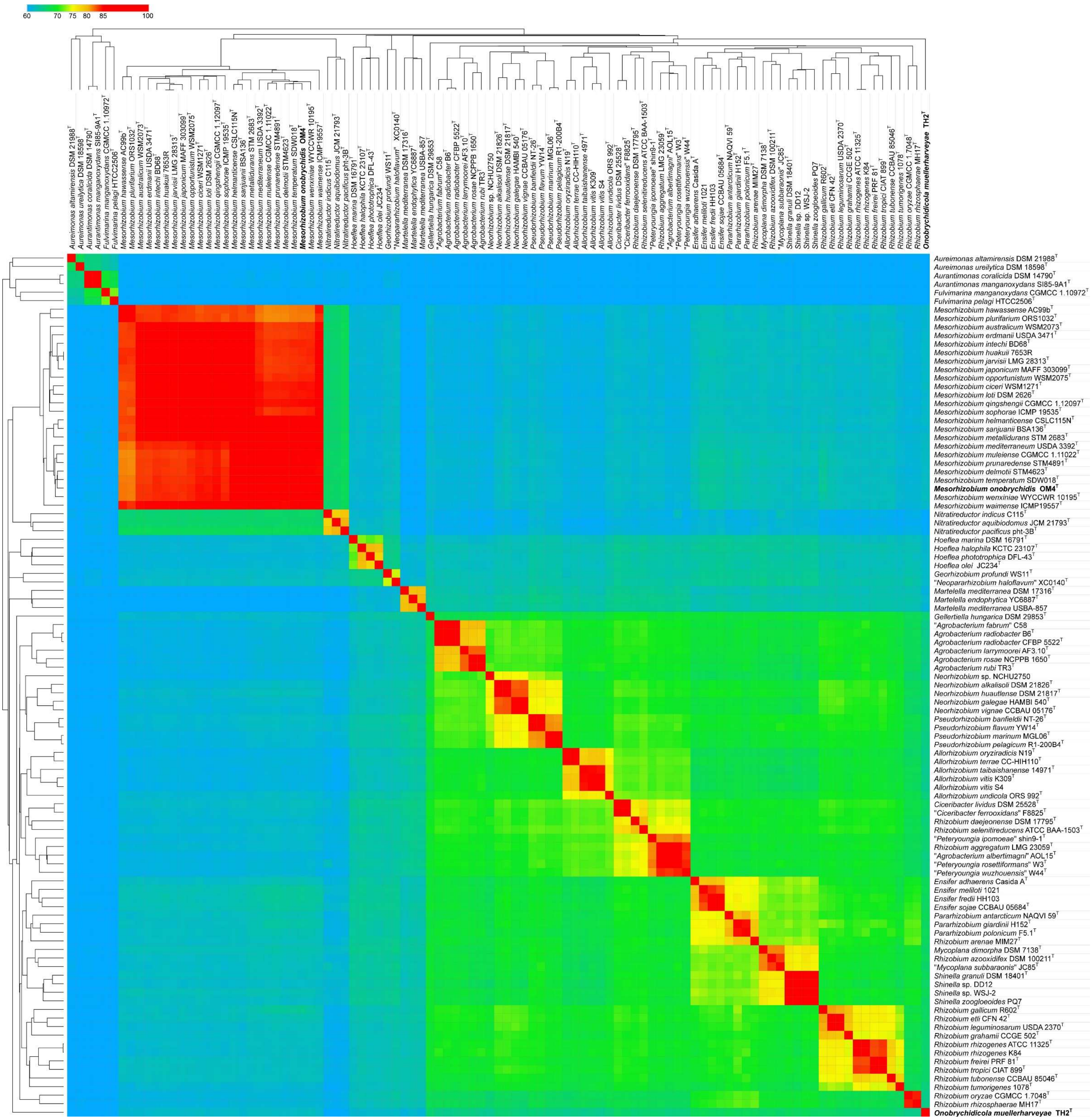


**Figure S5.** Dendrogram and heatmap based on average amino acid identity (AAI) metrics displaying genomic relatedness between strains TH2$^{T}$ and *Mesorhizobium onobrychidis* strain OM4$^{T}$, and other members of the families *Rhizobiaceae* (various genera) and *Phyllobacteriaceae* (genera *Mesorhizobium* and *Nitratireductor*). The representatives of genera *Aurantimonas*, *Aureimonas* and *Fulvimarina* were included as outgroups. The average amino acid identity (AAI) between pairs of genomes was calculated with the CompareM software. Morpheus software was used to generate heat map and for hierarchical clustering. Hierarchical clustering was performed based on precomputed AAI similarity values. The heatmap shows level of similarity on the scale of 60 to 100%. The respective thermal color scale is located at the top left corner of the figure.

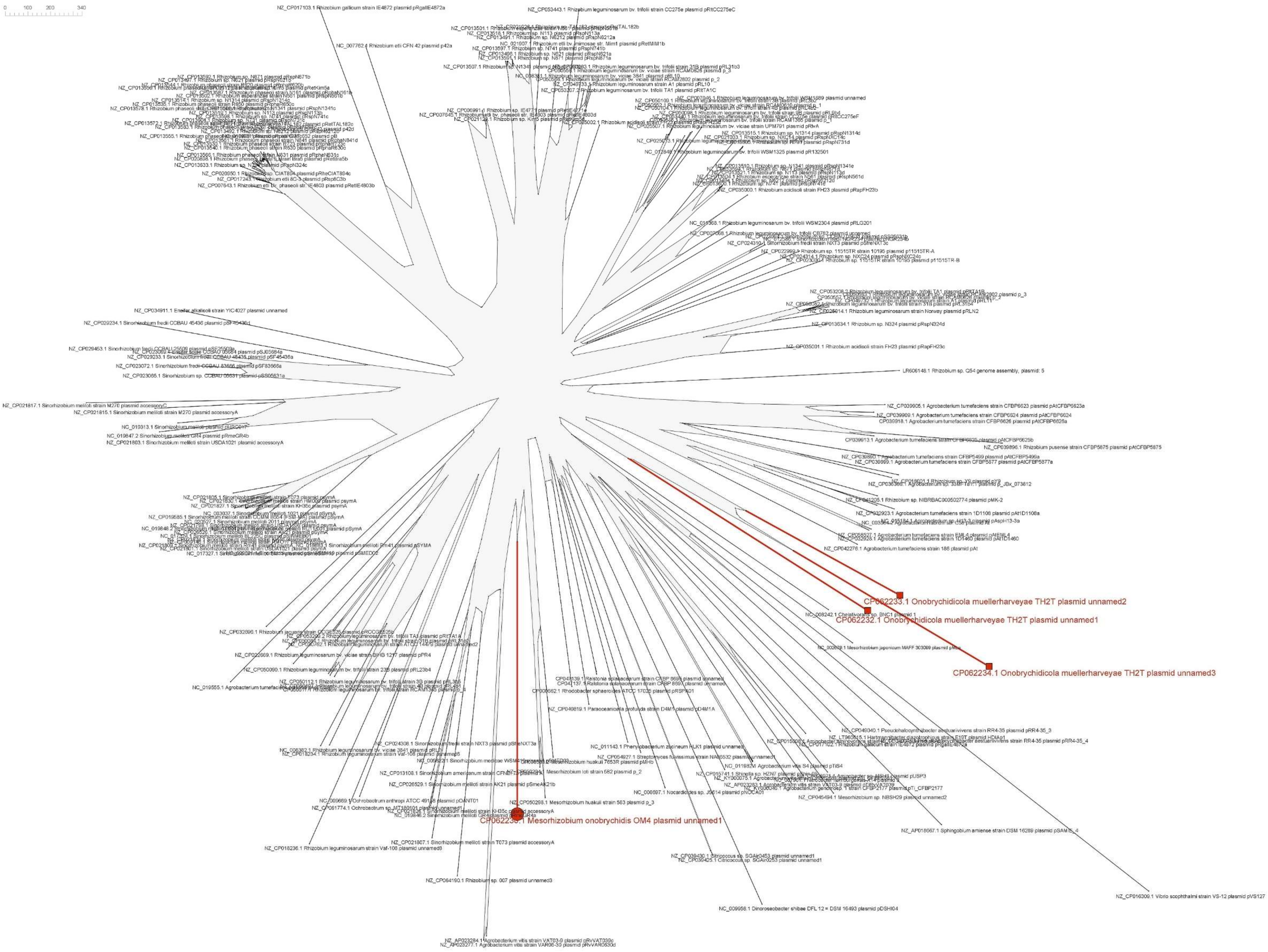


**Figure S6: Outline Neighbor Network for plasmids of strain *Mesorhizobium onobrychidis* OM4$^T$ and O*nobrychidicola muellerharveyae* TH2$^T$ based on mash distances**. The network was calculated on the mash distances with Splitstree5 for all 204 plasmid sequences of the Refseq and PLS database that show at least 1 overlapping hash to the plasmids of strain *M. onobrychidis* OM4$^T$ and *O. muellerharveyae* TH2$^T$, that are marked in yellow. The final tree was achieved with the outline algorithm [37].

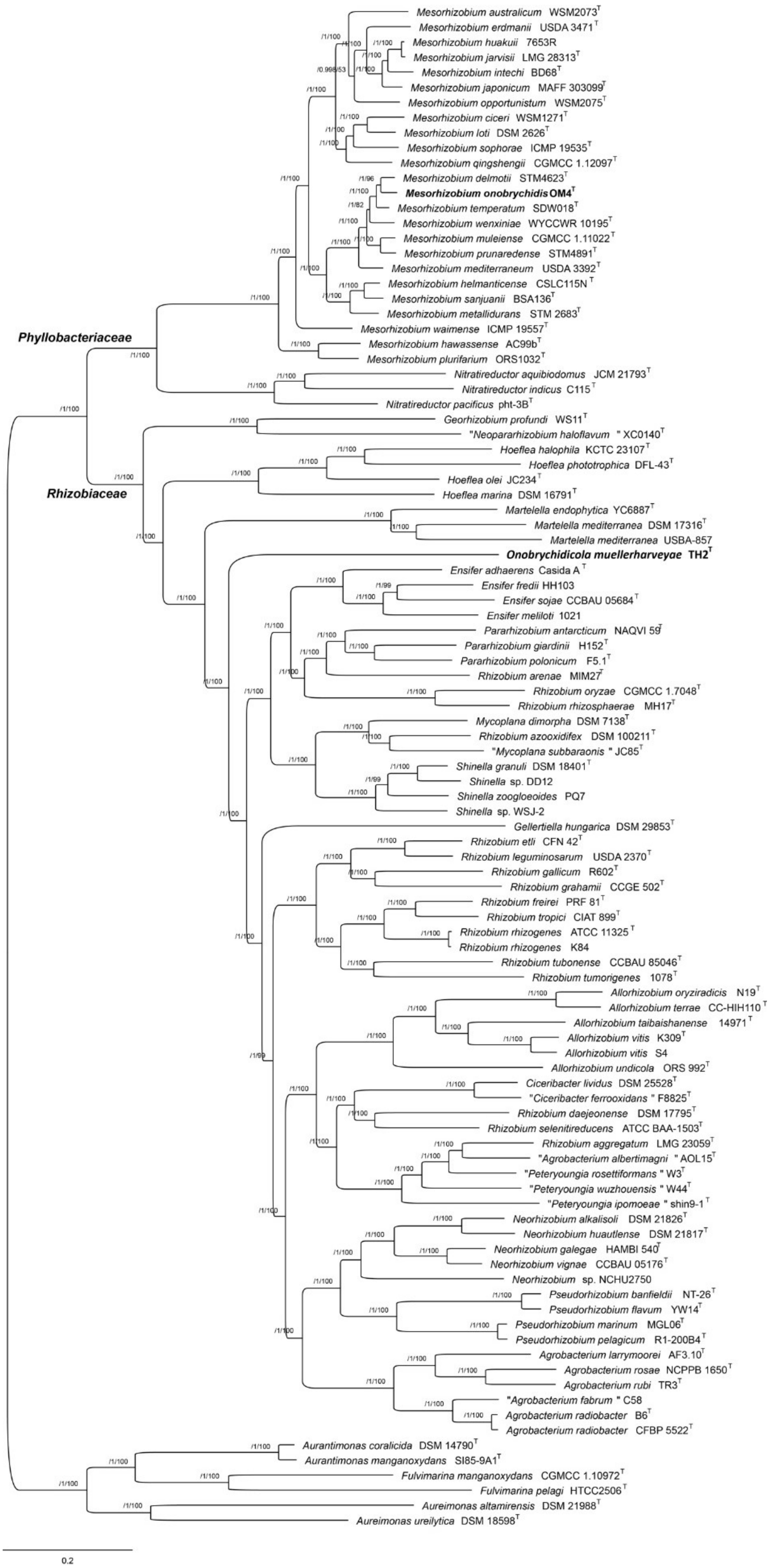


**Figure S7 -** Maximum-likelihood core-genome phylogeny of strains *Onobrychidicola muellerharveyae* TH2$^{T}$ and *Mesorhizobium onobrychidis* OM4$^{T}$, including representatives of families *Rhizobiaceae* (various genera) and *Phyllobacteriaceae* (genera *Mesorhizobium* and *Nitratireductor*). The tree was estimated with IQ-TREE from the concatenated alignment of 118 top-ranked genes selected using GET_PHYLOMARKERS software. The numbers on the nodes indicate the approximate Bayesian posterior probabilities support values (first value) and ultra-fast bootstrap values (second value), as implemented in IQ-TREE. The tree was rooted using the sequences of representatives of genera *Aurantimonas*, *Aureimonas* and *Fulvimarina* as outgroups. The scale bar represents the number of expected substitutions per site under the best-fitting GTR+F+ASC+R8 model.

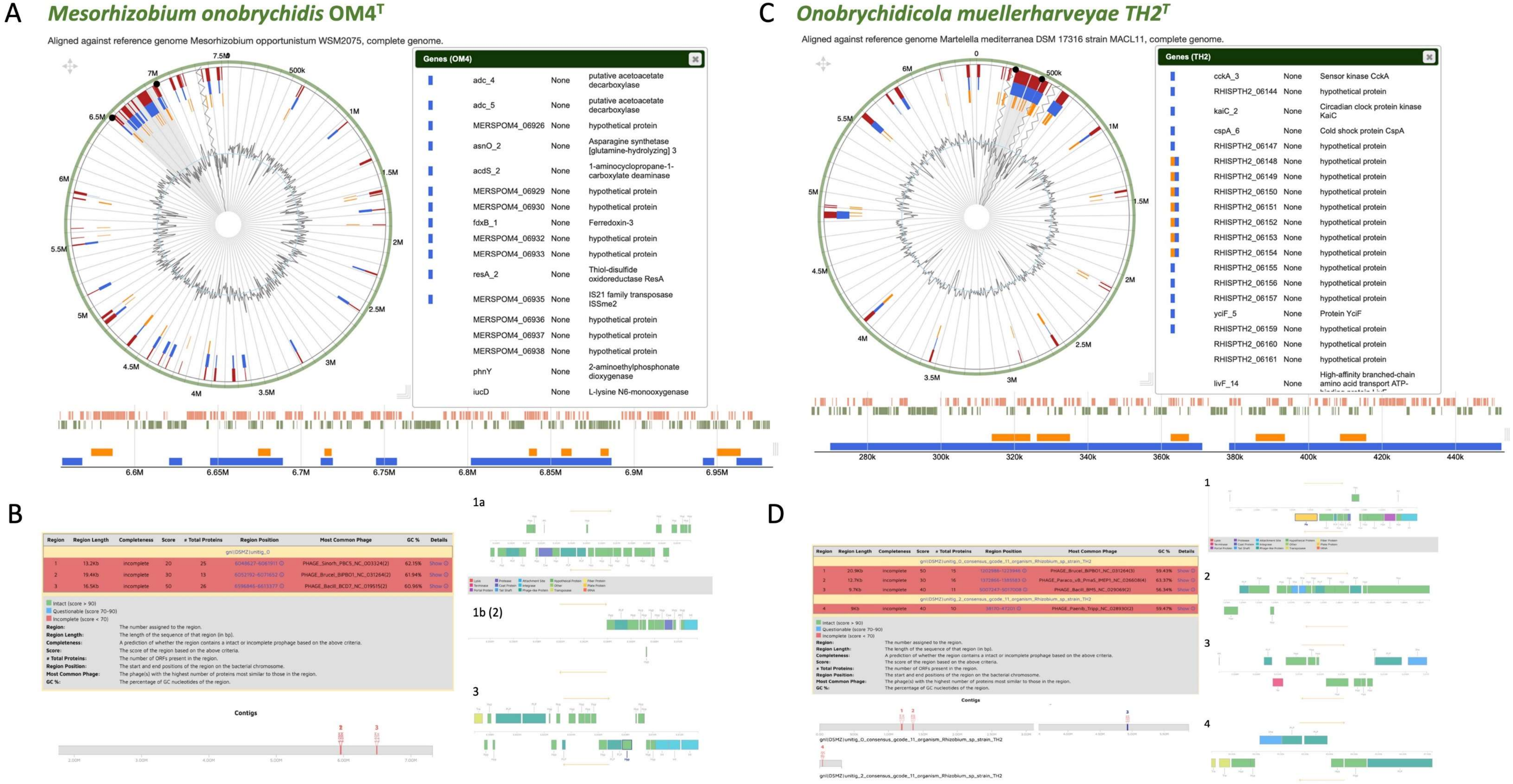


**Figure S8.** Genomic islands (A,C) and phage annotions (B,D) of *Mesorhizobium onobrychidis* strain OM4$^T$ and *Onobrychidicola muellerharveyae* TH2$^T$ based on the island viewer and phaster web tool.

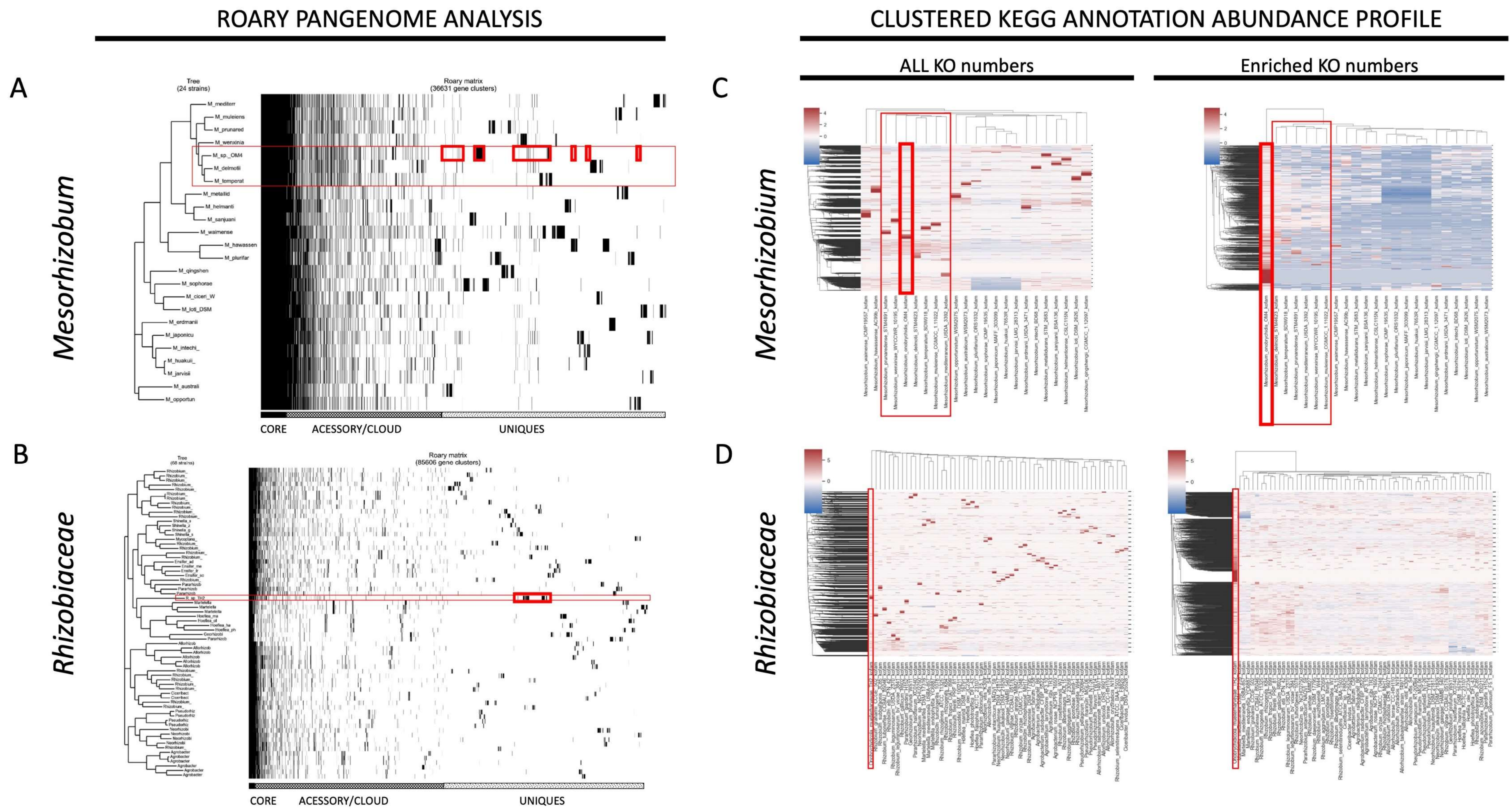


**Figure S9**. Pangenome analysis and KEGG abundance clustering confirm species and genus novelty of both strains. Results are presented for the *Mesorhizobia* (A,C) and *Rhizobioceae* clade (B,D). Figures A and D give a brief overview of core, accessory, cloud and unique genes for all strains ordered phylogenetic distance based on core gene SNPs. Figures C and D show the clusters considering all KO identifiers on the left or on the right only the *Mesorhizobium onobrychidis* strain OM4$^{T}$ or TH2$^{T}$ enriched ones compared to all strains of the respective clade. Here the strains are listed on the x-axis. Red bars indicate enrichments while blue mark depletions.

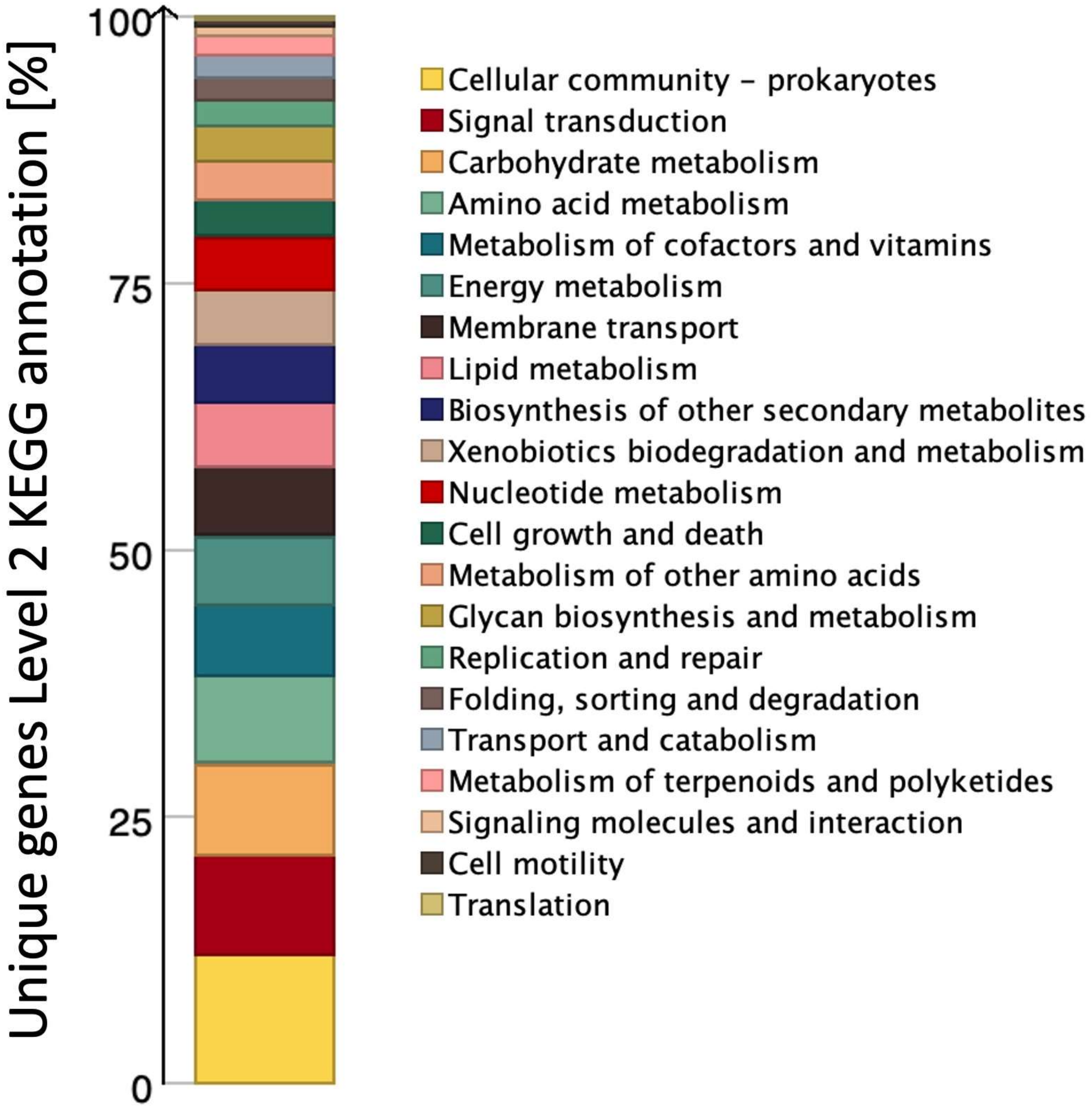


**Figure S10**. Functional annotation of unique genes detected for *Mesorhizobium onobrychidis* strain OM4$^{T}$ show enriched level 2 KEGG classes in a percentage scale.

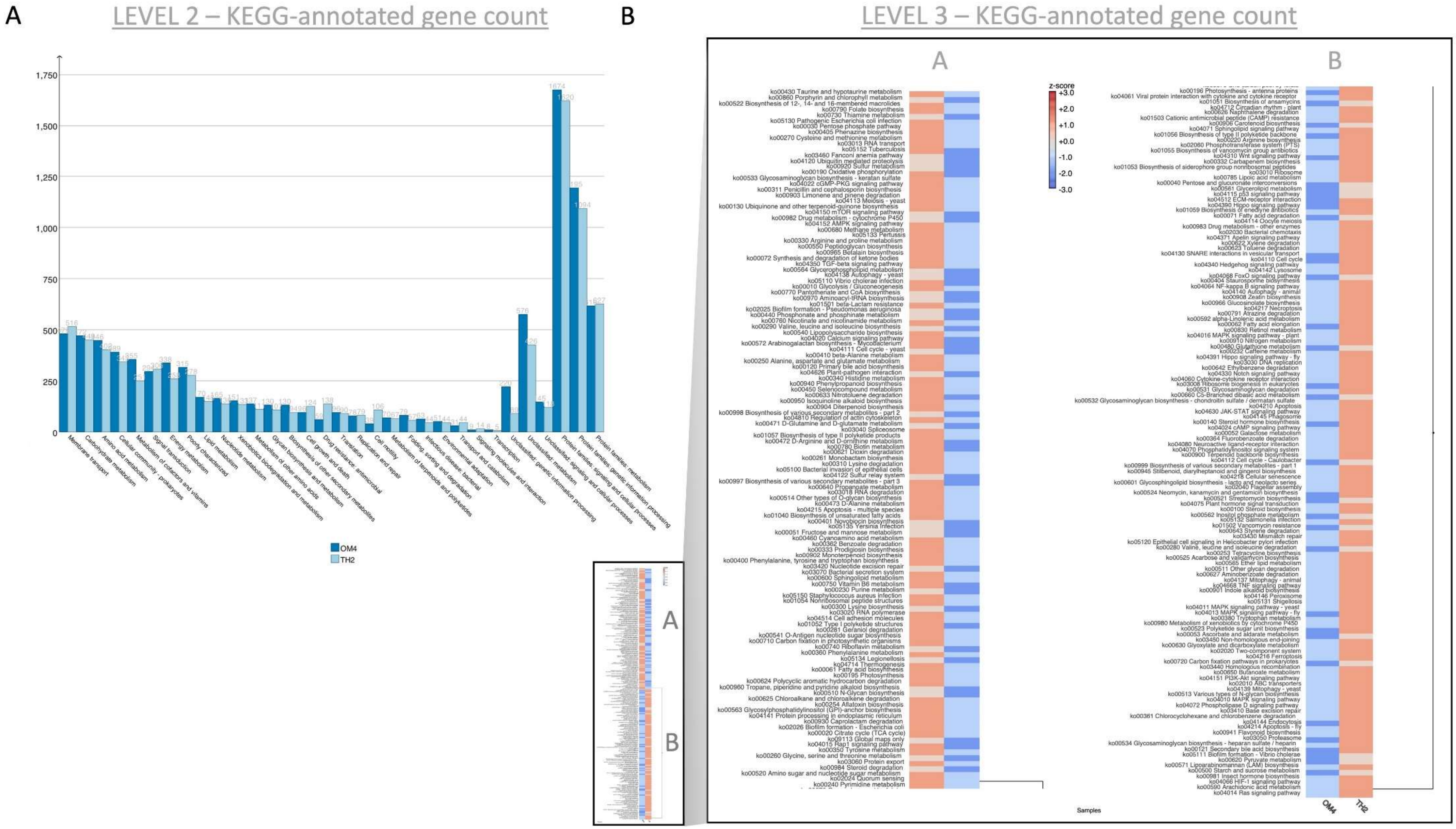


**Figure S11**. KEGG annotations of strains *Mesorhizobium onobrychidis* OM4$^{T}$ and TH2$^{T}$ on level 2 (A) show distinct class counts and on level 3 (B) specific abundance pattern as clustered heatmap of calculated Z-scores. While Cluster A is *Mesorhizobium onobrychidis* strain OM4$^{T}$ -specific cluster B is strongly associated with TH2$^{T}$.

A

### *Mesorhizobium onobrychidis* OM4$^T$

#### CHROMOSOME

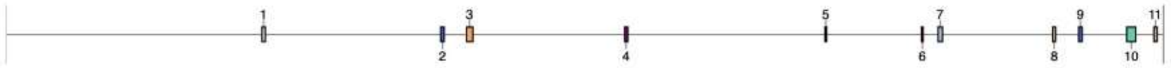

| Region | Type | From | To | Most similar known cluster | | Similarity |
|---|---|---|---|---|---|---|
| Region 1.1 | indole | 1,617,818 | 1,639,032 | | | |
| Region 1.2 | redox-cofactor | 2,748,569 | 2,770,738 | | | |
| Region 1.3 | T3PKS | 2,910,833 | 2,951,900 | surfactin | NRP:Lipopeptide | 8% |
| Region 1.4 | terpene | 3,911,865 | 3,932,692 | | | |
| Region 1.5 | RiPP-like | 5,179,457 | 5,190,374 | | | |
| Region 1.6 | siderophore | 5,790,051 | 5,801,316 | | | |
| Region 1.7 | RRE-containing, lassopeptide | 5,895,466 | 5,922,923 | | | |
| Region 1.8 | hserlactone | 6,620,762 | 6,639,995 | | | |
| Region 1.9 | proteusin | 6,784,936 | 6,806,435 | | | |
| Region 1.10 | phosphonate | 7,089,157 | 7,147,223 | 9-methylstreptimidone | Polyketide:Modular type I | 12% |
| Region 1.11 | hserlactone | 7,260,642 | 7,281,280 | | | |

B

### *Onobrychidicola muellerharveyae* TH2$^T$

#### CHROMOSOME

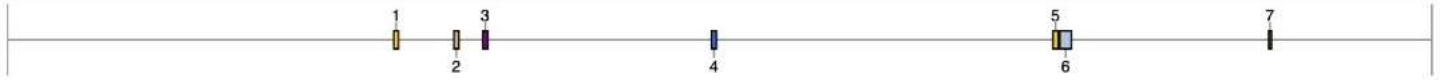

| Region | Type | From | To | Most similar known cluster | | Similarity |
|---|---|---|---|---|---|---|
| Region 1.1 | thioamitides | 1,595,037 | 1,614,575 | | | |
| Region 1.2 | hserlactone | 1,842,937 | 1,863,680 | | | |
| Region 1.3 | terpene | 1,962,657 | 1,983,514 | | | |
| Region 1.4 | RRE-containing | 2,906,412 | 2,926,693 | galactoglucan | Saccharide | 16% |
| Region 1.5 | thioamitides | 4,315,628 | 4,337,384 | | | |
| Region 1.6 | NRPS, T1PKS | 4,344,136 | 4,392,211 | | | |
| Region 1.7 | ectoine | 5,206,699 | 5,217,085 | | | |

#### PLASMID 1

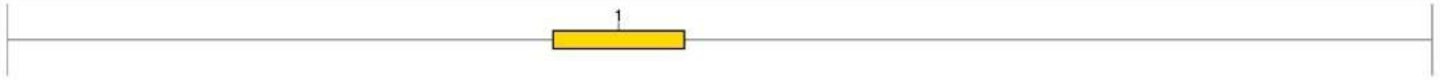

| Region | Type | From | To | Most similar known cluster | Similarity |
|---|---|---|---|---|---|
| Region 2.1 | thioamitides | 91,288 | 113,261 | | |

**Figure S12.** AntiSMASH analyses of strains *Mesorhizobium onobrychidis* OM4$^T$ (A) and TH2$^T$ (B) reveal specific secondary metabolite biosynthetic gene clusters, their genomic location and confirm strain specific lifestyles.

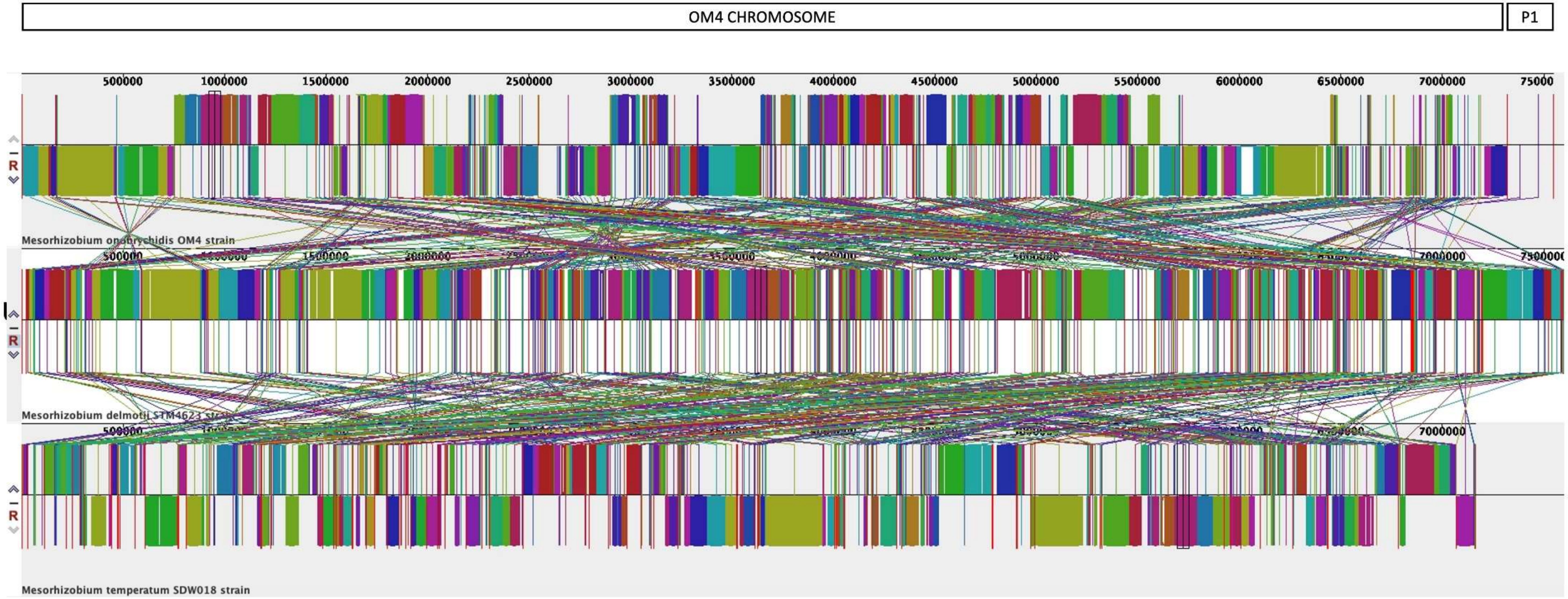


- Unaligned regions of OM4 to both other strains, that are not part of a locally collinear block (LCB) of minimal weight 52:
  - Total: 486 regions
  - Length = 2000 bp: 135 regions (max. length 91,884 bp)
  - Gene Count >= 5: 85 regions (max. gene count 77)
  - antiSMASH Clusters: 21 regions (covering 7 of 11 BGCs)
  - Symbiotic Island: 63 regions (covering 2 of 7 BGCs)
  - Unique Gene hits: 628 genes (of 1068 in total)

**Figure S13.** Whole genome MAUVE alignment of *Mesorhizobium onobrychidis* strain OM4$^T$ and its close relative *M. delmotii* STM4623T and *M. temperatum* SDW018T uncovers conserved locally collinear blocks of minimal weight 52 and unconserved unaligned unique regions per strain (Table S1).

**M. onobrychidis OM4$^T$**

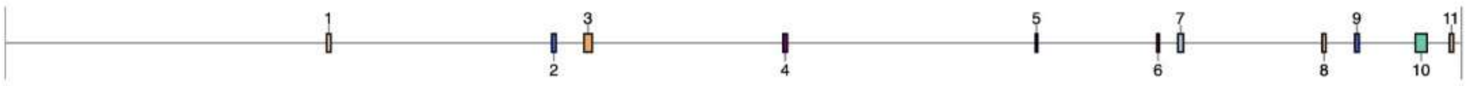


| Region | Type | From | To | | Most similar known cluster | | Similarity |
|---|---|---|---|---|---|---|---|
| Region 1.1 | indole | 1,617,818 | 1,639,032 | U | | | |
| Region 1.2 | redox-cofactor | 2,748,569 | 2,770,738 | | | | |
| Region 1.3 | T3PKS | 2,910,833 | 2,951,900 | | surfactin | NRP:Lipopeptide | 8% |
| Region 1.4 | terpene | 3,911,865 | 3,932,692 | | | | |
| Region 1.5 | RiPP-like | 5,179,457 | 5,190,374 | U | | | |
| Region 1.6 | siderophore | 5,790,051 | 5,801,316 | U | | | |
| Region 1.7 | RRE-containing, lassopeptide | 5,895,466 | 5,922,923 | U | | | |
| Region 1.8 | hserlactone | 6,620,762 | 6,639,995 | U | | | |
| Region 1.9 | proteusin | 6,784,936 | 6,806,435 | U | | | |
| Region 1.10 | phosphonate | 7,089,157 | 7,147,223 | U | 9-methylstreptimidone | Polyketide:Modular type I | 12% |
| Region 1.11 | hserlactone | 7,260,642 | 7,281,280 | | | | |

**M. delmotii STM4623$^T$**

| Region | Type | From | To | Most similar known cluster | | Similarity |
|---|---|---|---|---|---|---|
| Region 3.1 | hserlactone | 812,108 | 832,746 | | | |
| Region 3.2 | RiPP-like | 894,270 | 905,301 | | | |
| Region 3.3 | RRE-containing | 1,553,434 | 1,573,718 | | | |
| Region 5.1 | phosphonate, hserlactone | 100,180 | 157,149 | | | |
| Region 7.1 | RRE-containing | 31,833 | 53,008 | | | |
| Region 8.1 | NRPS-like | 3,623 | 46,460 | pyoverdin | NRP | 2% |
| Region 9.1 | T3PKS | 17,818 | 58,885 | | | |
| Region 10.1 | redox-cofactor | 156,194 | 178,363 | | | |
| Region 15.1 | hserlactone | 120,228 | 140,857 | | | |
| Region 22.1 | phosphonate | 11,092 | 52,051 | chejuenolide A / chejuenolide B | Polyketide | 7% |
| Region 24.1 | NRPS-like, T1PKS | 1 | 28,938 | | | |
| Region 27.1 | RiPP-like | 123 | 11,052 | | | |
| Region 28.1 | RiPP-like | 168,193 | 178,897 | | | |
| Region 32.1 | terpene | 231,363 | 252,214 | | | |

**M. temperatum SDW018$^T$**

| Region | Type | From | To | Most similar known cluster | | Similarity |
|---|---|---|---|---|---|---|
| Region 7.1 | siderophore | 103,419 | 116,881 | | | |
| Region 14.1 | NRPS-like | 1 | 32,901 | corpeptin A / corpeptin B | NRP | 18% |
| Region 19.1 | RiPP-like | 125,640 | 136,671 | | | |
| Region 31.1 | NRPS-like | 1,354 | 32,370 | succinoglycan | Saccharide | 33% |
| Region 48.1 | hserlactone | 81,818 | 102,456 | | | |
| Region 50.1 | terpene | 32,734 | 53,561 | | | |
| Region 57.1 | phosphonate | 43,665 | 89,806 | colicin V | RiPP | 2% |
| Region 69.1 | hserlactone | 1,523 | 22,155 | | | |
| Region 72.1 | proteusin | 5,034 | 26,518 | | | |
| Region 72.2 | hserlactone | 63,672 | 84,304 | | | |
| Region 75.1 | redox-cofactor | 57,177 | 79,346 | | | |
| Region 75.2 | T3PKS | 244,823 | 285,890 | | | |
| Region 76.1 | RiPP-like | 28,412 | 39,329 | | | |

**Figure S14.** AntiSMASH analysis of secondary metabolite biosynthesis gene cluster of *Mesorhizobium onobrychidis* OM4$^T$ and its close relative *M. delmotii* STM4623T and *M. temperatum* SDW018T points to strain specific encoding regions and BGC types. For *Mesorhizobium onobrychidis* OM4$^T$ seven biosynthesis gene clusters out of 11 are marked with an U specifying their association to an unaligned regions that are not part of a collinear block among all three strains.

**Literature**


[1] Weisburg WG, Barns SM, Pelletier DA, Lane DJ. 16S ribosomal DNA amplification for phylogenetic study. J Bacteriol 1991; 173(2): 697–703 [https://doi.org/10.1128/jb.173.2.697-703.1991][PMID: 1987160]

[2] Heuer H, Wieland G, Schönfeld J, Schönwälder A, Gomes N, Smalla K. Bacterial community profiling using DGGE or TGGE analysis. In: Rochelle PA, editor. Environmental molecular microbiology: Protocols and applications. Wymondham: Horizon Scientific Press 2001; 177–90.

[3] Gaunt MW, Turner SL, Rigottier-Gois L, Lloyd-Macgilp SA, Young JP. Phylogenies of atpD and recA support the small subunit rRNA-based classification of rhizobia. Int J Syst Evol Microbiol 2001; 51(Pt 6): 2037–48 [https://doi.org/10.1099/00207713-51-6-2037][PMID: 11760945]

[4] Kuzmanović N, Smalla K, Gronow S, Puławska J. *Rhizobium tumorigenes* sp. nov., a novel plant tumorigenic bacterium isolated from cane gall tumors on thornless blackberry. Sci Rep 2018; 8(1): 9051 [https://doi.org/10.1038/s41598-018-27485-z][PMID: 29899540]

[5] Altschul SF, Gish W, Miller W, Myers EW, Lipman DJ. Basic local alignment search tool. Journal of Molecular Biology 1990; 215(3): 403–10 [https://doi.org/10.1016/S0022-2836(05)80360-2]

[6] Li H, Durbin R. Fast and accurate short read alignment with Burrows-Wheeler transform. Bioinformatics 2009; 25(14): 1754–60 [https://doi.org/10.1093/bioinformatics/btp324][PMID: 19451168]

[7] Langmead B, Salzberg SL. Fast gapped-read alignment with Bowtie 2. Nat Methods 2012; 9(4): 357–9 [https://doi.org/10.1038/nmeth.1923][PMID: 22388286]

[8] Koboldt DC, Zhang Q, Larson DE*, et al.* VarScan 2: somatic mutation and copy number alteration discovery in cancer by exome sequencing. Genome Res 2012; 22(3): 568–76 [https://doi.org/10.1101/gr.129684.111][PMID: 22300766]

[9] Seemann T. Prokka: rapid prokaryotic genome annotation. Bioinformatics 2014; 30(14): 2068–9 [https://doi.org/10.1093/bioinformatics/btu153][PMID: 24642063]

[10] Tatusova T, DiCuccio M, Badretdin A*, et al.* NCBI prokaryotic genome annotation pipeline. Nucleic Acids Res 2016; 44(14): 6614–24 [https://doi.org/10.1093/nar/gkw569][PMID: 27342282]

[11] Katoh K, Rozewicki J, Yamada KD. MAFFT online service: multiple sequence alignment, interactive sequence choice and visualization. Brief Bioinformatics 2019; 20(4): 1160–6 [https://doi.org/10.1093/bib/bbx108][PMID: 28968734]

[12] Larsson A. AliView: a fast and lightweight alignment viewer and editor for large datasets. Bioinformatics 2014; 30(22): 3276–8 [https://doi.org/10.1093/bioinformatics/btu531][PMID: 25095880]

[13] Minh BQ, Schmidt HA, Chernomor O*, et al.* IQ-TREE 2: New Models and Efficient Methods for Phylogenetic Inference in the Genomic Era. Mol Biol Evol 2020; 37(5): 1530–4 [https://doi.org/10.1093/molbev/msaa015][PMID: 32011700]

[14] Hoang DT, Chernomor O, Haeseler A von, Minh BQ, Le Vinh S. UFBoot2: Improving the Ultrafast Bootstrap Approximation. Mol Biol Evol 2018; 35(2): 518–22 [https://doi.org/10.1093/molbev/msx281][PMID: 29077904]

[15] Ronquist F, Huelsenbeck JP. MrBayes 3: Bayesian phylogenetic inference under mixed models. Bioinformatics 2003; 19(12): 1572–4 [https://doi.org/10.1093/bioinformatics/btg180][PMID: 12912839]

[16] Nylander JA. MrModeltest v2. Program distributed by the author 2004.

[17] Contreras-Moreira B, Vinuesa P. GET_HOMOLOGUES, a versatile software package for scalable and robust microbial pangenome analysis. Appl Environ Microbiol 2013; 79(24): 7696–701 [https://doi.org/10.1128/AEM.02411-13][PMID: 24096415]

[18] Vinuesa P, Ochoa-Sánchez LE, Contreras-Moreira B. GET_PHYLOMARKERS, a software package to select optimal orthologous clusters for phylogenomics and inferring pan-genome phylogenies, used for a critical geno-taxonomic revision of the genus *Stenotrophomonas*. Front Microbiol 2018; 9: 771 [https://doi.org/10.3389/fmicb.2018.00771][PMID: 29765358]

[19] Nguyen L-T, Schmidt HA, Haeseler A von, Minh BQ. IQ-TREE: a fast and effective stochastic algorithm for estimating maximum-likelihood phylogenies. Mol Biol Evol 2015; 32(1): 268–74 [https://doi.org/10.1093/molbev/msu300][PMID: 25371430]

[20] Goris J, Konstantinidis KT, Klappenbach JA, Coenye T, Vandamme P, Tiedje JM. DNA-DNA hybridization values and their relationship to whole-genome sequence similarities. Int J Syst Evol Microbiol 2007; 57(Pt 1): 81–91 [https://doi.org/10.1099/ijs.0.64483-0][PMID: 17220447]

[21] Konstantinidis KT, Tiedje JM. Towards a genome-based taxonomy for prokaryotes. J Bacteriol 2005; 187(18): 6258–64 [https://doi.org/10.1128/JB.187.18.6258-6264.2005][PMID: 16159757]

[22] Konstantinidis KT, Rosselló-Móra R, Amann R. Uncultivated microbes in need of their own taxonomy. ISME J 2017; 11(11): 2399–406 [https://doi.org/10.1038/ismej.2017.113][PMID: 28731467]

[23] Kuzmanović N, Fagorzi C, Mengoni A, Lassalle F, diCenzo GC. Taxonomy of Rhizobiaceae revisited: proposal of a new framework for genus delimitation 2021.

[24] Richter M, Rosselló-Móra R. Shifting the genomic gold standard for the prokaryotic species definition. Proc Natl Acad Sci U S A 2009; 106(45): 19126–31 [https://doi.org/10.1073/pnas.0906412106][PMID: 19855009]

[25] Yoon S-H, Ha S-M, Lim J, Kwon S, Chun J. A large-scale evaluation of algorithms to calculate average nucleotide identity. Antonie Van Leeuwenhoek 2017; 110(10): 1281–6 [https://doi.org/10.1007/s10482-017-0844-4][PMID: 28204908]

[26] Meier-Kolthoff JP, Auch AF, Klenk H-P, Göker M. Genome sequence-based species delimitation with confidence intervals and improved distance functions. BMC Bioinformatics 2013; 14: 60 [https://doi.org/10.1186/1471-2105-14-60][PMID: 23432962]

[27] Meier-Kolthoff JP, Göker M. TYGS is an automated high-throughput platform for state-of-the-art genome-based taxonomy. Nat Commun 2019; 10(1): 2182 [https://doi.org/10.1038/s41467-019-10210-3][PMID: 31097708]

[28] Heimbrook ME, Wang WL, Campbell G. Staining bacterial flagella easily. J Clin Microbiol 1989; 27(11): 2612–5 [https://doi.org/10.1128/jcm.27.11.2612-2615.1989][PMID: 2478573]

[29] Smibert RM, Krieg NR. Phenotypic characterization. In: Kamlage B, editor. Methods for General and Molecular Bacteriology. Edited by P. Gerhardt, R. G. E. Murray, W. A. Wood and N. R. Krieg. Washington, D.C. 1994; 607–54.

[30] Bouzar H, Jones JB, Bishop AL. Simple cultural tests for identification of *Agrobacterium* biovars. In: Gartland KMA, Davey MR, editors. Agrobacterium Protocols. Totowa, NJ: Springer New York 1995; 9–13.

[31] Ryu E. On the gram-differentiation of bacteria by the simplest method. II. The caustic potash method. The Japanese Journal of Veterinary Science 1939; 1(2): 204–10 [https://doi.org/10.1292/jvms1939.1.204]

[32] Cerny G. Method for the distinction of gramnegative from grampositive bacteria. European J. Appl Microbiol. 1976; 3(3): 223–5 [https://doi.org/10.1007/BF01385437]
[33] Kovacs N. Identification of *Pseudomonas pyocyanea* by the oxidase reaction. Nature 1956; 178(4535): 703 [https://doi.org/10.1038/178703a0][PMID: 13369512]
[34] Sasser M. Identification of bacteria by gas chromatography of cellular fatty acids. Technical Note 1990; 101, DE, MID.
[35] Vieira S, Huber KJ, Neumann-Schaal M*, et al.* Usitatibacter rugosus gen. nov., sp. nov. and Usitatibacter palustris sp. nov., novel members of Usitatibacteraceae fam. nov. within the order Nitrosomonadales isolated from soil. Int J Syst Evol Microbiol 2021; 71(2) [https://doi.org/10.1099/ijsem.0.004631][PMID: 33433313]
[36] Broughton WJ, Dilworth MJ. Control of leghaemoglobin synthesis in snake beans. Biochem J 1971; 125(4): 1075–80 [https://doi.org/10.1042/bj1251075][PMID: 5144223]
[37] Bagci C, Bryant D, Cetinkaya B, Huson DH. Microbial Phylogenetic Context Using Phylogenetic Outlines. Genome Biol Evol 2021; 13(9) [https://doi.org/10.1093/gbe/evab213][PMID: 34519776]
